# A Bioinformatics-Driven ceRNA Network in Stomach Adenocarcinoma: Identification of Novel Prognostic mRNA-miRNA-lncRNA Interactions

**DOI:** 10.1101/2024.12.28.628529

**Authors:** Ebtihal Kamal, Zainab Mohammed Mahmoud Omar, Ayman Geddawy, Ahmad AA Omer

## Abstract

Stomach adenocarcinoma is a major contributor to worldwide mortality and poses a substantial challenge to improving life expectancy. The main objective of the current work was to identify diagnostic and prognostic biomarkers in stomach adenocarcinoma in order to advance translational medicine and patient outcomes. By accomplishing this objective, the research seeks to provide significant insights into the field of translational medicine. Seven novel unfavourable prognosis-associated genes (NALCN, CALCR, CPT1C, ELAVL3, FLJ16779, MYOZ3, and TPST1) were first identified. Additionally, 41 potential miRNAs were predicted. ELAVL3-hsa-mir-29a-3p and CALCR-hsa-mir-29a-3p axes were identified as two critical pathways in the carcinogenesis of stomach adenocarcinoma via a bioinformatics analysis. Following that, lncRNAs binding to hsa-mir-29a-3p were predicted via starBase and miRNet databases for predicting lncRNA binding sites. After conducting both expression and survival analyses for these predicted lncRNAs, we found that only one lncRNA (KCNQ1OT1) was markedly overexpressed in stomach adenocarcinoma, and its elevated expression was associated with an unfavourable prognosis. Next, we established a comprehensive mRNA-miRNA-lncRNA triple ceRNA network linked to the prognosis of patients with stomach adenocarcinoma. In summary, the current study provides an extensive ceRNA network that highlights novel diagnostic and prognostic biomarkers for stomach adenocarcinoma.

## Introduction

Despite advances in treatment approaches, gastric cancer (GC), particularly stomach adenocarcinoma (STAD), is a major global health concern, with a high mortality rate and a poor prognosis, ranking it as the fifth most common cancer and a leading cause of cancer-related mortality worldwide, based on GLOBOCAN 2020 (1). The risk of stomach adenocarcinoma (STAD) is increased by chronic gastropathies, such as Helicobacter pylori (H. pylori) or Epstein-Barr virus (EBV) infections, as well as demographic and lifestyle variables (2,3). Additionally, genetic predisposition contributes to the development and progression of STAD with attention to long non-coding RNAs (lncRNAs) such as HOX transcript antisense RNA (HOTAIR) (4), and DNA damage-activated lncRNA (NORAD) (5). STAD treatment modalities include surgery, radiotherapy, and anticancer drugs that may be used as neoadjuvant, adjuvant, or palliative (6,7). However, current medications have little efficacy in treating advanced GC patients. New therapeutic strategies are urgently needed (8). Immunotherapy was found to be more effective than standard treatments for GC patients, resulting in longer survival times and improved outcomes (9,10). Advanced pharmacotherapy of STAD shows a rapidly evolving landscape by adding targeted/immune therapies as anti-HER2 and FGFR2 inhibitors (11). Despite advances in carcinogenesis research and new therapeutic strategies, patients with STAD have poor prognosis and there are challenges for precision medicine in gastric cancer (12,13). Identification of more molecular markers associated with STAD helps to optimally manage STAD-patients and improve their prognosis. Consequently, elucidating the intricate mechanisms of STAD pathogenesis and identifying promising diagnostic and prognostic biomarkers may facilitate the development of effective therapeutic strategies and enhance patient outcomes. Salmena et al. proposed the ceRNA hypothesis, which posits that ncRNAs, such as lncRNAs, can modulate gene expression through competitive binding to shared miRNAs with mRNAs (14). Competing endogenous RNAs (ceRNAs), including long non-coding RNAs (lncRNAs) and circular RNAs (circRNAs), can modulate the expression of target genes by competing with messenger RNAs for identical microRNA response elements (MREs) (14,15).

Our aim is to build an extensive "mRNA-miRNA-lncRNA" ceRNA network in order to find new diagnostic and prognostic biomarkers for stomach adenocarcinoma that contribute to precision medicine in STAD by utilizing cutting-edge bioinformatics technologies to identify important molecular connections connected to patient outcomes.

Recent studies have extensively investigated the roles of ceRNA regulatory networks in a variety of human malignancies, generating significant findings. ceRNA networks have been used to find prognostic markers in thyroid cancer (16), hepatocellular carcinoma (17), pancreatic cancer (18), glioblastoma (19–21), and breast cancer (22). This study adopts a unique "mRNA-miRNA-lncRNA" framework that allows for the discovery of ceRNA components with diagnostic and prognostic value in STAD. By incorporating modern analytics, it creates a unique ceRNA network, connecting molecular interactions to patient outcomes and enhancing precision medicine.

## Material and methods

The steps of the study is summarized in the workflow figure 1

**Figure 1:**
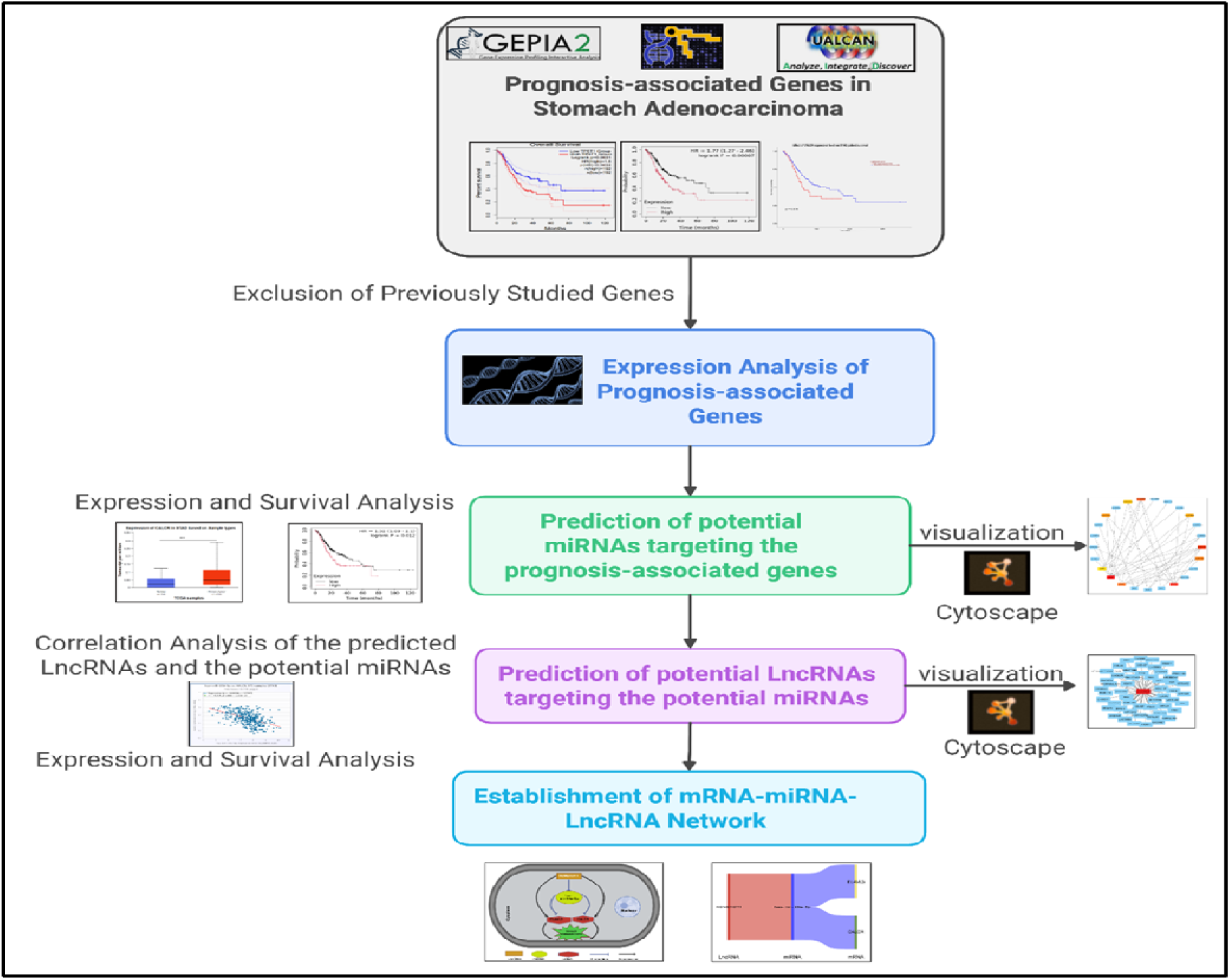
Workflow of the study.

### GEPIA Database Analysis

GEPIA (Gene Expression Profiling Interactive Analysis, http://gepia.cancer-pku.cn/detail.php) is a newly developed interactive web server for analysing the RNA sequencing expression data from the Cancer Genome Atlas (TCGA) and the Genotype-Tissue Expression (GTEx) projects (23). GEPIA was employed to obtain the genes most associated with overall survival and disease-free survival of patients with stomach adenocarcinoma. Logrank P < 0.05 was considered as statistically significant.

### UALCAN Database Analysis

A comprehensive, user-friendly, and interactive web resource for analysing cancer data (24). We used the survival analysis module of UALCAN to study the effect of gene expression on the overall survival in stomach adenocarcinoma. Ualcan is available at (http://ualcan.path.uab.edu/index.html)). P < 0.05 was considered statistically significant.

### MiRNet Database Analysis

MiRNet (http://www.mirnet.ca/), an easy-to-use online tool for miRNA-associated studies, was used to predict potential miRNAs binding to mRNAs as well it used to predict potential lncRNAs binding to miRNAs (25,26).

### StarBase Database Analysis

StarBase (http://starbase.sysu.edu.cn/) is an open-source database for investigating non-coding RNA interactions from CLIP-seq, degradome-seq, and RNA-RNA interactome data (27,28). StarBase was introduced to perform expression correlation analysis for mRNA-miRNA and miRNA-lncRNA pairs in Stomach adenocarcinoma R < 0.1 and P < 0.05 were set as the criteria for identifying significant interactions. MiRNA expression values in stomach adenocarcinoma were also determined using StarBase. P < 0.05 was considered as statistically significant.

### Kaplan-Meier Plotter Analysis

The Kaplan-Meier plotter database is capable of assessing the effect of miRNAs and genes on survival in 21 cancer types, including stomach adenocarcinoma (29). The prognostic values of potential miRNAs in stomach adenocarcinoma were evaluated using the Kaplan-Meier plotter (http://kmplot.com/analysis/). Each miRNA of interest was first submitted to this database. According to the median expression value, all cases were classified into a low expression group and a high expression group. Subsequently, Kaplan-Meier survival plots were generated, and statistical indices contained hazard ratios (HR) and 95% confidence intervals (CI).

### Visualization tools

Cytoscape is a powerful open-source software platform used for visualizing complex networks (30). It is widely used in bioinformatics for visualizing molecular interaction networks and biological pathways. MRNA-miRNA and miRNA-lncRNA regulatory networks were subsequently established. We used cytoscape to visualize the relations between the miRNAs and the corresponding mRNAs and LncRNAs.

Bioinformatics.cn.com is a freely accessible, easy-to-use web server, available at https://www.bioinformatics.com.cn/en. It integrates more than 120 commonly used data visualization and graphing functions together, including heatmaps, Venn diagrams, volcano plots, bubble plots, scatter plots, etc. We used the Sankey diagram from Bioinformatics.cn.com to visualize the interaction between mRNA, miRNA, and LncRNA.

### Statistical Analysis

Most of the statistical analyses were done by the bioinformatics online tools. P-values from GEPIA Expression analysis, logrank P-values from GEPIA, and Kaplan-Meier plotter survival analysis were corrected by false discovery rate, and other reported P-values by online tools were not adjusted for false discovery rate correction.

## Results

### 14 genes were identified as novel prognosis-associated Genes in stomach adenocarcinoma

GEPIA was employed to obtain the genes most associated with overall survival (OS) and disease-free survival (DFS) of patients with stomach adenocarcinoma Logrank P < 0.05 was considered as statistically significant. The 500 OS-associated genes and the top 500 RFS-associated genes were identified as listed in supplementary Tables S1 and S 2 respectively. By intersecting OS-associated genes and RFS-associated genes, we identify 90 -OS and DFS-associated genes. After reviewing the published literature and previous studies, we found that ten genes (CALCR, CFHR1, CPT1C, ELAVL3, FLJ16779, MYOZ3, NALCN, TIGD6, TPST1, and ZNF474) have not been studied for their prognostic values in stomach adenocarcinoma so far. The prognostic values (OS and RFS) of the 10 genes using GEPIA were presented in (Figure S1). The results suggested that high expression of the ten genes indicated poor prognosis in patients with stomach adenocarcinoma carcinoma. Therefore, the ten genes were considered novel potential prognostic biomarkers for stomach adenocarcinoma, and further studies were concentrated on these genes. We further used the Kaplan-Meier plotter and survival module in UALCAN. The results of the Kaplan Meier plotter indicate that all the previously identified prognosis-associated genes by GEPIA were also identified as prognosis-associated genes in the Kaplan Meier plotter. Kaplan-Meier plotter compares the OS between the two groups (upregulated and downregulated expression). We found that the up-regulated group was associated with poor survival and thus poor prognosis (Figure S2). Using the survival analysis module of UALCAN, only 7 out of the 10 previously identified prognosis-associated genes were significantly associated with poor prognosis (NALCN, CALCR, CPT1C, ELAVL3, FLJ16779, MYOZ3, and TPST1) (Figure S3).

### Identification of the upregulated prognosis-associated genes

Next, we intended to determine expression levels of the ten novel genes in stomach adenocarcinoma. We detected their expression in TCGA stomach adenocarcinoma tissues and normal tissues using the UALCAN database, and all genes were significantly upregulated in stomach adenocarcinoma samples compared with normal samples, as shown in figure S4.

### Prediction of Potential miRNAs binding to novel Prognosis-Associated Genes

Next, we predicted upstream regulatory miRNAs of the ten novel prognosis-associated genes through a comprehensive miRNA study-associated database, miRNet. A total of 41 mRNA-miRNA pairs was identified, as shown in Table 1.

**Table 1.** MRNA-miRNA pairs of the novel Prognosis-Associated Genes.

| mRNA | Number of mRNA-miRNA pairs | miRNA |
| --- | --- | --- |
| NALCN | 13 | hsa-miR-15b-5p, hsa-miR-17-5p, hsa-miR-200b-3p, hsa-miR-200c-3p, hsa-miR-20a-5p, hsa-miR-22-3p, hsa-miR-26a-5p, hsa-miR-29a-3p, hsa-miR-29b-3p, hsa-miR-29c-3p, hsa-miR-497-5p, hsa-miR-503-5p, and hsa-miR-9-5p |
| CPT1C | 2 | hsa-miR-17-5p, hsa-miR-26a-5p |
| ELAVL3 | 8 | hsa-miR-15b-5p, hsa-miR-17-5p, hsa-miR-20a-5p, hsa-miR-22-5p, hsa-miR-26a-5p, hsa-miR-29a-3p, hsa-miR-29c-3p, and hsa-miR-503-5p |
| MYOZ3 | 5 | hsa-miR-17-5p, hsa-miR-22-3p, and hsa-miR-26a-5p, hsa-miR-9-5p hsa-mir-22-3p |
| TPST1 | 3 | hsa-miR-17-5p, hsa-miR-200c-3p, hsa-miR-29c-3p |
| TIGD6 | 7 | hsa-mir-10b-5p, hsa-miR-10b-5p, hsa-miR-15b-5p, hsa-miR-17-5p, hsa-miR-20a-5p, hsa-miR-26a-5p and hsa-miR-449a |
| CALCR | 3 | hsa-miR-200c-3p, hsa-miR-200b-3p, hsa-miR-29a-3p |

For better visualization, an mRNA-miRNA interactive network was constructed using cytoscape software, and detailed mRNA-miRNA pairs were shown as presented in Figure 2.

**Figure 2:**
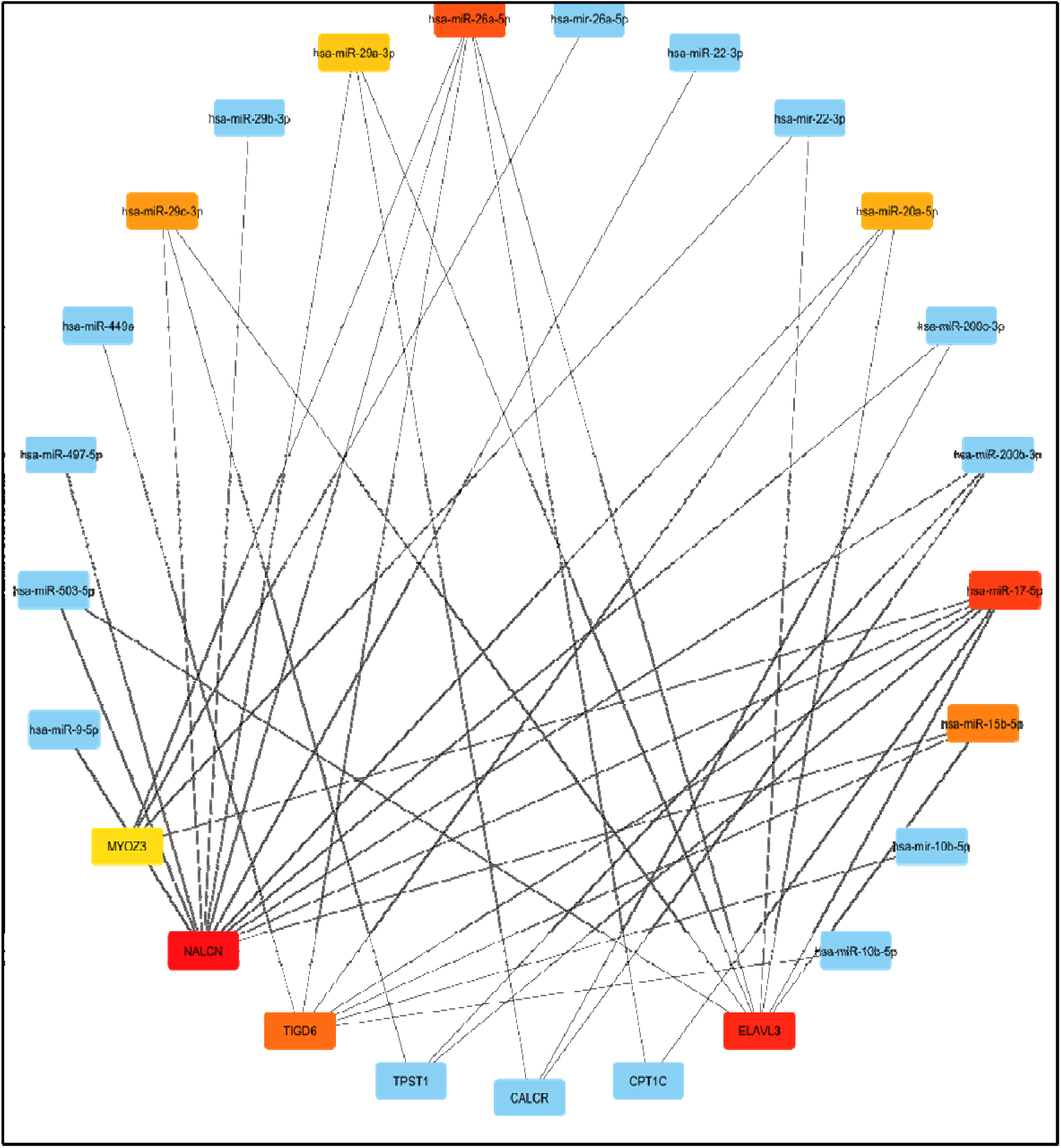
Construction of NALCN/ELAVL3/TPST1/CALCR/MYOZ3/TIGD6 and CPT1C-miRNA network by miRNet database and visualization by Cytoscape software.

According to the classic action mechanism of miRNA in negative regulation of gene expression, there should be an inverse expression relationship between the predicted mRNA-miRNA interactions. Thus, we employed a StarBase database to perform expression correlation analysis for these mRNA-miRNA interactions in stomach adenocarcinoma. Those mRNA-miRNA pairs with R<−0.1 and P<0.05 were considered as significant interactions

Among the 41 interactions, only 19 mRNA-miRNA pairs were identified as significant interactions. As shown in figure S5,

Theoretically, miRNAs that potentially bind to oncogenic genes should be downregulated in stomach adenocarcinoma Expression study of the significant miRNAs was done by the StarBase database and only 5 miRNAs showed down-regulated expression in STAD (hsa-miR-26a-5p, hsa-miR-29c-3p, hsa-miR-497-5p, hsa-miR-9-5p, and hsa-miR-10b-5p) shown in figure S6 while The prognostic roles of these potential miRNAs in stomach adenocarcinoma were determined using Kaplan-Meier plotter database We found only hsa-mir-26a-5p and hsa-mir-29a-3p have down regulated expression, which was associated with poor prognosis. By combination of expression analysis and survival analysis, hsa-mir-29a-3p was the potential miRNA in STAD figure 3.

**Figure 3.**
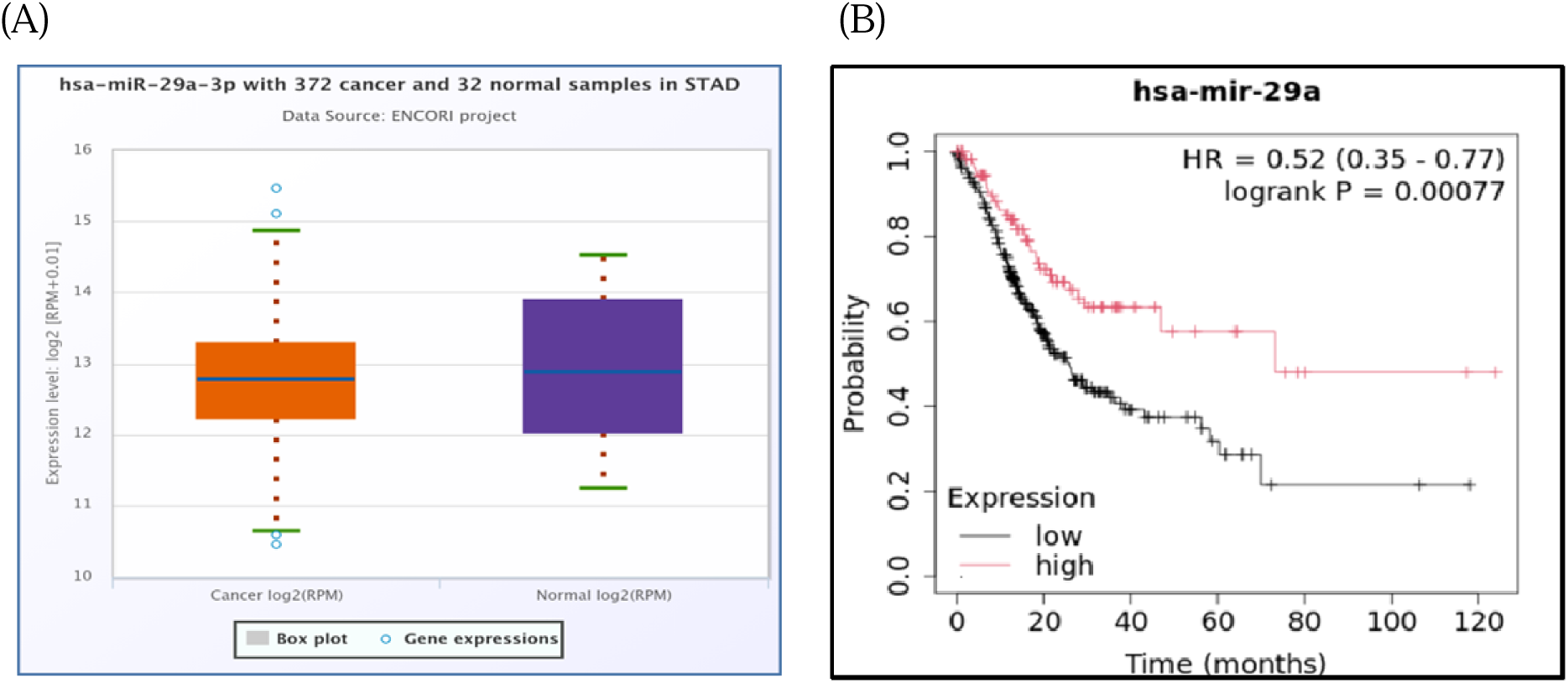
Identification of the most potential miRNAs associated with prognosis of stomach adenocarcinoma. Low expression of hsa-mir-29a-3p in tumor in comparison with normal tissue (A) and prognostic value (B) with downregulated hsa-mir-29a-3p expression in stomach adenocarcinoma was associated with poor survival and worse prognosis in comparison with the upregulated group.

### Prediction of Key LncRNA binding to Potential MiRNA

Previous studies have suggested that LncRNAs can bind to miRNA, and mediate regulation of target gene expression, and play biological roles. Thus, miRNet and starBase databases were used to predict potential lncRNAs that may target hsa-mir-29a-3p.Fifty two and thirty-five LncRNAs were predicted to target hsa-mir-29a-3p by miRNet and starBase, respectively.

As shown in figures 4A–B, 13, lncRNAs binding to hsa-mir-29a-3p commonly appeared in both miRNet and starBase databases. These lncRNAs were selected for subsequent analysis. For better visualization, the miRNA-LncRNA regulatory network was established by Cytoscape software (Figure 4 B).

**Figure 4:**
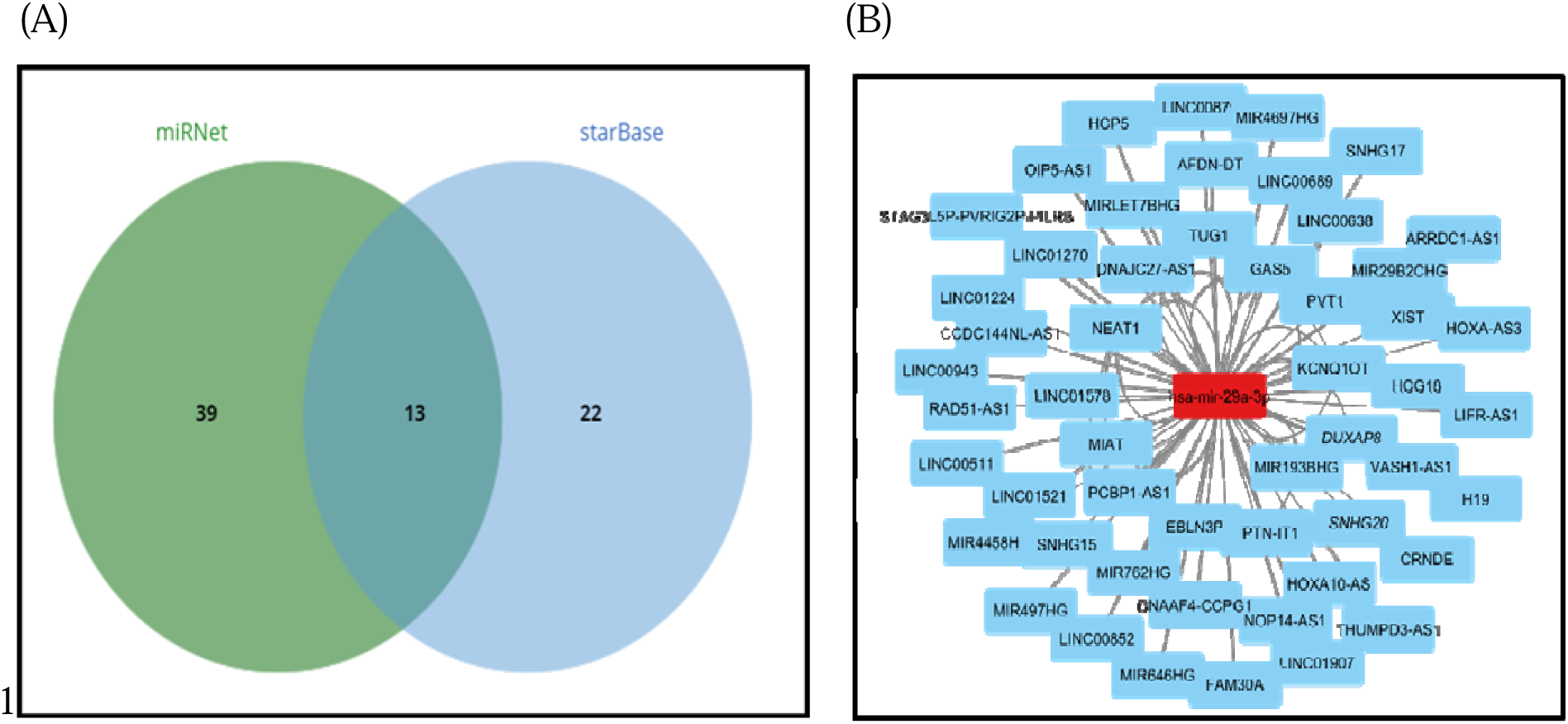
Prediction of upstream lncRNAs potentially binding to hsa-mir-29a-3p(A) miRNA-LncRNA regulatory network was established by Cytoscape software (B)

Based on the ceRNA hypothesis, LncRNAs targeting hsa-mir-29a-3p should be oncogenic LncRNAs in STAD. We found that both KCNQ1OT1 and OIP5-AS1 LncRNAs have a significant negative correlation with hsa-mir-29a-3p (Table 2, Figure 5A&B).

**Figure 5.**
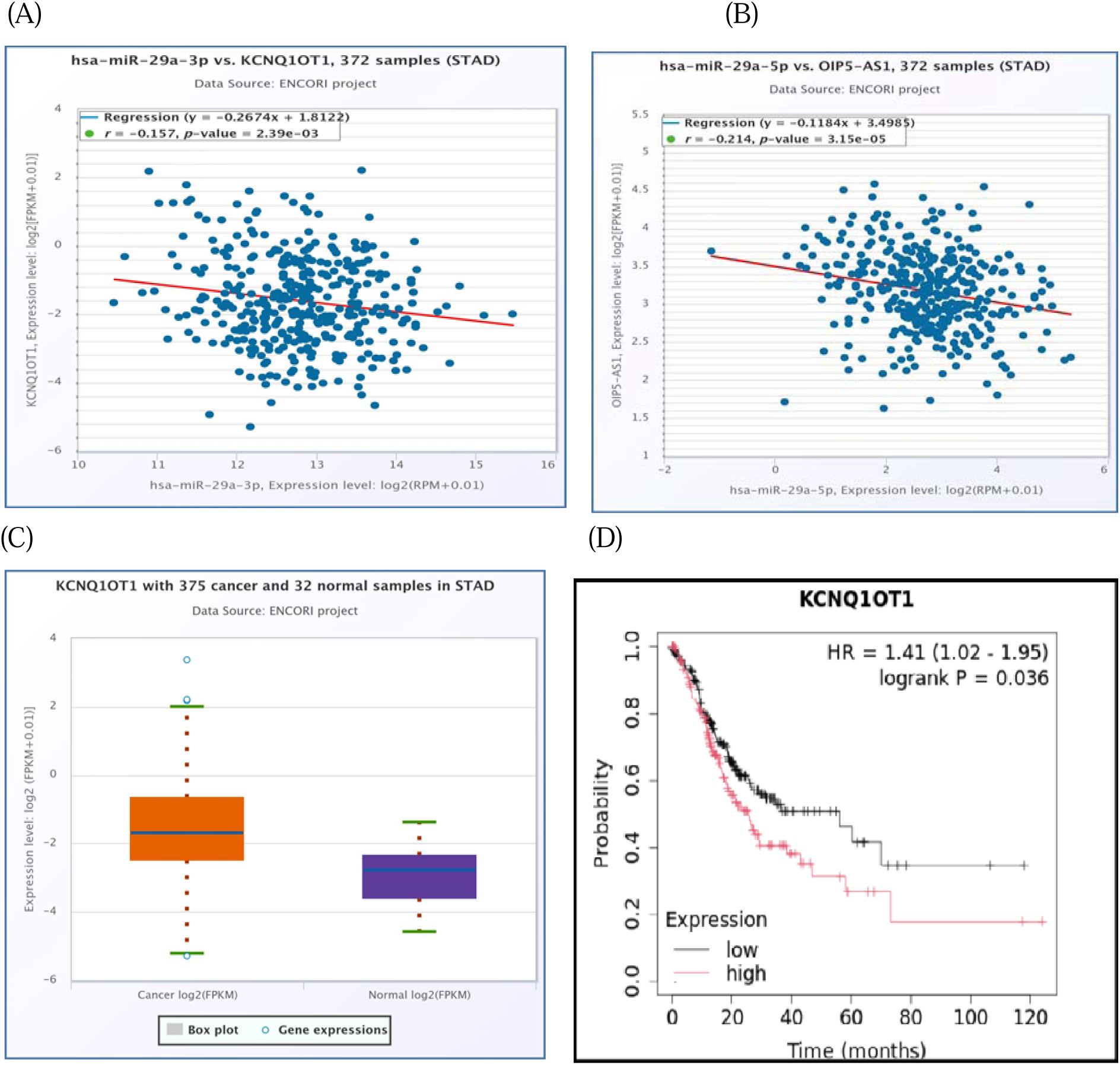
Correlation between the potential miRNAs and predicted LncRNAs using StarBase database (A) correlation between hsa-mir-29a-3p and KCNQ1OT1 LncRNA; (B) correlation between hsa-mir-29a-3p and OIP5-AS1LncRNA. (C) Expression of KCNQ1OT1 LncRNA in STAD using StarBase database. (D) Survival analysis of KCNQ1OT1 in STAD.

**Table 2.**
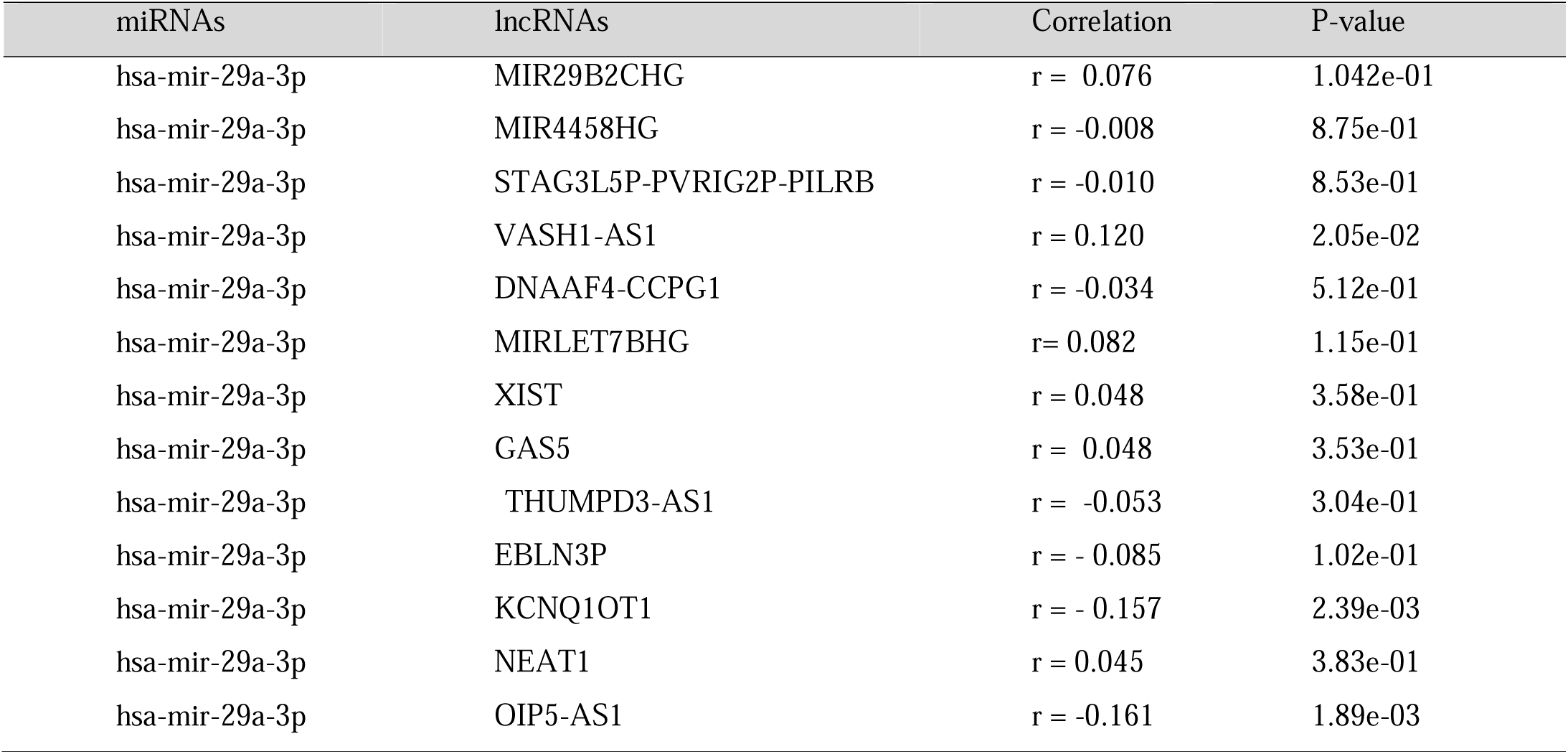
The correlation between potential miRNA-LncRNA pairs identified by starBase (the pairs.

By combination of expression and survival analysis, we found that only one LncRNA, KCNQ1OT1, was significantly upregulated in STAD, and its upregulation was linked to the poor prognosis of patients with STAD (Figure 5-C&D). The current findings support that KCNQ1OT1 was significantly negatively correlated with hsa-mir-29a-3p, upregulated, and linked to poor prognosis in STAD. It might be the most potential LncRNA that binds to previously identified miRNA, hsa-mir-29a-3p.

### Prediction of Upstream miRNA-lncRNA Network of “Known” mRNA in Stomach adenocarcinoma

Next, based on the ceRNA mechanism, we further constructed a “known” genes-miRNA-lncRNA network in stomach adenocarcinoma. Firstly, we predicted upstream miRNAs of NALCN CPT1C ELAVL3 MYOZ3 TPST1 TIGD6 CALCR using miRNet. As shown in Table 1, a total of 41 miRNA-mRNA pairs were predicted. Then, we performed expression correlation analysis for the miRNA-mRNA pairs in stomach adenocarcinoma and found that only 19 pairs presented significantly negative relationships about their expression (Figure S5). Expression and survival analyses revealed that only 1 miRNA (has-mir-29a-3p) was significantly downregulated in stomach adenocarcinoma and correlated with a favorable prognosis (Figure 3).

Next, we predicted the upstream lncRNAs of the potential miRNAs through miRNet and starBase databases. As presented in Figure 4, 13 LncRNA was identified to potentially target hsa-mir-29a-3p (Table 2).

Correlation analysis of the predicted lncRNAs, we found KCNQ1OT1 and OIP5-AS1 were significantly negatively correlated with hsa-mir-29a-3p (Figure 5A,B) by combination of expression and survival analysis we found only KCNQ1OT1was upregulated in STAD and its upregulated expression is linked to poor prognosis (Figure 5 C,D). Establishment of Key mRNA-miRNA-lncRNA Triple ceRNA Network in stomach adenocarcinoma 1 potential lncRNAs KCNQ1OT1 together with 1 potential miRNAs (hsa-mir-29a-3p) made up a miRNA-LncRNA sub-network. According to the ceRNA hypothesis, there should be a positive association between mRNA expression and LncRNA expression. StarBase was employed to analyze the expression correlation of 3 mRNA-LncRNA pairs (KCNQ1OT1-NALCN, KCNQ1OT1-ELAVL3, and KCNQ1OT1-CALCR). Notably, as presented in Figure 8, significant positive expression associations of these mRNA-LncRNA pairs were observed in both ELAVL3 and CALCR.

**Figure 8:**
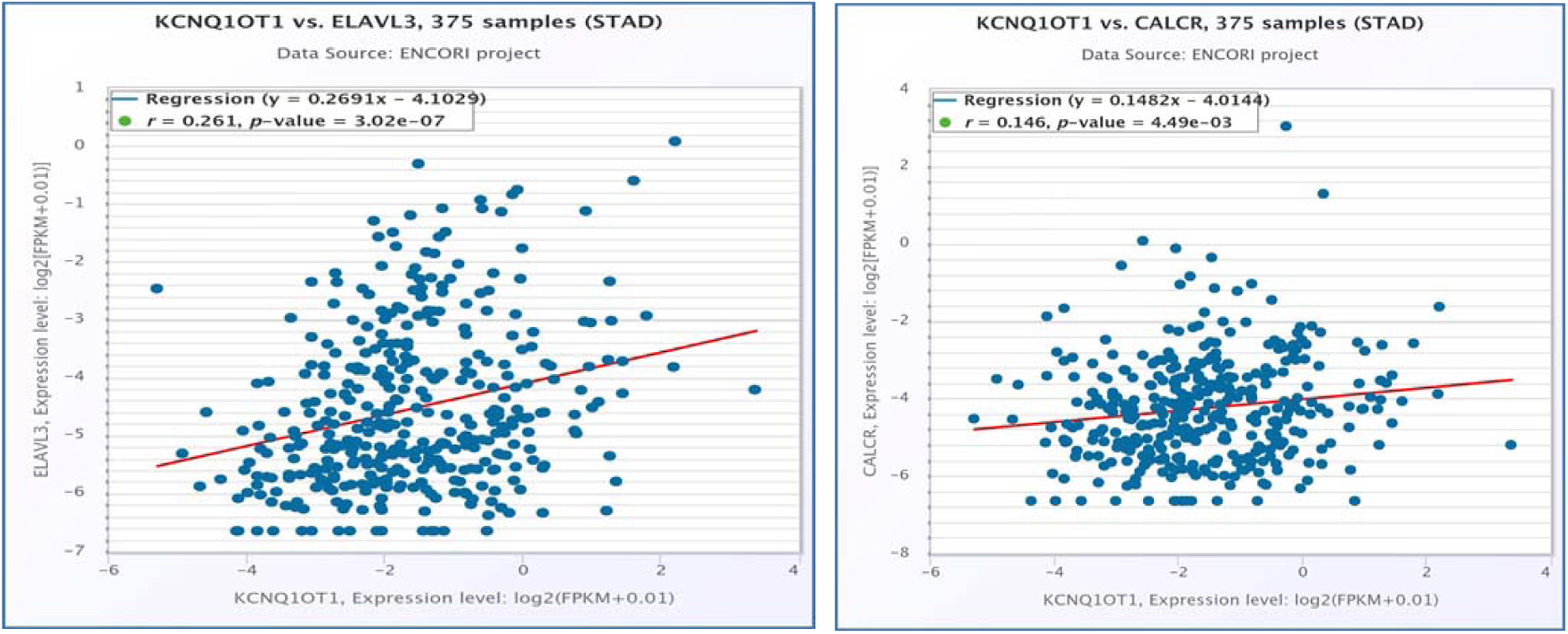
Correlation analysis of potential mRNA-lncRNA pairs in STAD determined by starBase. (A) Expression was significantly positively correlated with ELAVL3 (A) CALCR (B) expression in stomach adenocarcinoma.

By integration of results from in silico analysis, we established a key mRNA-miRNA-lncRNA triple regulatory network associated with the prognosis of stomach adenocarcinoma (Figures 9 and 10).

**Figure 9:**
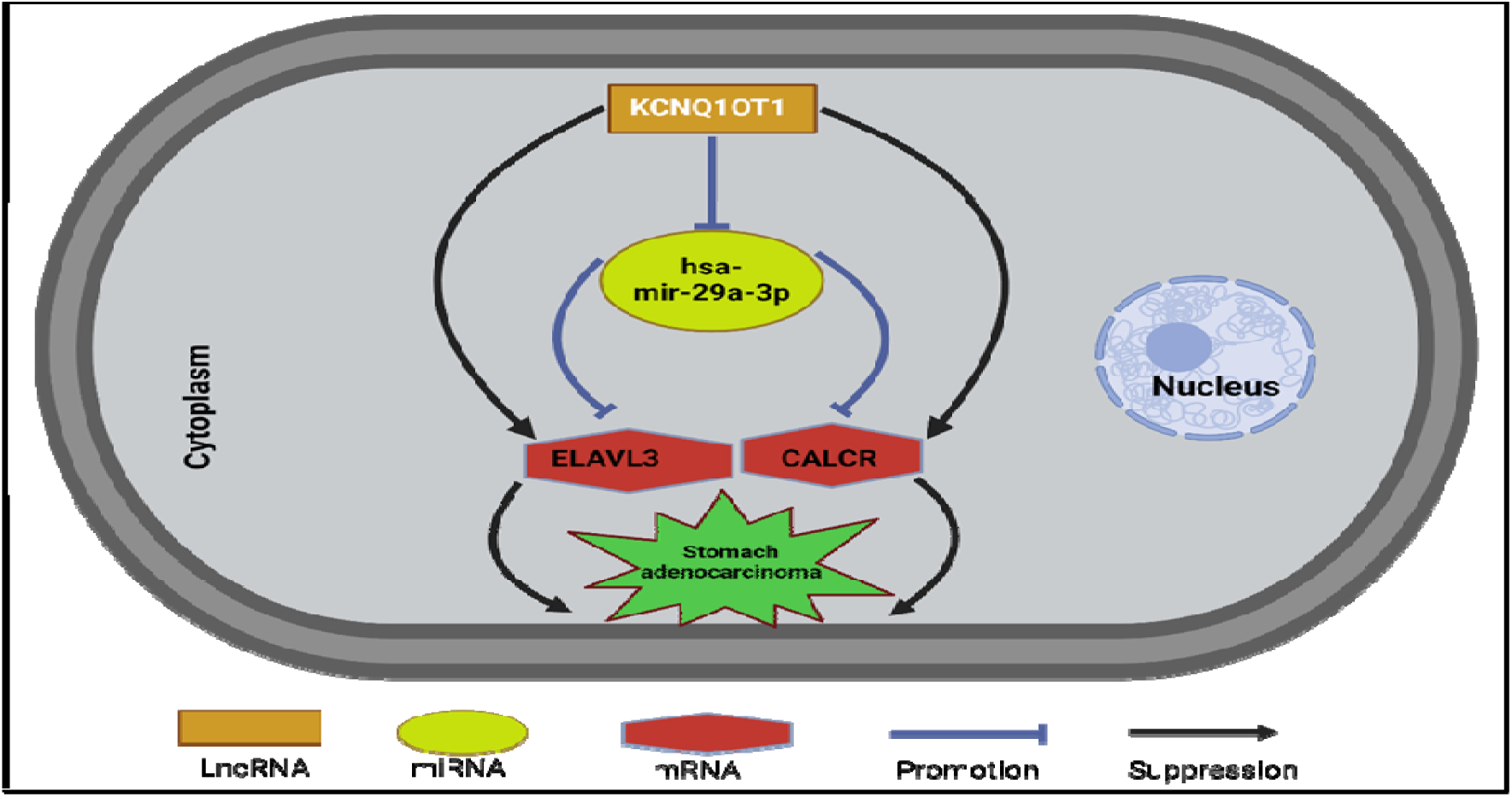
The established mRNA-miRNA-LncRNA competing endogenous RNA (ceRNA) triple network associated with progression and prognosis of stomach adenocarcinoma.

**Figure 10:**
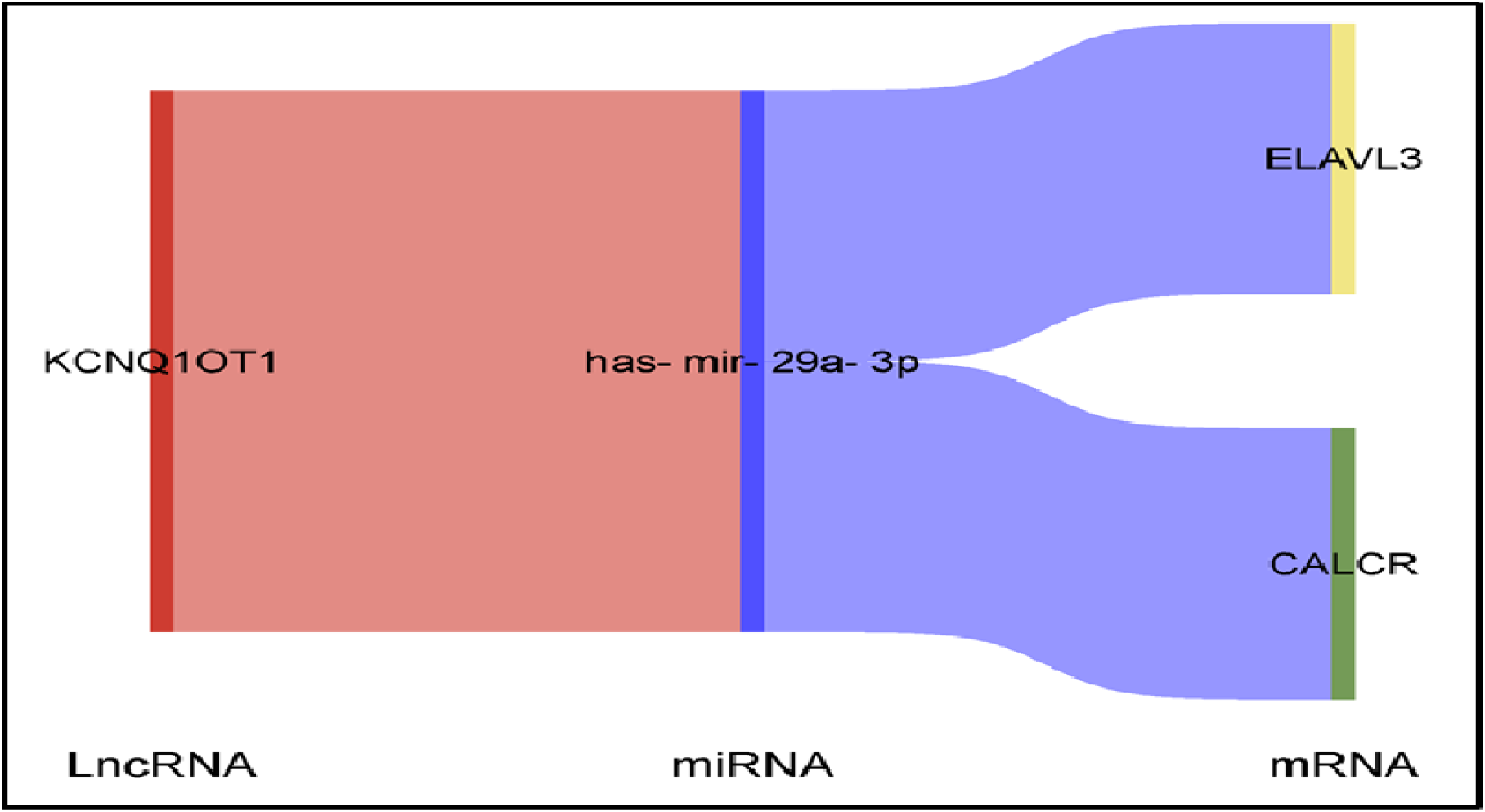
Sankey diagram showing the novel LncRNA-mRNA interaction in STAD.

## Discussion

Stomach adenocarcinoma is characterized by its poor prognosis and aggressive clinical behaviour. Understanding the molecular mechanisms underlying this malignancy and identifying robust biomarkers for diagnosis and prognosis are critical for refining therapeutic strategies and improving patient outcomes. The ceRNA regulatory network is increasingly recognized as a key player in the initiation and progression of human cancers. However, to date, a comprehensive ceRNA regulatory network incorporating the mRNA-miRNA-lncRNA axis in stomach adenocarcinoma remains unexplored. To address this gap, we aimed to construct a prognosis-and diagnosis-associated mRNA-miRNA-lncRNA-ceRNA triple sub-network. By performing survival (OS and RFS) analysis using TCGA stomach adenocarcinoma data, ten novel genes, CALCR, CFHR1, CPT1C, ELAVL3, FLJ16779, MYOZ3, NALCN, TIGD6, TPST1, and ZNF474, were identified as prognosis-associated genes in stomach adenocarcinoma. While these genes have not been previously studied in the context of stomach adenocarcinoma, their oncogenic and biomarker roles have been documented in other cancers. For instance, CALCR overexpression in renal cell carcinoma has been linked to poor prognosis (31). CFHR1 has a documented relevance in lung adenocarcinoma (32), and as a biomarker in bladder cancer (33).

Similarly, elevated ELAVL3 expression correlates with increased cell proliferation and survival in breast cancer (34). The role of FLJ16779 in cancer is not extensively documented in the available literature. FLJ16779 is a gene that encodes a protein, but its specific biological function is not well characterized. While the specific role of FLJ16779 in cancer remains unclear, NALCN is involved in several signalling pathways that are critical for cancer cell behaviour. Its activity can influence pathways related to cell migration, invasion, and the tumor microenvironment (35,36). While the specific role of TIGD6 in cancer is not yet fully understood, the involvement of other TIGD family members in immune modulation and potential impact on tumor progression (37) makes it a candidate for further investigation.

The expression levels of TPST1 may correlate with cancer prognosis. For example, altered levels of TPST1 in tumor tissues could serve as a biomarker for specific cancer types, indicating aggressive behaviour or poor patient outcomes (38)

On the other hand, ZNF proteins promote ovarian cancer cell proliferation and migration (39). Conversely, others act as tumor suppressor in various cancers (40–42).

In silico analysis suggested that the ten genes were significantly upregulated in stomach adenocarcinoma. Collectively, these findings indicate that high expression of CALCR, CFHR1, CPT1C, ELAVL3, FLJ16779, MYOZ3, NALCN, TIGD6, TPST1, and ZNF474 links to poor prognosis in patients with STAD. MRNA can compete with lncRNA by binding to shared miRNAs (7). As such, potential miRNAs of NALCN, CPT1C, ELAVL3, MYOZ3, TPST1, TIGD6, CALCR, and lncRNAs that bind to potential miRNAs can be predicted. 41 miRNAs of NALCN, CPT1C, ELAVL3, MYOZ3, TPST1, TIGD6, and CALCR were initially described using the miRNet database. Considering the action of miRNA on mRNA and presumed oncogenic roles of the prognosis-associated genes, potential miRNAs, being tumor suppressive, should be negatively correlated with NALCN, CPT1C, ELAVL3, MYOZ3, TPST1, TIGD6, and CALCR. Accordingly, we identified 19 potential pairs (NALCN-hsa-miR-15b-5p,NALCN-hsa-miR-17-5p,NALCN-hsa-miR-200b-3p,NALCN-hsa-miR-200c-3p,NALCN-hsa-miR-29a-3p,NALCN-hsa-miR-29b-3p,NALCN-hsa-miR-503-5p,CPT1C-hsa-miR-17-5p,ELAVL3-hsa-miR-15b-5p,ELAVL3-hsa-miR-17-5p,ELAVL3-hsa-miR-20a-5p,ELAVL3-hsa-miR-22-5p,ELAVL3-hsa-miR-29a-3p,ELAVL3-hsa-miR-503-5p,MYOZ3-hsa-miR-17-5p,TPST1-hsa-miR-17-5p,TPST1-hsa-miR-200c-3p, CALCR-hsa-miR-200b-3p, and CALCR-hsa-miR-29a-3p), using correlation analysis for these mRNA-miRNA interactions in STAD. After performing expression and survival analyses, only one of the 19 mRNA-miRNA pairs was selected for subsequent analysis. Hsa-miR-29a-3p has been suggested to regulate the expression of several genes, including those involved in cellular processes such as proliferation, migration, and apoptosis (43). In breast cancer and hepatocellular carcinoma, hsa-miR-29a-3p has been identified as a significant player in tumor progression (44,45). In consistence with our finding, the expressed levels of miR-29a-3p in GC tissue samples were markedly decreased in comparison to that in normal adjacent tissue samples (46).

Together with our results, these studies suggest that CALCR-hsa-miR-29a-3p and ELAVL3-hsa-miR-29a-3p may represent pivotal pathways in the pathogenesis of STAD. Subsequently, lncRNAs interacting with hsa-miR-29a-3p were predicted using miRNet and starBase databases. According to the ceRNA hypothesis, candidate lncRNAs of hsa-mir-29a-3p should act as oncogenic LNCRNAs in STAD. Among all predicted lncRNAs, only one lncRNA (KCNQ1OT1) was significantly upregulated in STAD, and their upregulation was linked to the poor prognosis of patients with STAD.

KCNQ1OT1 as oncogenic LncRNA, has been implicated in critical signaling pathways that promote cancer progression, including the Wnt/β-catenin pathway, which governs cell proliferation and migration (47–50) By integrating these mRNA-miRNA and miRNA-lncRNA interactions, we constructed a potential ceRNA sub-network associated with STAD prognosis. Further studies are needed in the future for experimental validation of these results.

While these findings provide novel insights, they are subject to certain limitations. Experimental validation of the identified ceRNA interactions is necessary to confirm functional relevance. Additionally, further investigation is required to elucidate the molecular roles of less-characterized genes, such as FLJ16779 and TIGD6. Future studies should focus on experimental validation, clinical utility assessment, and therapeutic exploration of the ceRNA network to advance STAD diagnosis and treatment.

## Conclusion

This paper establishes a critical framework for understanding STAD’s ceRNA regulation network. Although more experimental confirmation is required, these findings highlight interesting biomarkers for diagnostic and therapeutic applications, providing new insights into STAD pathophysiology.

## Acknowledgements

We acknowledge and thank all participants for their cooperation and scientific contributions.

## Author contributions

Conceptualization, E.K, Z.O, A.G.; Data curation, E.K, Z.O; Funding acquisition, E.K.; Investigation, E.K, Z.O, A.G, A.O.; Methodology, E.K, Z.O,A.G; Software, E.K; Supervision, AO.; Validation, E.K, Z.O,A.G,A.O; Writing—original draft, E.K, Z.O, A.G, and A.O; Writing—review & editing, E.K.,Z.O,A.G.

## Ethics declarations

No human or animal samples were used in this study

## Bibliography

1. Sung H, Ferlay J, Siegel RL, Laversanne M, Soerjomataram I, Jemal A, et al. Global cancer statistics 2020: GLOBOCAN estimates of incidence and mortality worldwide for 36 cancers in 185 countries. CA Cancer J Clin. 2021 May;71(3):209–49.

2. Thrift AP, Wenker TN, El-Serag HB. Global burden of gastric cancer: epidemiological trends, risk factors, screening and prevention. Nat Rev Clin Oncol. 2023 May;20(5):338–49.

3. Wang S, Zheng R, Li J, Zeng H, Li L, Chen R, et al. Global, regional, and national lifetime risks of developing and dying from gastrointestinal cancers in 185 countries: a population-based systematic analysis of GLOBOCAN. Lancet Gastroenterol Hepatol. 2024 Mar;9(3):229– 37.

4. Bhan A, Hussain I, Ansari KI, Kasiri S, Bashyal A, Mandal SS. Antisense transcript long noncoding RNA (lncRNA) HOTAIR is transcriptionally induced by estradiol. J Mol Biol. 2013 Oct 9;425(19):3707–22.

5. Pérez-Navarro Y, Salinas-Vera YM, López-Camarillo C, Figueroa-Angulo EE, Alvarez-Sánchez ME. The role of long non-coding RNA NORAD in digestive system tumors. Noncoding RNA Res. 2025 Feb;10:55–62.

6. Piazuelo MB, Correa P. Gastric cáncer: Overview. Colomb Med (Cali). 2013 Jul;44(3):192– 201.

7. Guan W-L, He Y, Xu R-H. Gastric cancer treatment: recent progress and future perspectives. J Hematol Oncol. 2023 May 27;16(1):57.

8. Yang W-J, Zhao H-P, Yu Y, Wang J-H, Guo L, Liu J-Y, et al. Updates on global epidemiology, risk and prognostic factors of gastric cancer. World J Gastroenterol. 2023 Apr 28;29(16):2452–68.

9. Jin X, Liu Z, Yang D, Yin K, Chang X. Recent progress and future perspectives of immunotherapy in advanced gastric cancer. Front Immunol. 2022 Jul 1;13:948647.

10. Kono K. Advances in cancer immunotherapy for gastroenterological malignancy. Ann Gastroenterol Surg. 2018 Jul;2(4):244–5.

11. Liu YJ, Shen D, Yin X, Gavine P, Zhang T, Su X, et al. HER2, MET and FGFR2 oncogenic driver alterations define distinct molecular segments for targeted therapies in gastric carcinoma. Br J Cancer. 2014 Mar 4;110(5):1169–78.

12. Taieb J, Bennouna J, Penault-Llorca F, Basile D, Samalin E, Zaanan A. Treatment of gastric adenocarcinoma: A rapidly evolving landscape. Eur J Cancer. 2023 Dec;195:113370.

13. Pihlak R, Fong C, Starling N. Targeted therapies and developing precision medicine in gastric cancer. Cancers (Basel). 2023 Jun 19;15(12).

14. Salmena L, Poliseno L, Tay Y, Kats L, Pandolfi PP. A ceRNA hypothesis: the Rosetta Stone of a hidden RNA language? Cell. 2011 Aug 5;146(3):353–8.

15. Pu X, Sheng S, Fu Y, Yang Y, Xu G. Construction of circRNA-miRNA-mRNA ceRNA regulatory network and screening of diagnostic targets for tuberculosis. Ann Med. 2024 Dec;56(1):2416604.

16. Zhao Y, Wang H, Wu C, Yan M, Wu H, Wang J, et al. Construction and investigation of lncRNA-associated ceRNA regulatory network in papillary thyroid cancer. Oncol Rep. 2018 Mar;39(3):1197–206.

17. Zhang J, Lou W. A Key mRNA-miRNA-lncRNA Competing Endogenous RNA Triple Sub-network Linked to Diagnosis and Prognosis of Hepatocellular Carcinoma. Front Oncol. 2020 Mar 17;10:340.

18. Wang W, Lou W, Ding B, Yang B, Lu H, Kong Q, et al. A novel mRNA-miRNA-lncRNA competing endogenous RNA triple sub-network associated with prognosis of pancreatic cancer. Aging (Albany NY). 2019 May 6;11(9):2610–27.

19. Li X, Dai D, Wang H, Wu B, Wang R. Identification of prognostic signatures associated with long-term overall survival of thyroid cancer patients based on a competing endogenous RNA network. Genomics. 2020 Mar;112(2):1197–207.

20. Wang H, Zhang H, Zeng J, Tan Y. ceRNA network analysis reveals prognostic markers for glioblastoma. Oncol Lett. 2019 Jun;17(6):5545–57.

21. Paul Y, Thomas S, Patil V, Kumar N, Mondal B, Hegde AS, et al. Genetic landscape of long noncoding RNA (lncRNAs) in glioblastoma: identification of complex lncRNA regulatory networks and clinically relevant lncRNAs in glioblastoma. Oncotarget. 2018 Jul 3;9(51):29548–64.

22. Song H, Sun J, Kong W, Ji Y, Xu D, Wang J. Construction of a circRNA-Related ceRNA Prognostic Regulatory Network in Breast Cancer. Onco Targets Ther. 2020 Aug 20;13:8347– 58.

23. Tang Z, Li C, Kang B, Gao G, Li C, Zhang Z. GEPIA: a web server for cancer and normal gene expression profiling and interactive analyses. Nucleic Acids Res. 2017 Jul 3;45(W1):W98–102.

24. Chandrashekar DS, Karthikeyan SK, Korla PK, Patel H, Shovon AR, Athar M, et al. UALCAN: An update to the integrated cancer data analysis platform. Neoplasia. 2022 Mar;25:18–27.

25. Chang L, Zhou G, Soufan O, Xia J. miRNet 2.0: network-based visual analytics for miRNA functional analysis and systems biology. Nucleic Acids Res. 2020 Jul 2;48(W1):W244–51.

26. Fan Y, Siklenka K, Arora SK, Ribeiro P, Kimmins S, Xia J. miRNet -dissecting miRNA-target interactions and functional associations through network-based visual analysis. Nucleic Acids Res. 2016 Jul 8;44(W1):W135–41.

27. Li J-H, Liu S, Zhou H, Qu L-H, Yang J-H. starBase v2.0: decoding miRNA-ceRNA, miRNA-ncRNA and protein-RNA interaction networks from large-scale CLIP-Seq data. Nucleic Acids Res. 2014 Jan;42(Database issue):D92–7.

28. Yang J-H, Li J-H, Shao P, Zhou H, Chen Y-Q, Qu L-H. starBase: a database for exploring microRNA-mRNA interaction maps from Argonaute CLIP-Seq and Degradome-Seq data. Nucleic Acids Res. 2011 Jan;39(Database issue):D202–9.

29. Dudley WN, Wickham R, Coombs N. An Introduction to Survival Statistics: Kaplan-Meier Analysis. J Adv Pract Oncol. 2016 Jan 1;7(1):91–100.

30. Doncheva NT, Morris JH, Holze H, Kirsch R, Nastou KC, Cuesta-Astroz Y, et al. Cytoscape stringApp 2.0: Analysis and Visualization of Heterogeneous Biological Networks. J Proteome Res. 2023 Feb 3;22(2):637–46.

31. Yan H, Xing Z, Liu S, Gao P, Wang Q, Guo G. CALCR exacerbates renal cell carcinoma progression via stabilizing CD44. Aging (Albany NY). 2024 Jul 9;16(13):10765–83.

32. Wu G, Yan Y, Wang X, Ren X, Chen X, Zeng S, et al. CFHR1 is a potentially downregulated gene in lung adenocarcinoma. Mol Med Report. 2019 Oct;20(4):3642–8.

33. Zhang J, Guo F, Zhu J, He Z, Hao L, Weng L, et al. Ultrasensitive Electrochemiluminescence Immunosensor for Bladder Marker Human Complement Factor H-Related Protein Detection. Anal Chem. 2023 Aug 1;95(30):11440–8.

34. Lord BD, Harris AR, Ambs S. The impact of social and environmental factors on cancer biology in Black Americans. Cancer Causes Control. 2023 Mar;34(3):191–203.

35. He J, Xu J, Chang Z, Yan J, Zhang L, Qin Y. NALCN is a potential biomarker and therapeutic target in human cancers. Front Genet. 2023 Apr 19;14:1164707.

36. Rahrmann EP, Shorthouse D, Jassim A, Hu LP, Ortiz M, Mahler-Araujo B, et al. The NALCN channel regulates metastasis and nonmalignant cell dissemination. Nat Genet. 2022 Dec;54(12):1827–38.

37. Xia L, Yang Z, Xv M, Wang G, Mao Y, Yang Y, et al. Bioinformatics analysis and experimental verification of TIGD1 in non-small cell lung cancer. Front Med (Lausanne). 2024 Apr 8;11:1374260.

38. Jiang Z, Zhu J, Ma Y, Hong C, Xiao S, Jin L. Tyrosylprotein sulfotransferase 1 expression is negatively correlated with c Met and lymph node metastasis in human lung cancer. Mol Med Report. 2015 Oct;12(4):5217–22.

39. Zhang J, Wu X, Huang L. ZNF574 Promotes Ovarian Cancer Cell Proliferation and Migration through Regulating AKT and AMPK Signaling Pathways. Ann Clin Lab Sci. 2022 Jul;52(4):611–20.

40. Tao C, Luo J, Tang J, Zhou D, Feng S, Qiu Z, et al. The tumor suppressor Zinc finger protein 471 suppresses breast cancer growth and metastasis through inhibiting AKT and Wnt/β-catenin signaling. Clin Epigenetics. 2020 Nov 17;12(1):173.

41. Cao L, Wang S, Zhang Y, Wong K-C, Nakatsu G, Wang X, et al. Zinc-finger protein 471 suppresses gastric cancer through transcriptionally repressing downstream oncogenic PLS3 and TFAP2A. Oncogene. 2018 Jun;37(26):3601–16.

42. Cao L, Wang S, Zhang Y, Wong K-C, Nakatsu G, Wang X, et al. Correction to: Zinc-finger protein 471 suppresses gastric cancer through transcriptionally repressing downstream oncogenic PLS3 and TFAP2A. Oncogene. 2022 Jun;41(23):3298–9.

43. Mo W-Y, Cao S-Q. MiR-29a-3p: a potential biomarker and therapeutic target in colorectal cancer. Clin Transl Oncol. 2023 Mar;25(3):563–77.

44. Lin G, Lin L, Lin H, Xu Y, Chen W, Liu Y, et al. C1QTNF6 regulated by miR-29a-3p promotes proliferation and migration in stage I lung adenocarcinoma. BMC Pulm Med. 2022 Jul 25;22(1):285.

45. Zhang H, Du Y, Xin P, Man X. The LINC00852/miR-29a-3p/JARID2 axis regulates the proliferation and invasion of prostate cancer cell. BMC Cancer. 2022 Dec 5;22(1):1269.

46. Pan H, Ding Y, Jiang Y, Wang X, Rao J, Zhang X, et al. LncRNA LIFR-AS1 promotes proliferation and invasion of gastric cancer cell via miR-29a-3p/COL1A2 axis. Cancer Cell Int. 2021 Jan 6;21(1):7.

47. Cagle P, Qi Q, Niture S, Kumar D. KCNQ1OT1: an oncogenic long noncoding RNA. Biomolecules. 2021 Oct 29;11(11).

48. Jiang L, Jin H, Gong S, Han K, Li Z, Zhang W, et al. LncRNA KCNQ1OT1-mediated cervical cancer progression by sponging miR-1270 as a ceRNA of LOXL2 through PI3k/Akt pathway. J Obstet Gynaecol Res. 2022 Apr;48(4):1001–10.

49. Wang X, Ren Z, Xu Y, Gao X, Huang H, Zhu F. KCNQ1OT1 sponges miR-34a to promote malignant progression of malignant melanoma via upregulation of the STAT3/PD-L1 axis. Environ Toxicol. 2023 Feb;38(2):368–80.

50. Tang W, Zhang L, Li J, Guan Y. KCNQ1OT1 promotes retinoblastoma progression by targeting miR-339-3p that suppresses KIF23. Int Ophthalmol. 2023 Jul;43(7):2419–32.

