## supplementary figures for "A Bioinformatics-Driven ceRNA Network in Stomach Adenocarcinoma: Identification of Novel Prognostic mRNA-miRNA-lncRNA Interactions"

CALCR

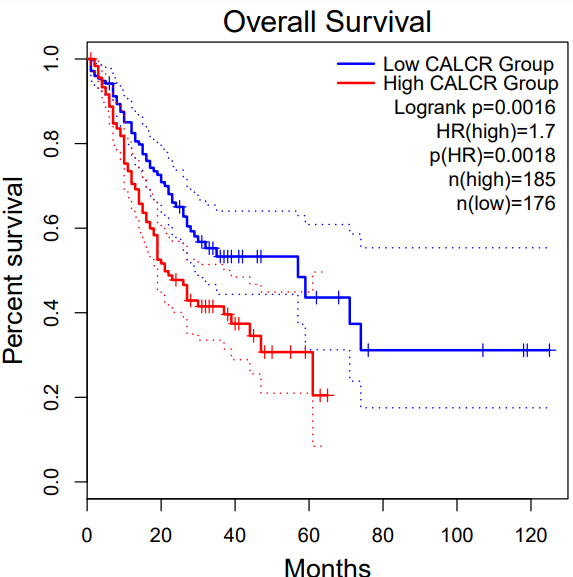

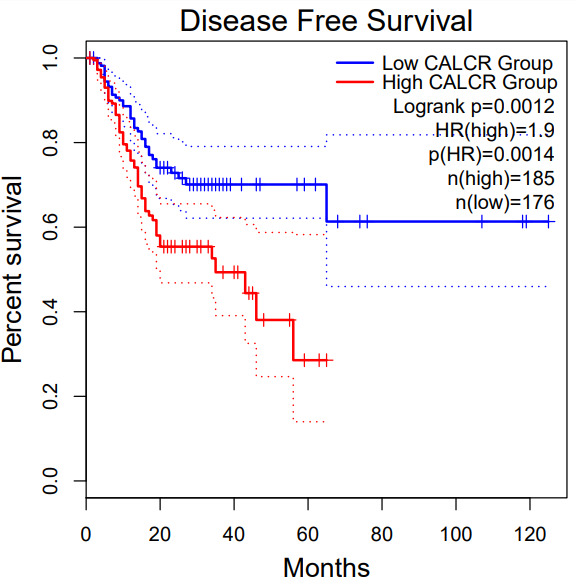

CFHR1

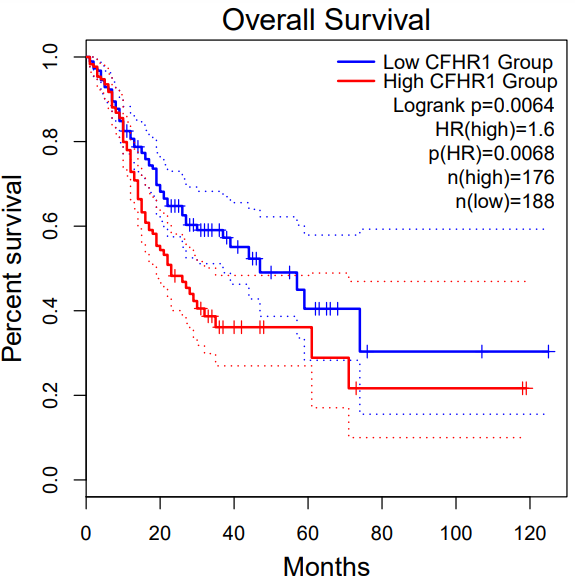

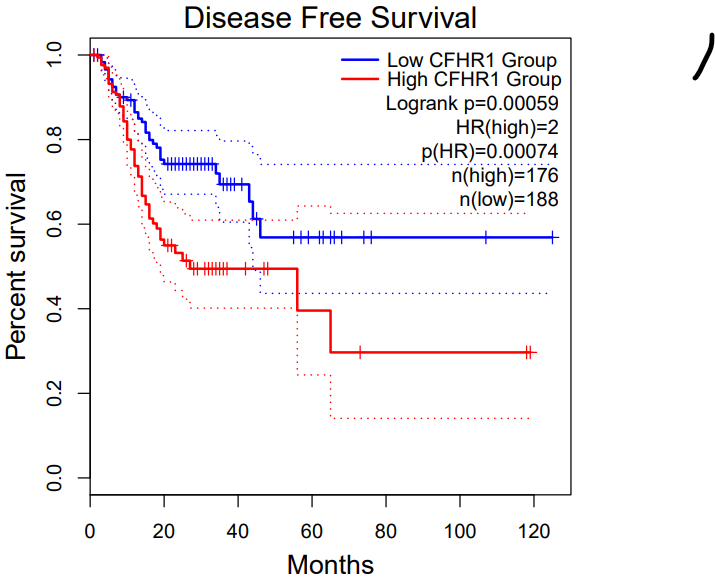

CPT1C

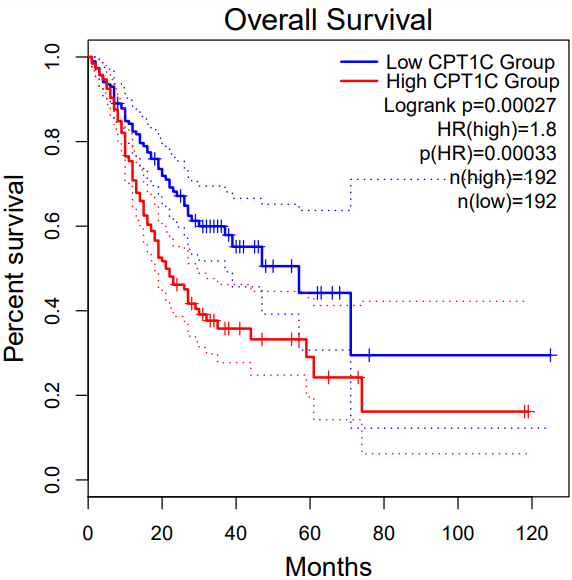

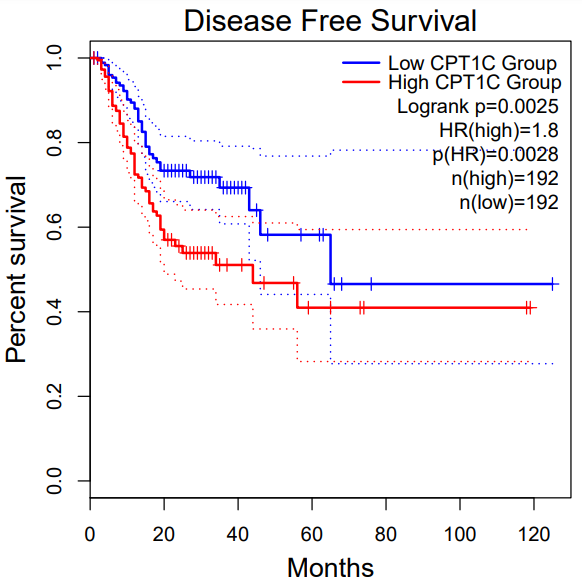

ELAVL3

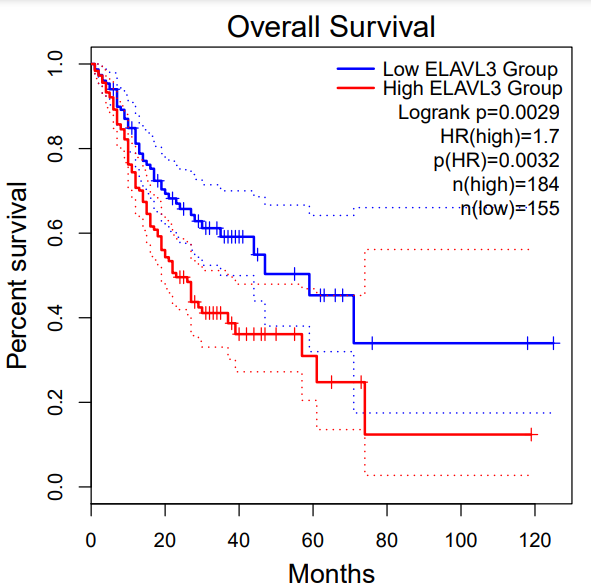

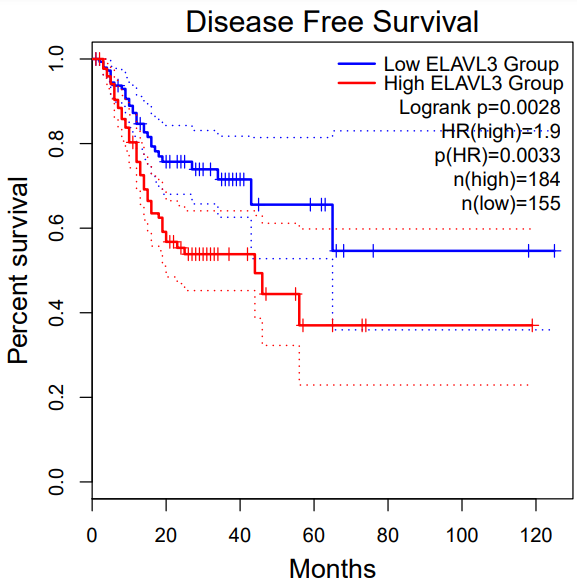

FLJ16779

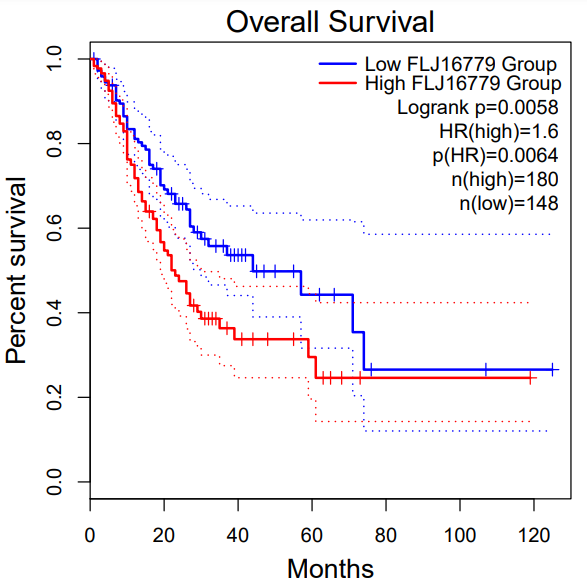

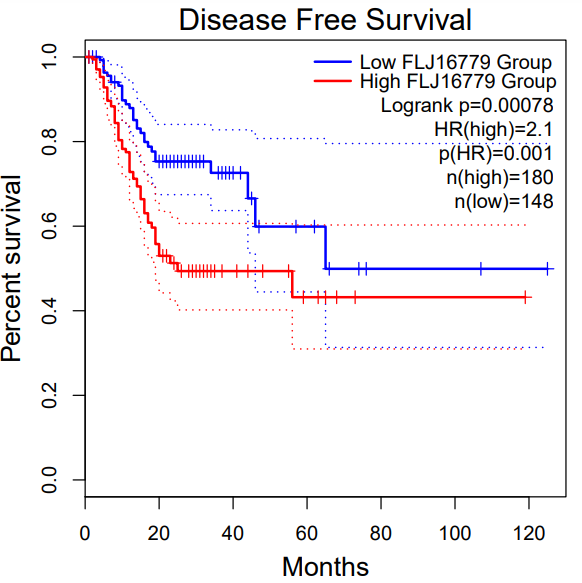

MYOZ3

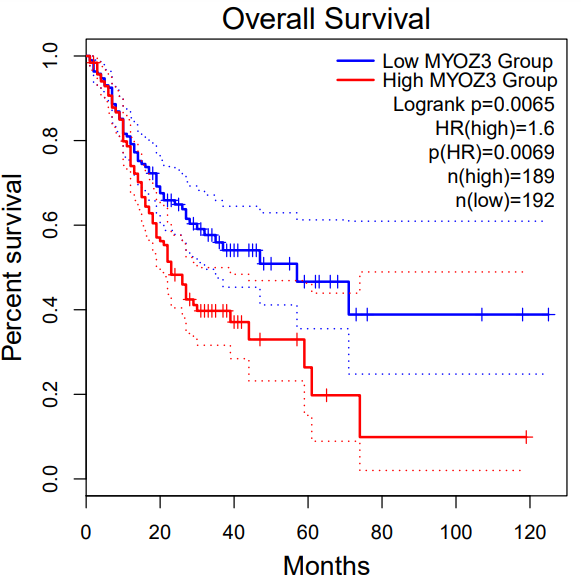

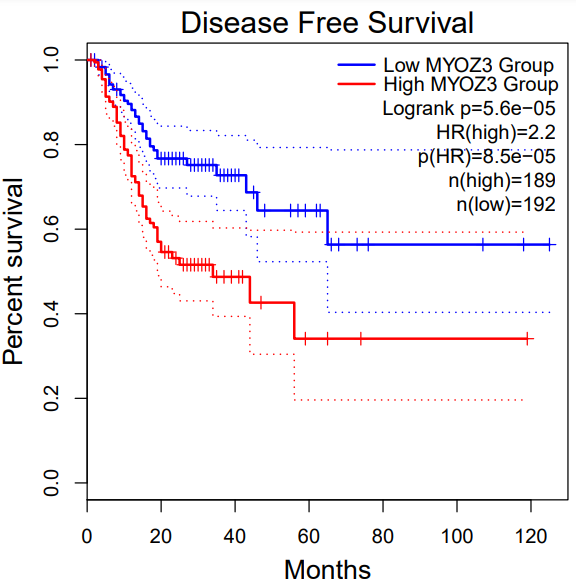

NALCN

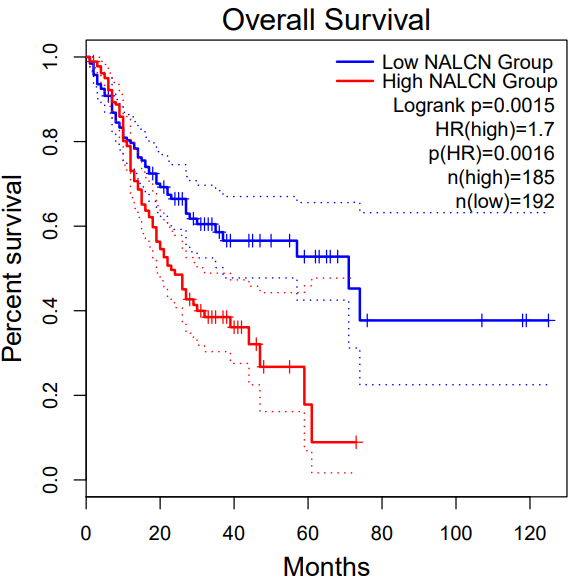

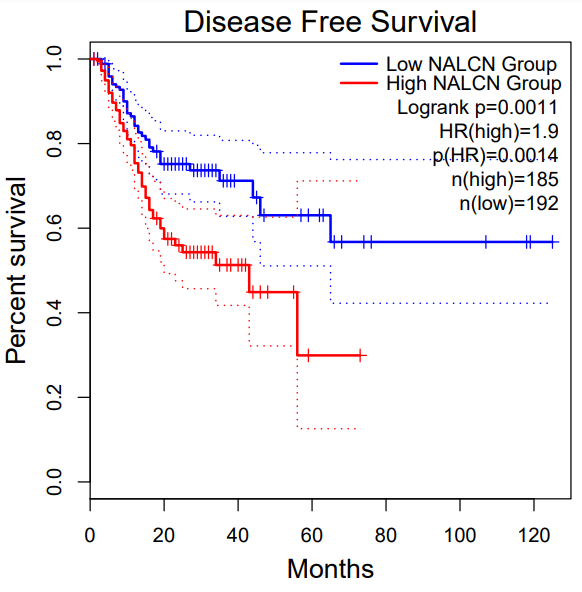

TIGD6

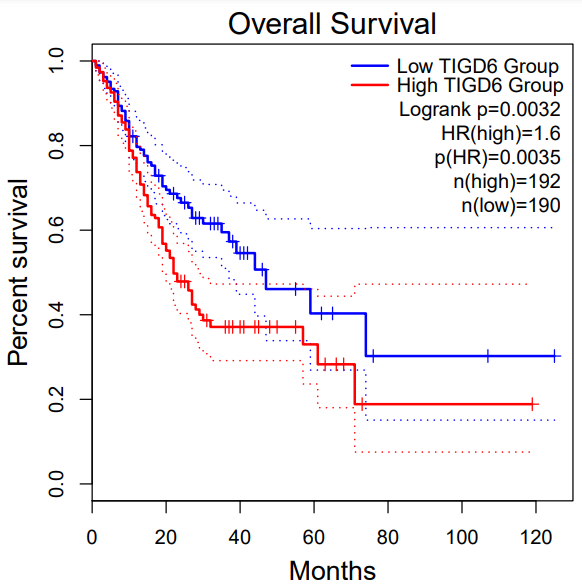

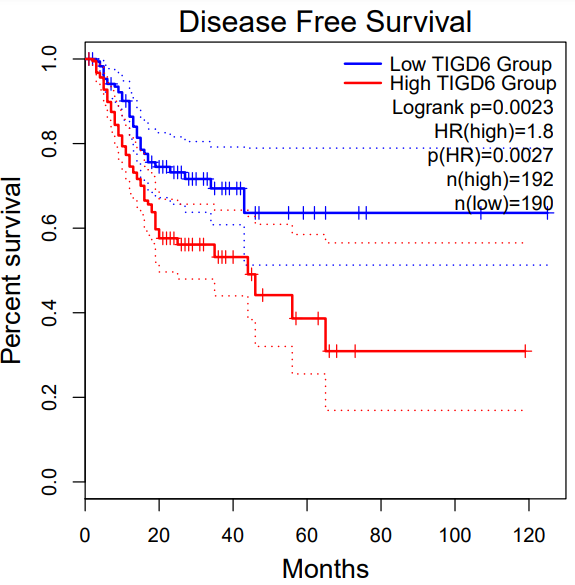

TPST1

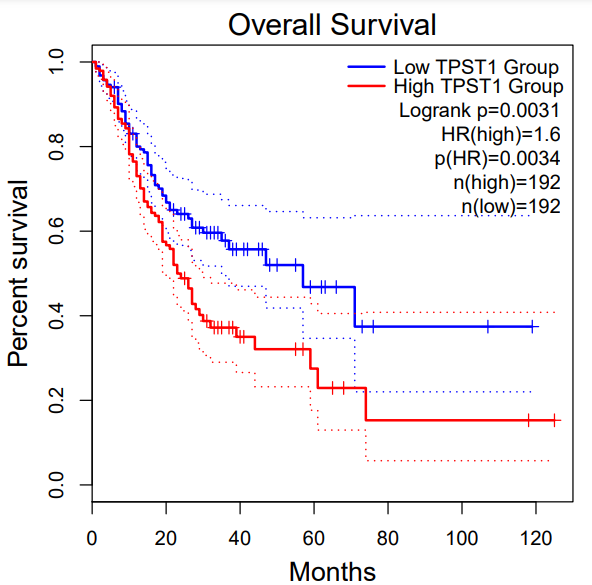

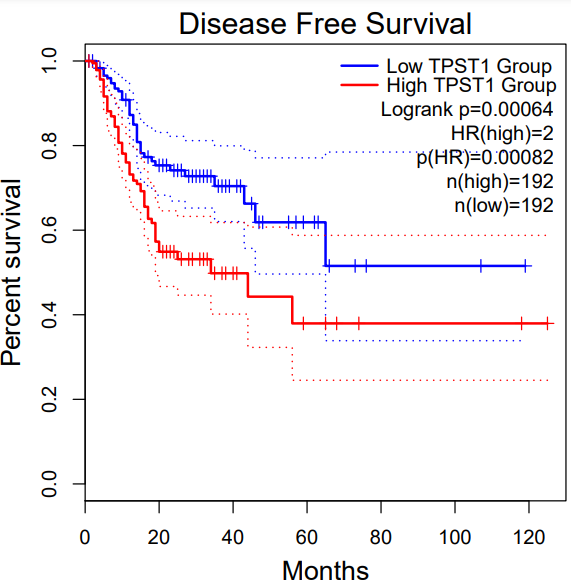

ZNF474

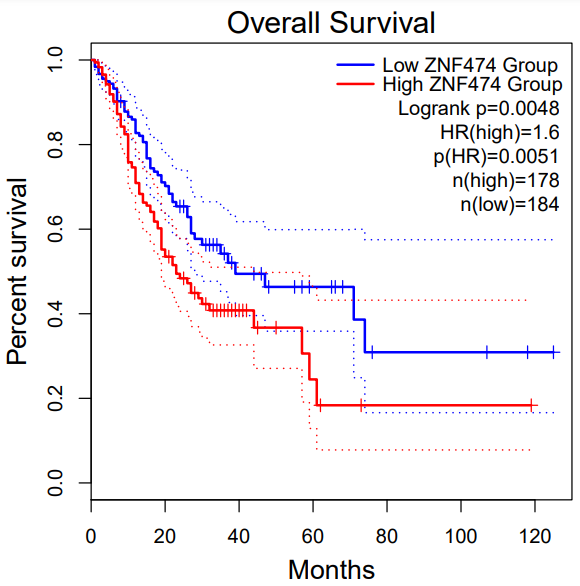

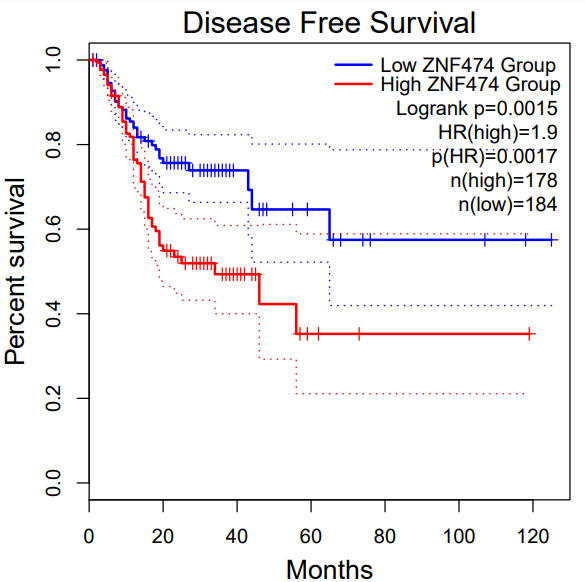

Figure S1: Shows high expression of the 10 novel genes, significantly associated with poor prognosis of stomach adenocarcinoma. Using Gepia overall survival (OS) and disease free survival (DFS).

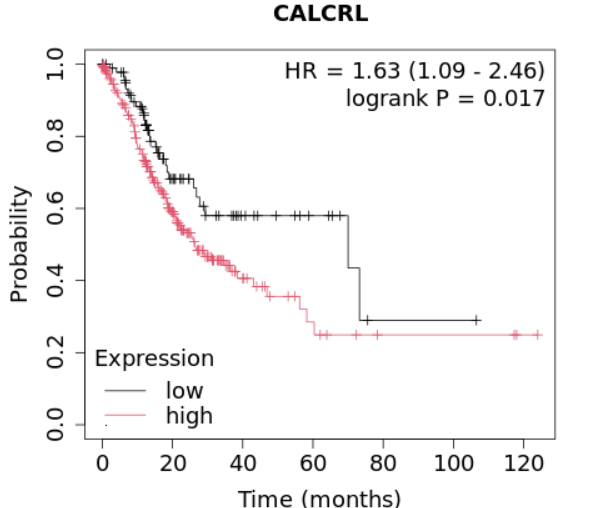

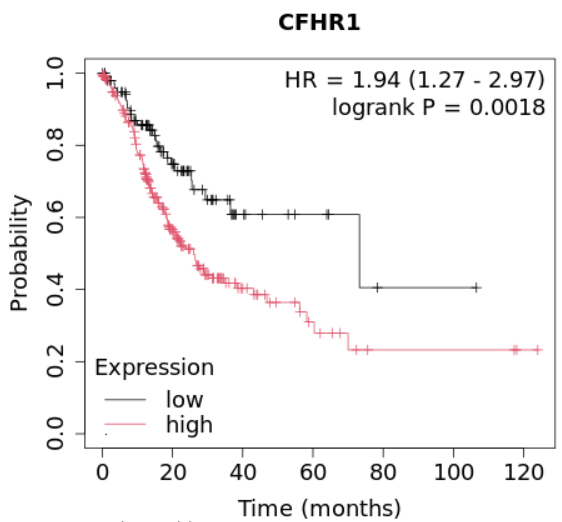

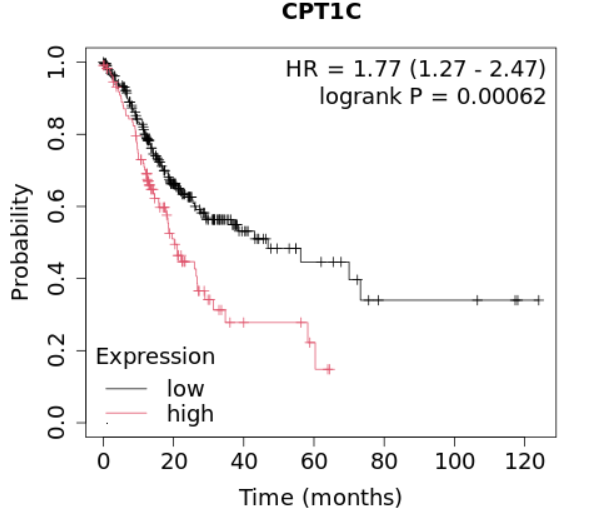

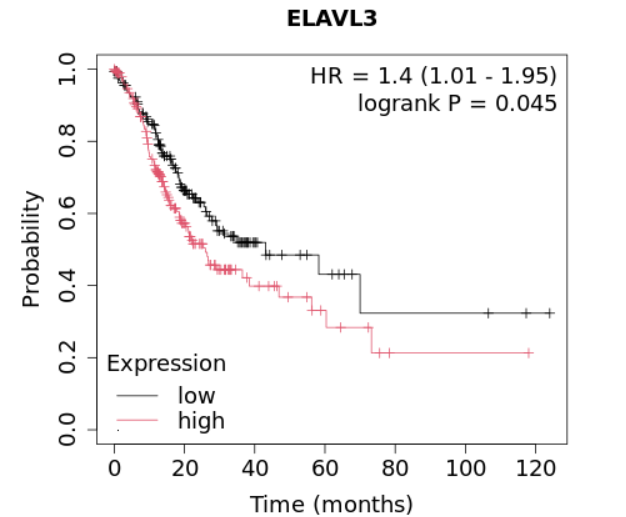

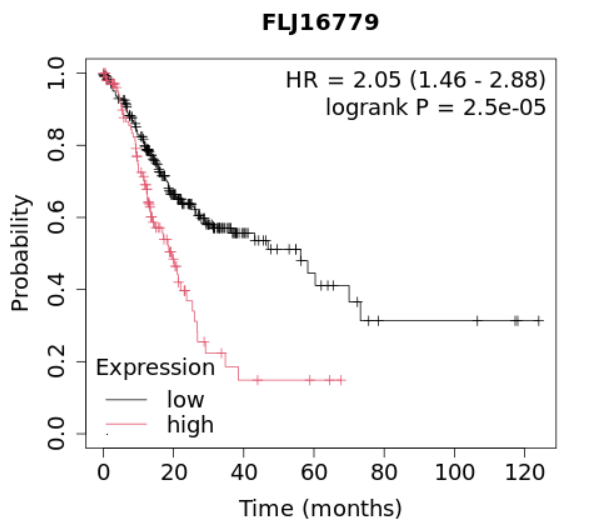

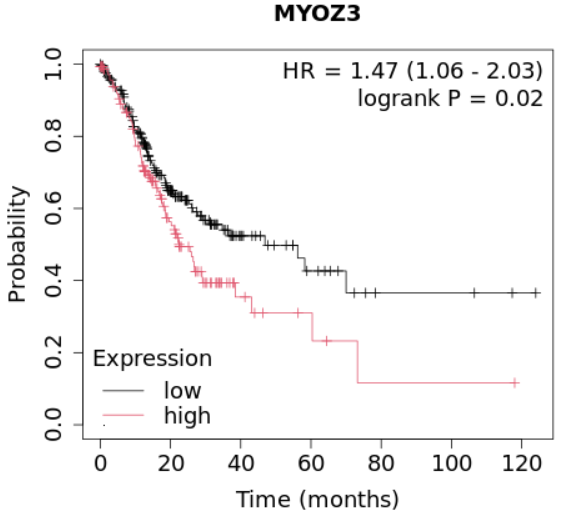

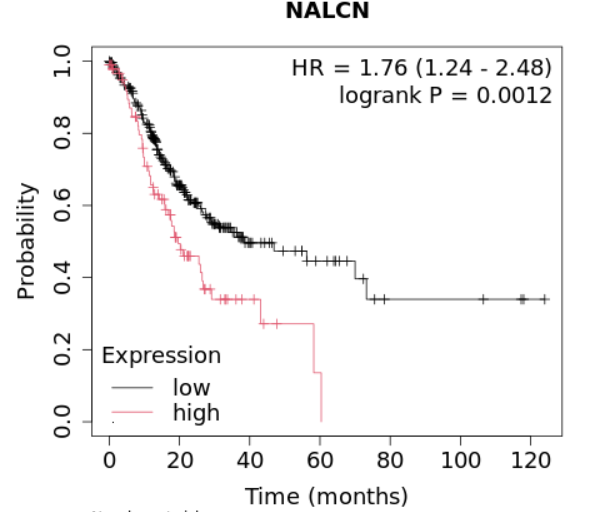

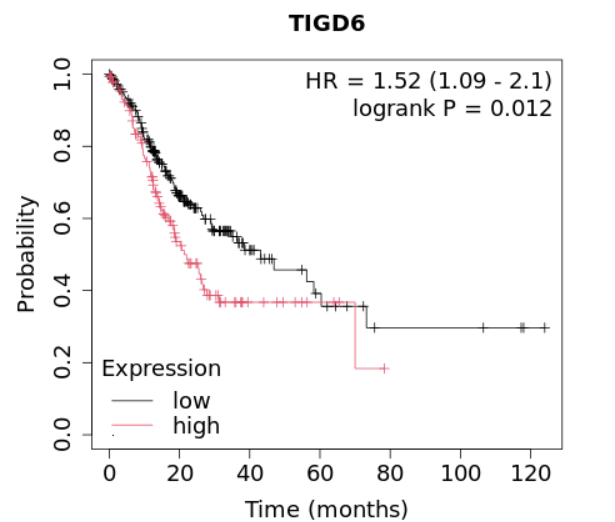

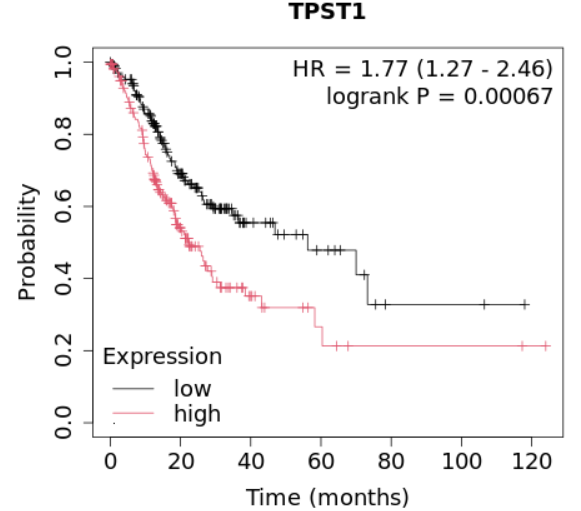

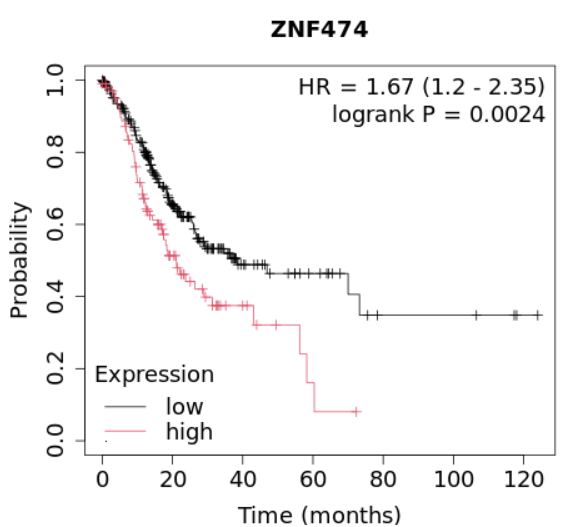

Figure S2: Shows high expression of the 10 novel genes significantly associated with poor prognosis of stomach adenocarcinoma

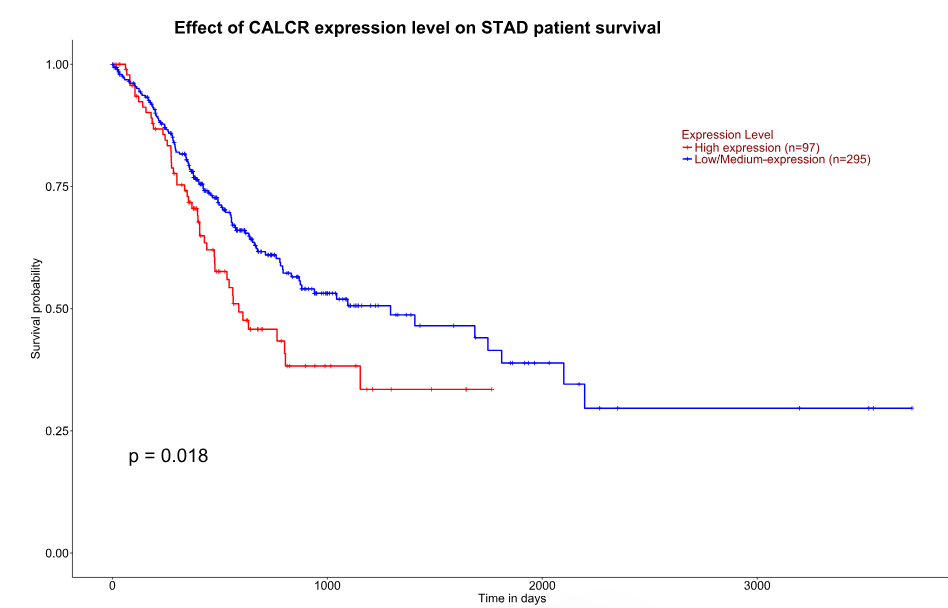

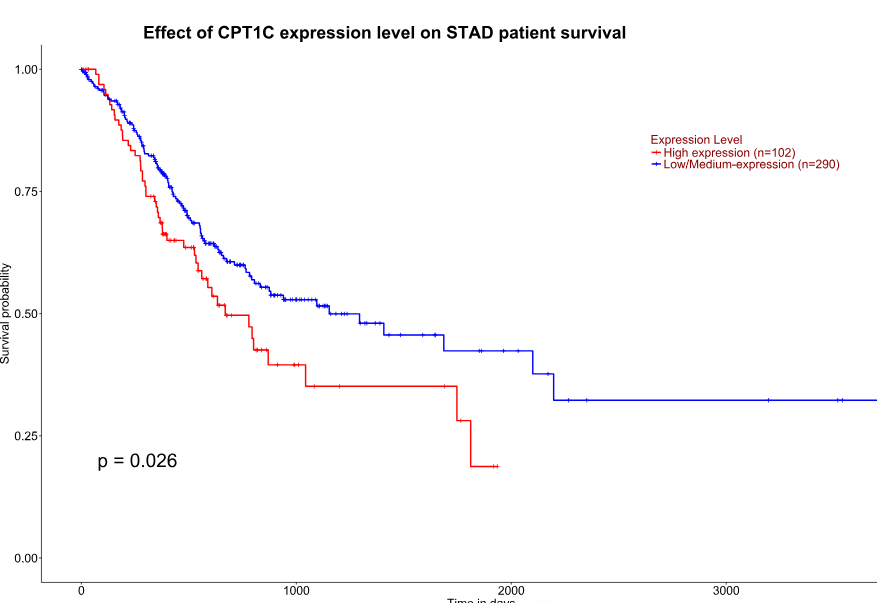

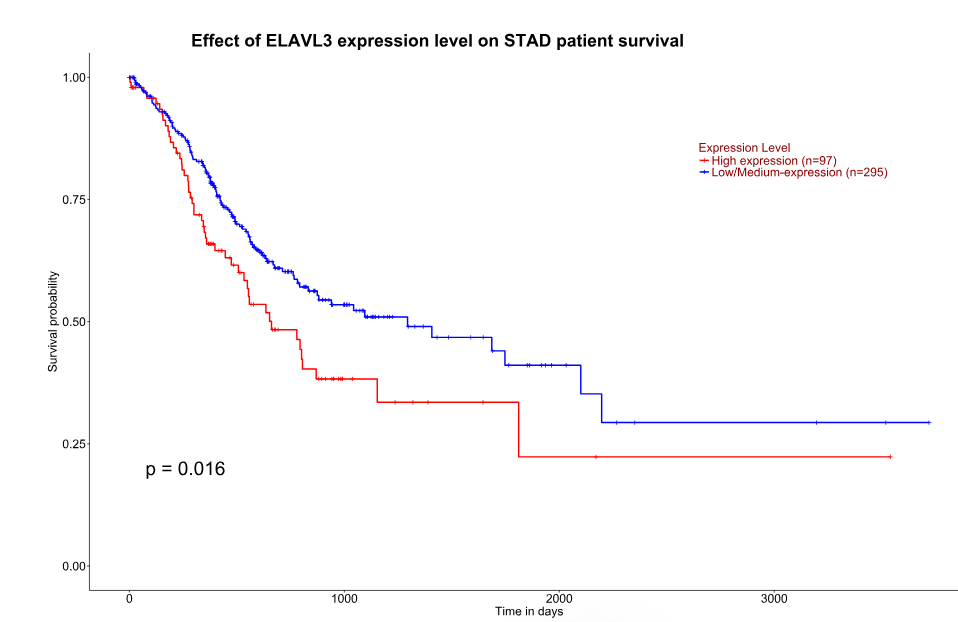

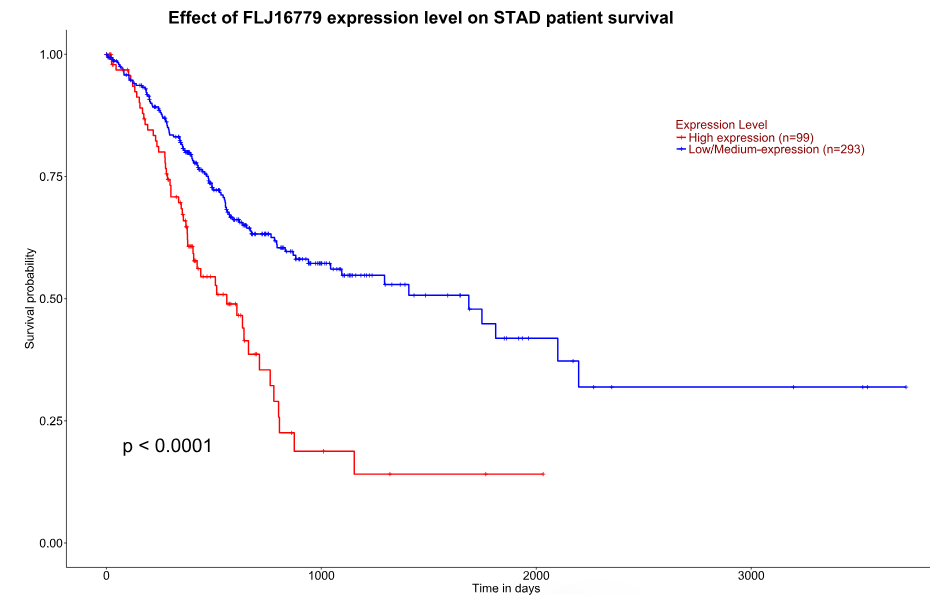

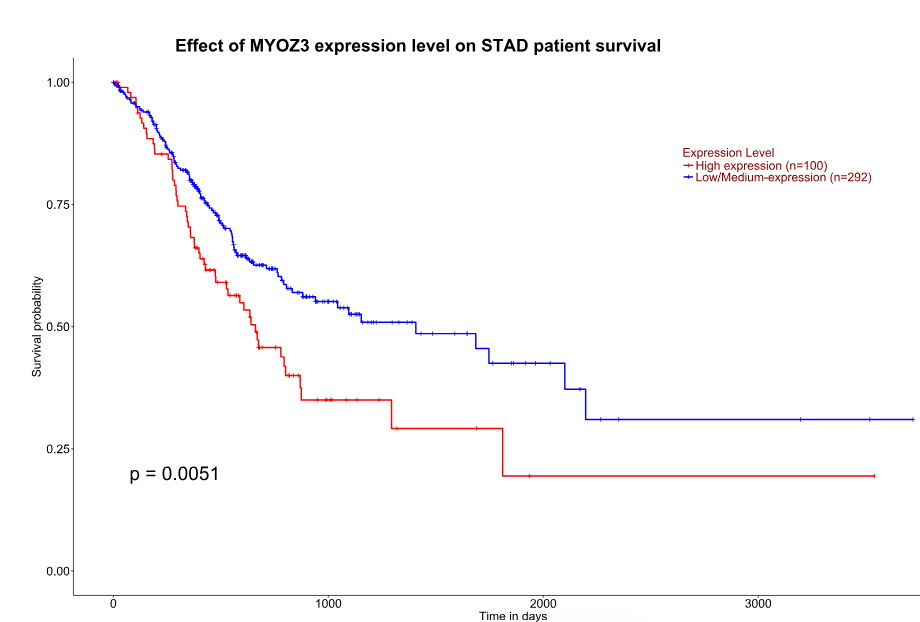

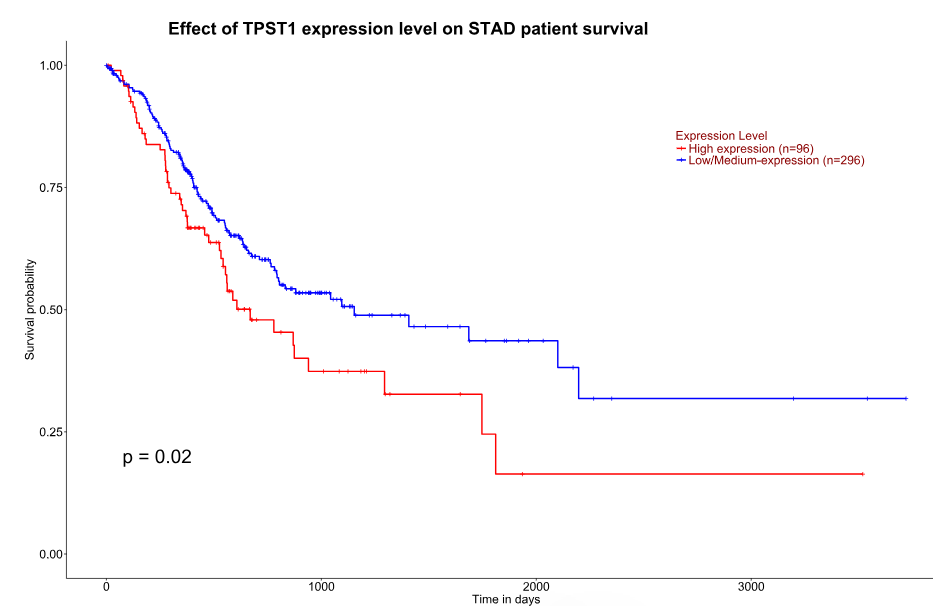

Figure S3: Shows high expression of the 6 of the 10 novel genes significantly associated with poor prognosis of stomach adenocarcinoma using UALCAN.

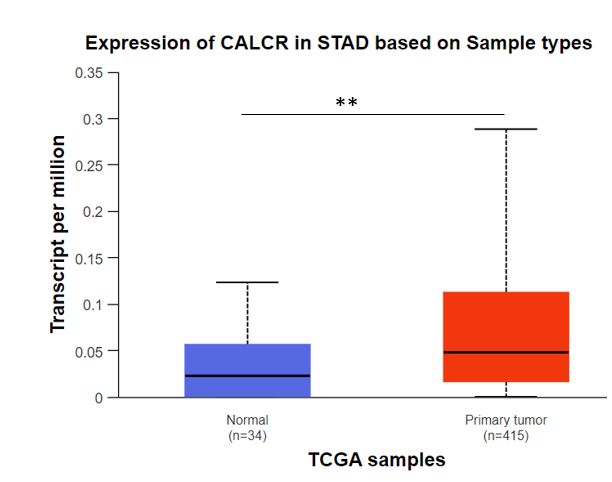

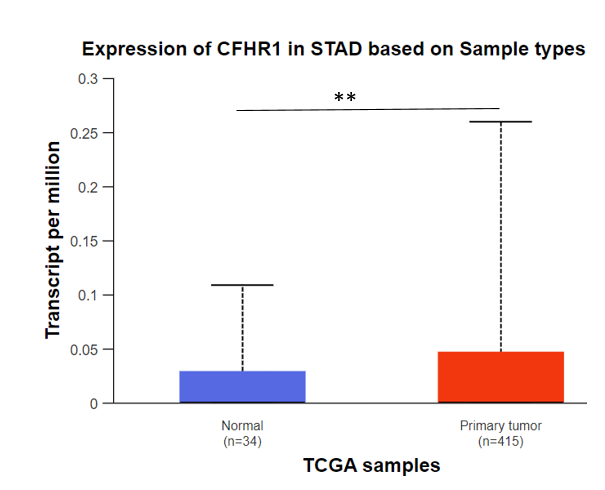

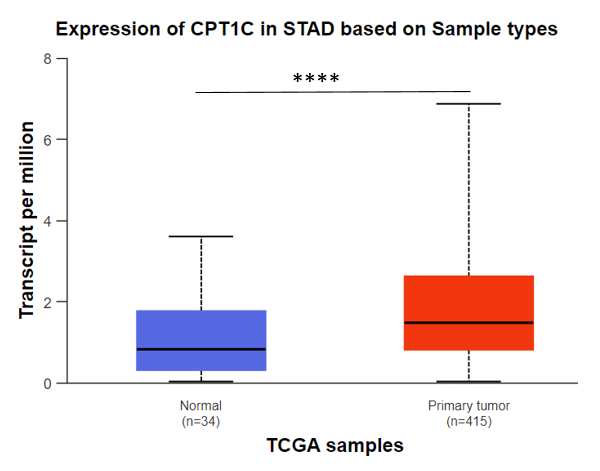

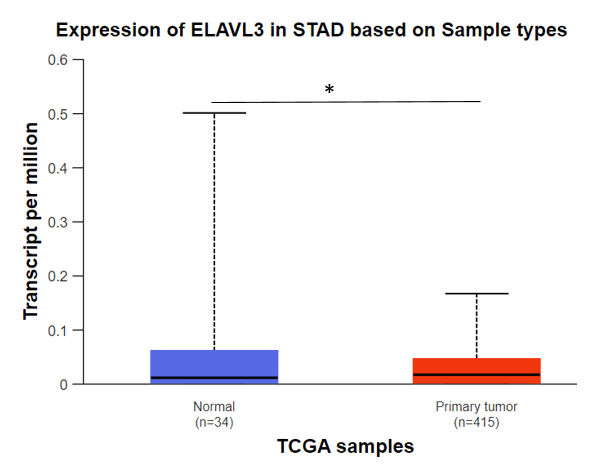

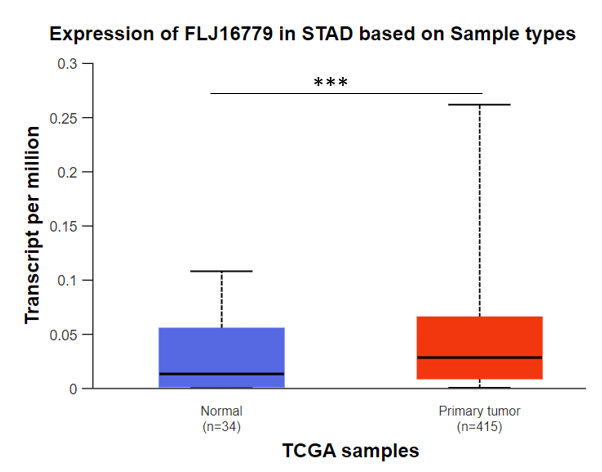

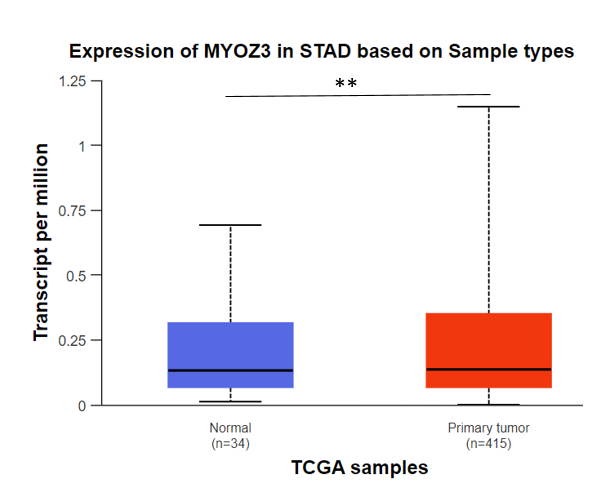

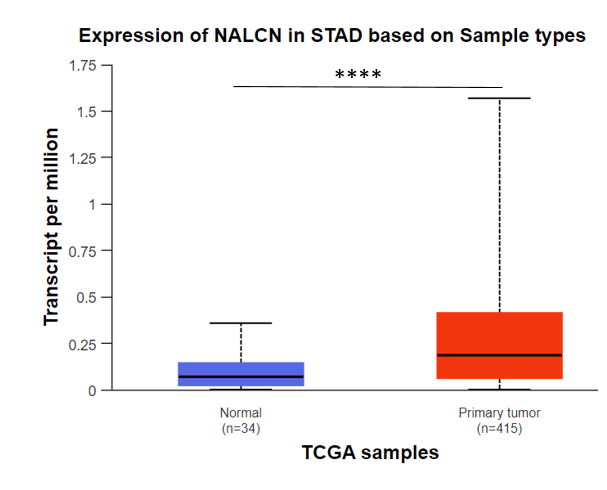

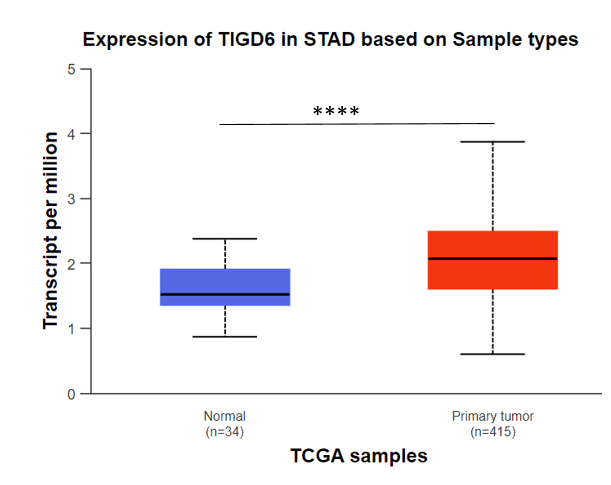

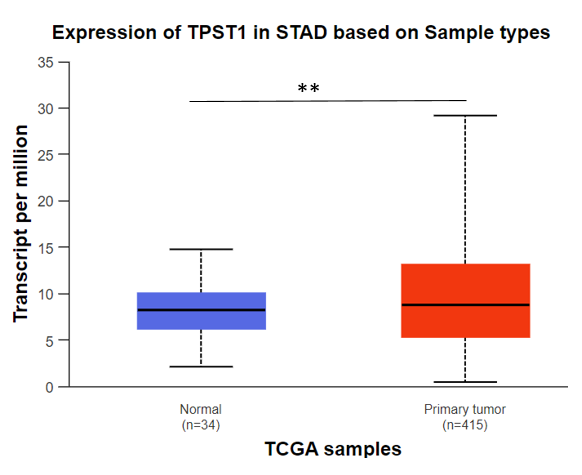

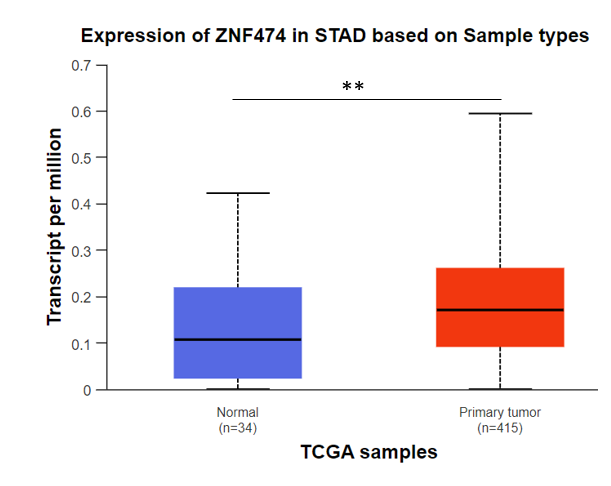

Figure S4: The mRNA expression levels of the 10 novel Prognosis-Associated Genes, shows significant increase in tumor tissues compared to normal controls determined by UALCAN database.

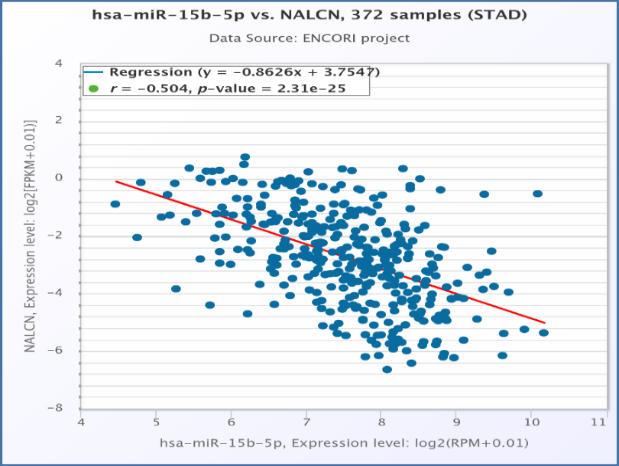

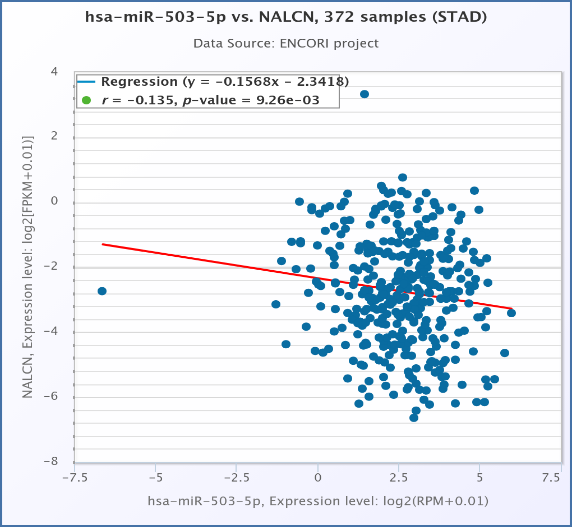

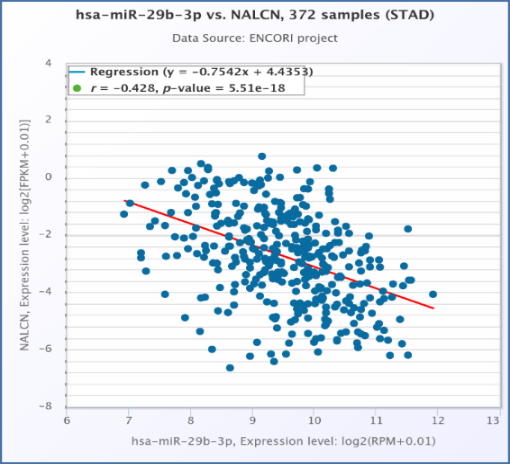

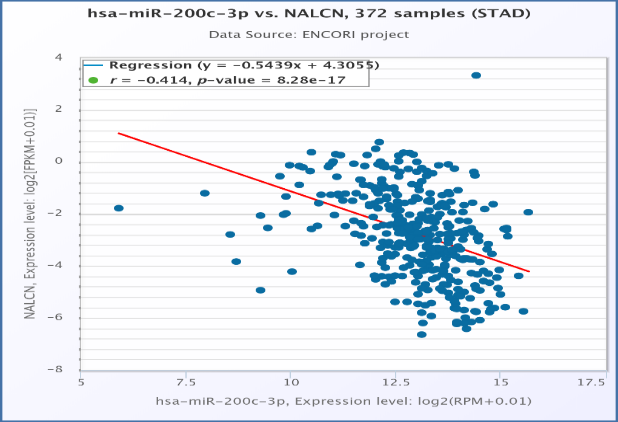

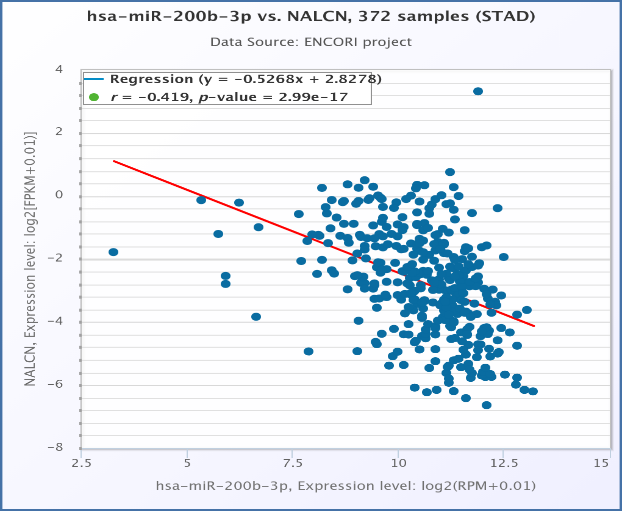

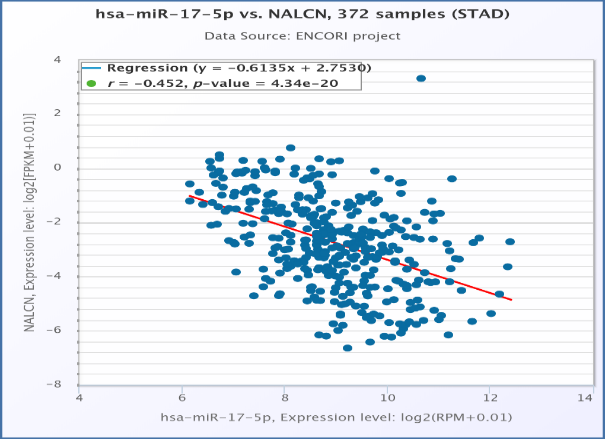

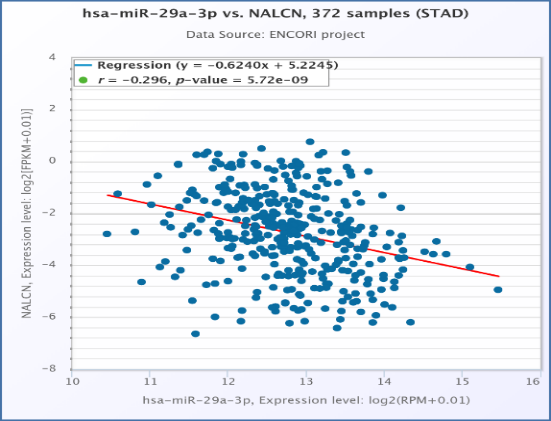

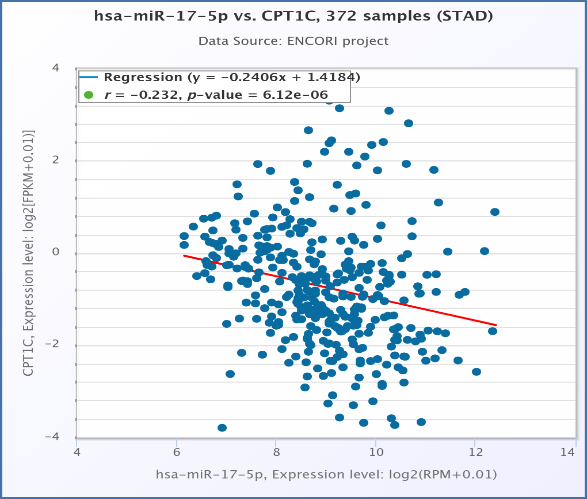

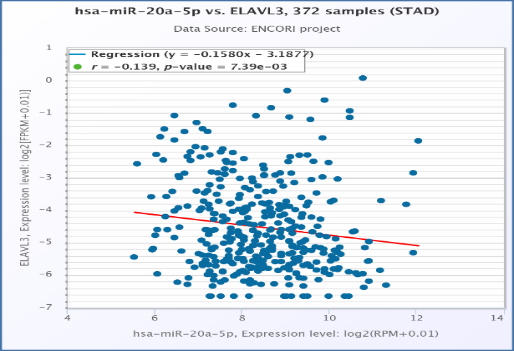

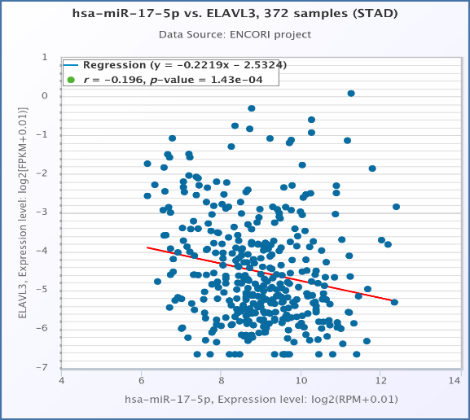

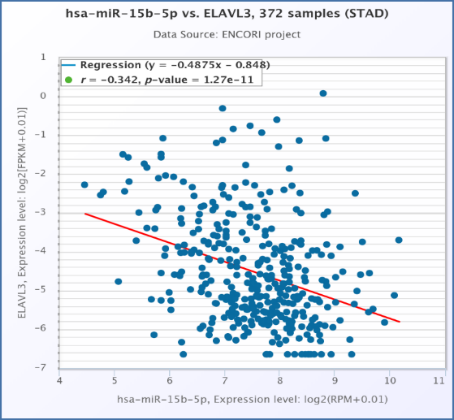

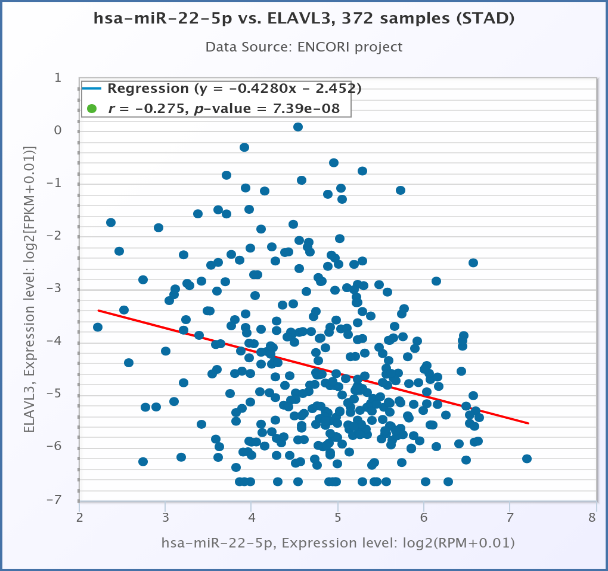

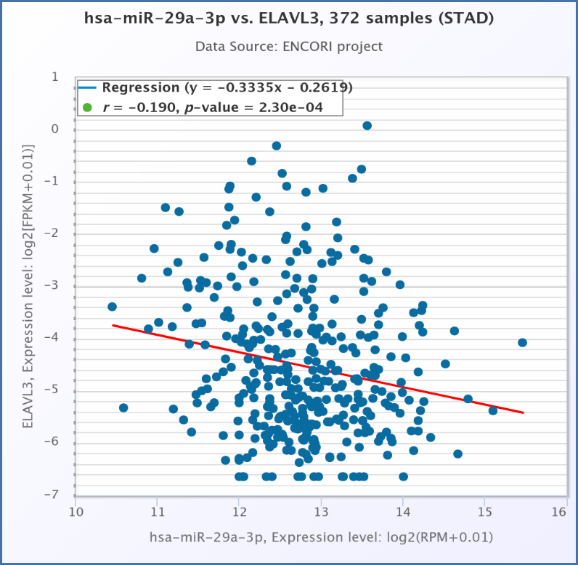

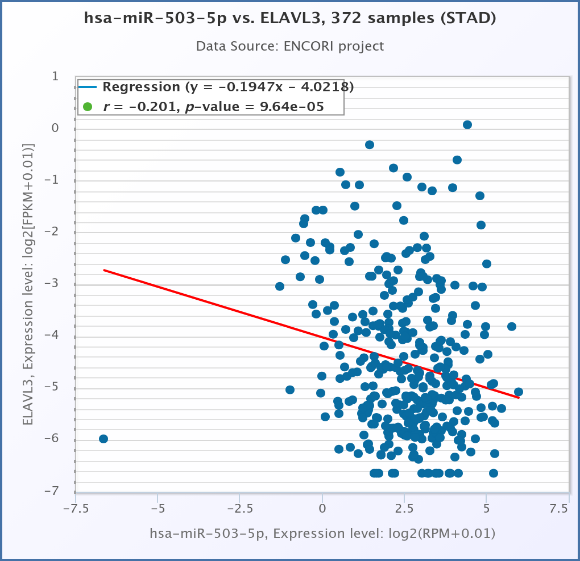

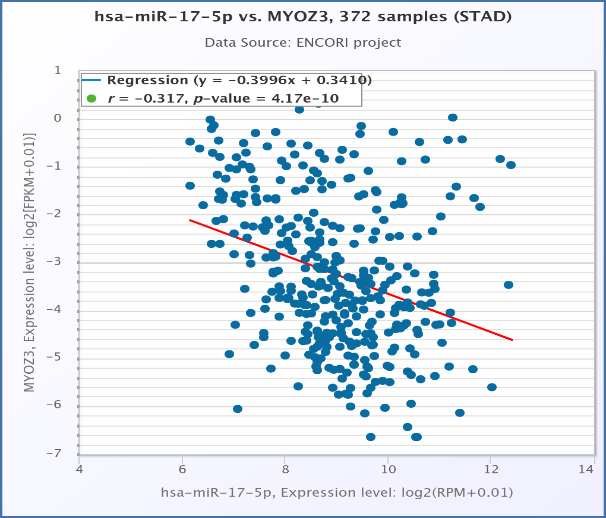

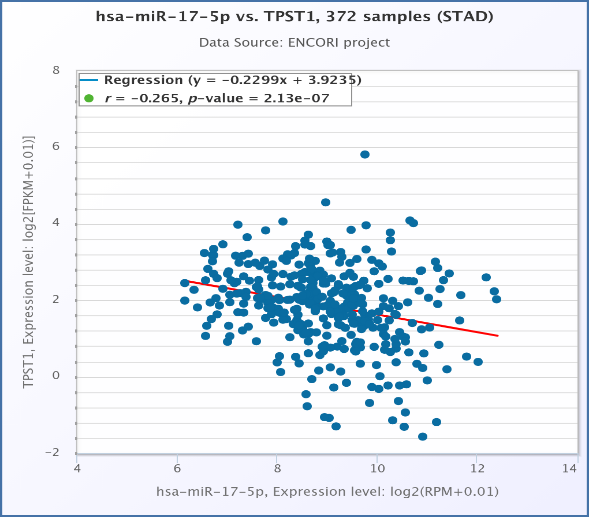

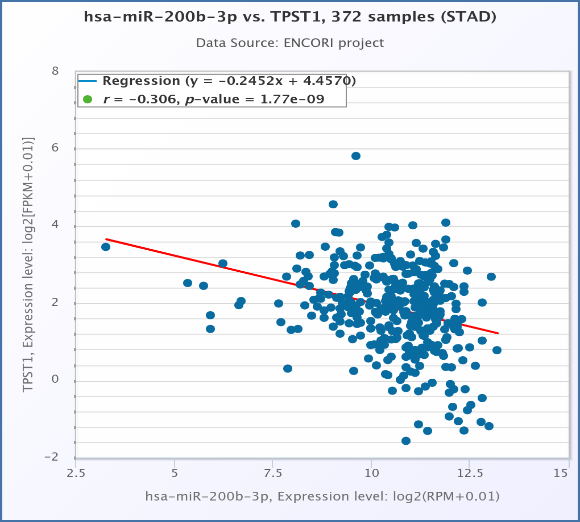

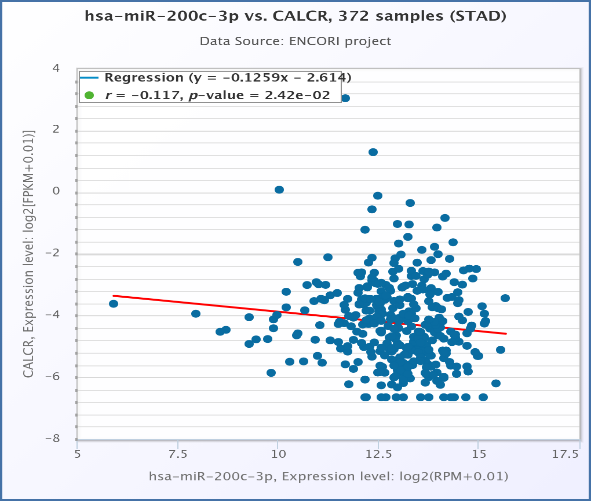

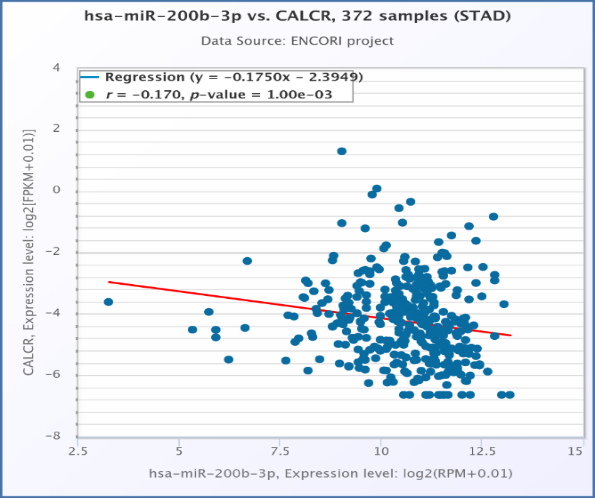

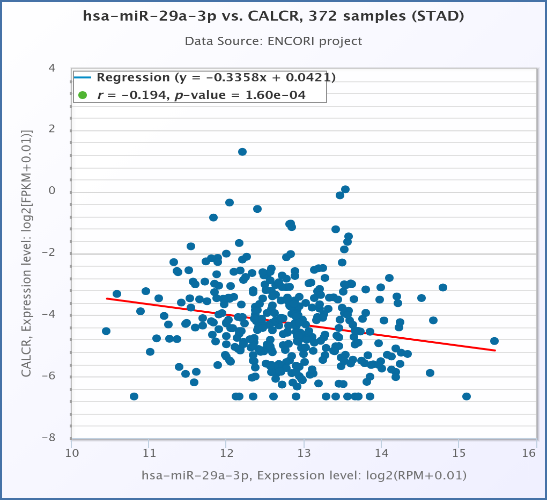

Figure S5: Expression of NALCN mRNA was significantly negatively associated with hsa-mir-15b-5p(A), hsa-mir-503-5p(B), hsa-mir-29b-3p(C), hsa-mir-200c-3p(D), hsa-mir-17-5p(E), hsa-mir-29a-3p(F), hsa-mir-200b-3p (G) and hsa-mir-20a-5p(H) expression. Expression of CPT1C mRNA was significantly negatively associated with hsa-mir-17-5P(I). Expression of ELAVL3 mRNA was significant negative expression with hsa-mir-20a-5p(J), hsa-mir-17-5p(K), hsa-mir-15b-5p(L), hsa-mir-22-3p(M), hsa-mir-29a-3p(N), and hsa-mir-503-5p(O). Expression of MYOZ3 mRNA was significantly negatively associated with hsa-mir-17-5p(P). hsa-mir-17-5p(Q) and hsa-mir-29c-3p(R) were negatively associated with TPST1mRNA expression. The CALCR mRNA was significantly negatively associated with hsa-mir-200c-3p(S), hsa-mir-200b-3p (T), and hsa-mir- 29a-3p(U).

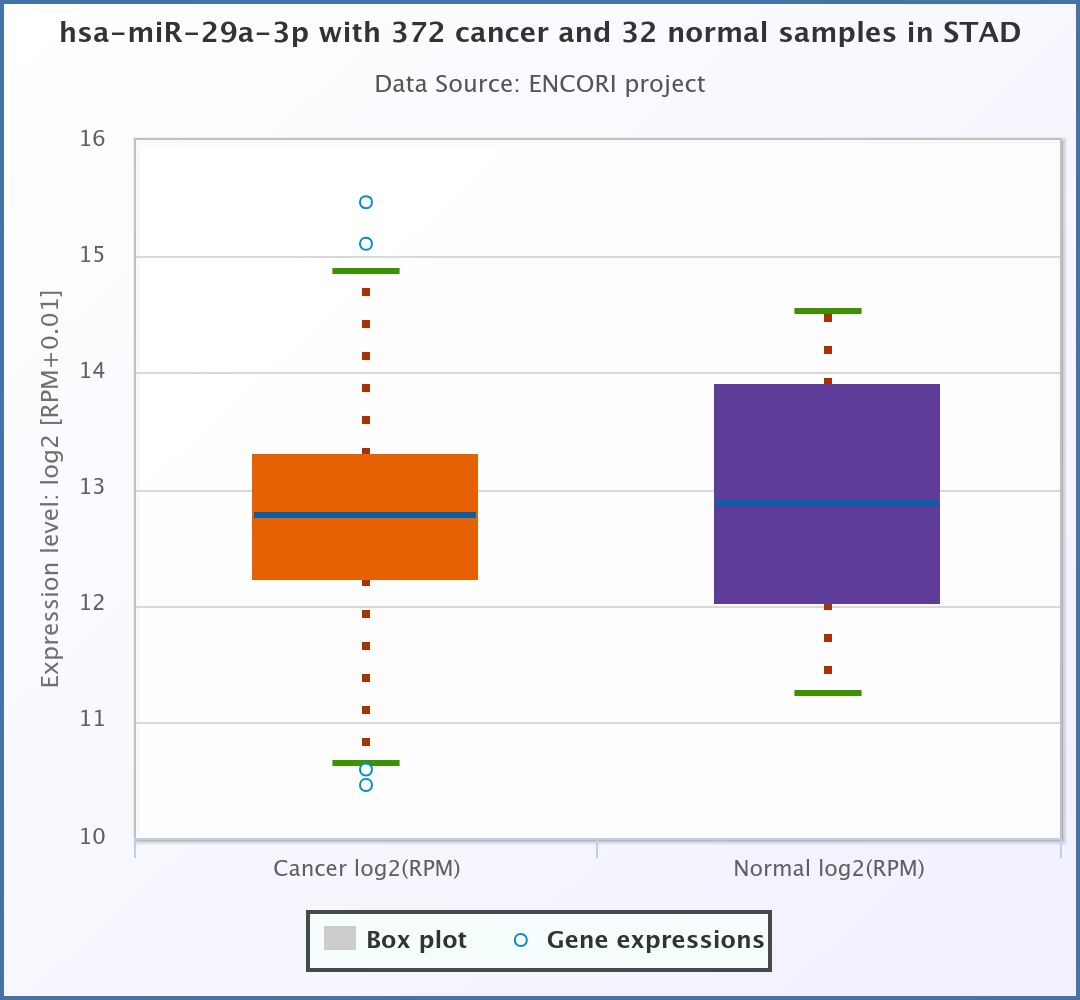

Figure S6. The significant miRNA with low expression in STAD.
