## supplementary tables for "A Bioinformatics-Driven ceRNA Network in Stomach Adenocarcinoma: Identification of Novel Prognostic mRNA-miRNA-lncRNA Interactions"

Table S1: The genes most associated with overall survival (OS) of patients with

Stomach adenocarcinoma determined by GEPIA database.

| Gene Symbol | Gene ID | P-Value (Survival os) |
| --- | --- | --- |
| GFAP | ENSG00000131095.11 | 2.12E-05 |
| RP11-497E19.1 | ENSG00000205562.2 | 2.33E-05 |
| ASPA | ENSG00000108381.10 | 2.69E-05 |
| SERPINE1 | ENSG00000106366.8 | 3.41E-05 |
| ZNF883 | ENSG00000228623.3 | 3.59E-05 |
| AOC4P | ENSG00000260105.6 | 3.86E-05 |
| CBLN4 | ENSG00000054803.3 | 3.97E-05 |
| NT5E | ENSG00000135318.11 | 4.74E-05 |
| AC002480.3 | ENSG00000232759.1 | 6.37E-05 |
| MEI4 | ENSG00000269964.2 | 6.39E-05 |
| CTD-2054N24.2 | ENSG00000259363.5 | 6.93E-05 |
| RP11-1069G10.2 | ENSG00000259727.1 | 9.09E-05 |
| EMX2OS | ENSG00000229847.8 | 9.24E-05 |
| ZNF192P1 | ENSG00000226314.7 | 9.45E-05 |
| RAI14 | ENSG00000039560.13 | 9.67E-05 |
| TRHDE-AS1 | ENSG00000236333.3 | 1.07E-04 |
| GUCY1A2 | ENSG00000152402.10 | 1.10E-04 |
| PCDHB17P | ENSG00000255622.3 | 1.12E-04 |
| PLCXD3 | ENSG00000182836.9 | 1.33E-04 |
| RP11-102K13.5 | ENSG00000278309.1 | 1.36E-04 |
| RP11-322E11.5 | ENSG00000267583.5 | 1.40E-04 |
| SORCS3 | ENSG00000156395.12 | 1.75E-04 |
| AKR1B1 | ENSG00000085662.13 | 1.80E-04 |
| CDR1 | ENSG00000184258.6 | 1.90E-04 |
| NRP1 | ENSG00000099250.17 | 2.03E-04 |
| PJA2 | ENSG00000198961.9 | 2.17E-04 |
| SLITRK2 | ENSG00000185985.8 | 2.32E-04 |
| INHBB | ENSG00000163083.5 | 2.34E-04 |
| LIFR-AS1 | ENSG00000244968.6 | 2.39E-04 |
| CPT1C | ENSG00000169169.14 | 2.65E-04 |
| GOLGA8IP | ENSG00000277561.4 | 2.92E-04 |
| SLC52A3 | ENSG00000101276.14 | 3.09E-04 |
| CPNE8 | ENSG00000139117.13 | 3.12E-04 |
| PYGO1 | ENSG00000171016.11 | 3.13E-04 |
| PKNOX2 | ENSG00000165495.15 | 3.29E-04 |
| TFPI2 | ENSG00000105825.11 | 3.39E-04 |
| HTR1F | ENSG00000179097.5 | 3.43E-04 |
| LINC00315 | ENSG00000184274.3 | 3.53E-04 |
| WDR49 | ENSG00000174776.10 | 3.88E-04 |
| SVEP1 | ENSG00000165124.17 | 3.92E-04 |
| ERBB4 | ENSG00000178568.13 | 3.93E-04 |
| DYNC1I1 | ENSG00000158560.14 | 3.98E-04 |
| ITIH3 | ENSG00000162267.12 | 4.19E-04 |
| ACOT1 | ENSG00000184227.7 | 4.27E-04 |
| MAGED4B | ENSG00000187243.16 | 4.47E-04 |
| FGF1 | ENSG00000113578.17 | 4.64E-04 |
| COLEC12 | ENSG00000158270.11 | 4.65E-04 |
| CCDC181 | ENSG00000117477.12 | 4.70E-04 |
| DGKQ | ENSG00000145214.13 | 4.72E-04 |
| EIF3EP1 | ENSG00000234882.1 | 4.84E-04 |
| ATP8A2 | ENSG00000132932.16 | 4.86E-04 |
| PPM1E | ENSG00000175175.5 | 4.86E-04 |
| CCDC178 | ENSG00000166960.16 | 4.92E-04 |
| AK5 | ENSG00000154027.18 | 4.97E-04 |
| PTPRQ | ENSG00000139304.12 | 4.98E-04 |
| MUM1L1 | ENSG00000157502.12 | 5.03E-04 |
| BLMH | ENSG00000108578.14 | 5.04E-04 |
| ANKRD6 | ENSG00000135299.16 | 5.54E-04 |
| PRTG | ENSG00000166450.12 | 5.68E-04 |
| PCDHB6 | ENSG00000113211.5 | 5.94E-04 |
| BASP1 | ENSG00000176788.8 | 5.95E-04 |
| ASTN1 | ENSG00000152092.15 | 6.06E-04 |
| RP11-469N6.1 | ENSG00000251226.1 | 6.24E-04 |
| C16orf47 | ENSG00000197445.2 | 6.28E-04 |
| VASH2 | ENSG00000143494.15 | 6.30E-04 |
| CTB-31O20.8 | ENSG00000267141.1 | 6.31E-04 |
| AC007228.11 | ENSG00000269696.1 | 6.33E-04 |
| CDC37 | ENSG00000105401.6 | 6.42E-04 |
| GPC3 | ENSG00000147257.13 | 6.44E-04 |
| P4HA3 | ENSG00000149380.11 | 6.45E-04 |
| CASC10 | ENSG00000204682.5 | 6.51E-04 |
| TCEAL7 | ENSG00000182916.7 | 6.90E-04 |
| PDK4 | ENSG00000004799.7 | 7.06E-04 |
| SGCE | ENSG00000127990.15 | 7.26E-04 |
| MCC | ENSG00000171444.17 | 7.32E-04 |
| MATN3 | ENSG00000132031.12 | 7.44E-04 |
| RP11-316O14.1 | ENSG00000268603.1 | 7.48E-04 |
| ABCB5 | ENSG00000004846.16 | 7.49E-04 |
| ZNF662 | ENSG00000182983.14 | 7.51E-04 |
| COL4A5 | ENSG00000188153.12 | 7.66E-04 |
| LRFN5 | ENSG00000165379.13 | 7.73E-04 |
| RP11-462L8.1 | ENSG00000229656.6 | 7.89E-04 |
| IQCA1 | ENSG00000132321.16 | 7.95E-04 |
| DPP3 | ENSG00000254986.7 | 7.96E-04 |
| RP11-310P5.1 | ENSG00000249650.1 | 8.09E-04 |
| DSCAM | ENSG00000171587.14 | 8.13E-04 |
| NPY1R | ENSG00000164128.6 | 8.14E-04 |
| EGFLAM | ENSG00000164318.17 | 8.18E-04 |
| FABP4 | ENSG00000170323.8 | 8.20E-04 |
| PCA3 | ENSG00000225937.1 | 8.27E-04 |
| ANKRD53 | ENSG00000144031.11 | 8.36E-04 |
| ADAMTS18 | ENSG00000140873.15 | 8.47E-04 |
| STK32A | ENSG00000169302.14 | 8.87E-04 |
| CSRNP1 | ENSG00000144655.14 | 9.08E-04 |
| ABCA8 | ENSG00000141338.13 | 9.09E-04 |
| CLRN3 | ENSG00000180745.4 | 9.16E-04 |
| NOVA1 | ENSG00000139910.19 | 9.38E-04 |
| AC005754.8 | ENSG00000272108.1 | 9.43E-04 |
| LINC01436 | ENSG00000231106.2 | 9.50E-04 |
| COMMD10 | ENSG00000145781.8 | 9.50E-04 |
| PCDHB5 | ENSG00000113209.8 | 9.71E-04 |
| PRR15L | ENSG00000167183.2 | 9.75E-04 |
| ARMCX1 | ENSG00000126947.11 | 9.77E-04 |
| PCOLCE2 | ENSG00000163710.7 | 9.82E-04 |
| NBAT1 | ENSG00000260455.1 | 9.87E-04 |
| ABCA4 | ENSG00000198691.11 | 1.01E-03 |
| SYN2 | ENSG00000157152.16 | 1.01E-03 |
| NAP1L2 | ENSG00000186462.8 | 1.04E-03 |
| SYPL2 | ENSG00000143028.8 | 1.04E-03 |
| RP11-566K19.6 | ENSG00000274253.4 | 1.04E-03 |
| LINC01606 | ENSG00000253301.5 | 1.10E-03 |
| SPX | ENSG00000134548.9 | 1.10E-03 |
| AC092667.2 | ENSG00000230393.1 | 1.11E-03 |
| CREB5 | ENSG00000146592.16 | 1.11E-03 |
| PGM5P3-AS1 | ENSG00000277631.4 | 1.13E-03 |
| C1GALT1C1L | ENSG00000223658.6 | 1.14E-03 |
| KCNT2 | ENSG00000162687.16 | 1.14E-03 |
| EMX2 | ENSG00000170370.11 | 1.15E-03 |
| GPNMB | ENSG00000136235.15 | 1.15E-03 |
| CTD-2525P14.5 | ENSG00000280073.1 | 1.17E-03 |
| RP11-295P9.8 | ENSG00000268549.1 | 1.19E-03 |
| AC005682.6 | ENSG00000226329.2 | 1.19E-03 |
| CD300LG | ENSG00000161649.12 | 1.20E-03 |
| ANXA8 | ENSG00000265190.6 | 1.21E-03 |
| FREM1 | ENSG00000164946.19 | 1.21E-03 |
| TCEAL5 | ENSG00000204065.2 | 1.27E-03 |
| APOD | ENSG00000189058.8 | 1.30E-03 |
| TAL2 | ENSG00000186051.6 | 1.32E-03 |
| NLGN4X | ENSG00000146938.14 | 1.32E-03 |
| GULP1 | ENSG00000144366.15 | 1.33E-03 |
| TRPC6 | ENSG00000137672.12 | 1.34E-03 |
| HOXD1 | ENSG00000128645.12 | 1.35E-03 |
| RP11-731F5.1 | ENSG00000254140.1 | 1.42E-03 |
| CRTAC1 | ENSG00000095713.13 | 1.44E-03 |
| PCDH9 | ENSG00000184226.14 | 1.45E-03 |
| PCDHB4 | ENSG00000081818.3 | 1.47E-03 |
| CDH6 | ENSG00000113361.12 | 1.49E-03 |
| PZP | ENSG00000126838.9 | 1.50E-03 |
| VCAN | ENSG00000038427.15 | 1.50E-03 |
| RERG | ENSG00000134533.6 | 1.54E-03 |
| NALCN | ENSG00000102452.15 | 1.54E-03 |
| STEAP4 | ENSG00000127954.12 | 1.54E-03 |
| CNRIP1 | ENSG00000119865.8 | 1.54E-03 |
| POU6F2 | ENSG00000106536.19 | 1.56E-03 |
| RP11-60A14.1 | ENSG00000279778.1 | 1.57E-03 |
| LRCOL1 | ENSG00000204583.9 | 1.58E-03 |
| RP3-425C14.4 | ENSG00000279453.1 | 1.58E-03 |
| KCND2 | ENSG00000184408.9 | 1.59E-03 |
| SYT14 | ENSG00000143469.16 | 1.59E-03 |
| AC026904.1 | ENSG00000233858.4 | 1.60E-03 |
| RARB | ENSG00000077092.18 | 1.61E-03 |
| CYP27C1 | ENSG00000186684.12 | 1.62E-03 |
| RP11-456K23.1 | ENSG00000267414.1 | 1.63E-03 |
| CALCR | ENSG00000004948.13 | 1.65E-03 |
| DNM1P46 | ENSG00000182397.14 | 1.65E-03 |
| SLC35F1 | ENSG00000196376.10 | 1.70E-03 |
| C1orf95 | ENSG00000203685.9 | 1.71E-03 |
| LINC01537 | ENSG00000227467.3 | 1.73E-03 |
| CADM2 | ENSG00000175161.13 | 1.73E-03 |
| AP000892.6 | ENSG00000280143.1 | 1.74E-03 |
| ITPRIPL1 | ENSG00000198885.9 | 1.74E-03 |
| HOXA10-AS | ENSG00000253187.2 | 1.77E-03 |
| ACSM5 | ENSG00000183549.10 | 1.77E-03 |
| RBMS1 | ENSG00000153250.17 | 1.81E-03 |
| NOXO1 | ENSG00000196408.11 | 1.81E-03 |
| CPA6 | ENSG00000165078.11 | 1.83E-03 |
| SLC2A3 | ENSG00000059804.15 | 1.85E-03 |
| RAB34 | ENSG00000109113.17 | 1.85E-03 |
| FAM20A | ENSG00000108950.11 | 1.86E-03 |
| RASSF8 | ENSG00000123094.15 | 1.87E-03 |
| RP11-567J20.2 | ENSG00000253688.1 | 1.87E-03 |
| ZFHX4 | ENSG00000091656.15 | 1.88E-03 |
| MPP4 | ENSG00000082126.17 | 1.89E-03 |
| EIF2AK4 | ENSG00000128829.11 | 1.89E-03 |
| ZNF22 | ENSG00000165512.4 | 1.92E-03 |
| PDLIM1P4 | ENSG00000249274.1 | 1.92E-03 |
| MFAP3L | ENSG00000198948.11 | 1.92E-03 |
| SRMS | ENSG00000125508.3 | 1.94E-03 |
| ZNF415P1 | ENSG00000266127.1 | 1.95E-03 |
| TSPAN5 | ENSG00000168785.7 | 1.96E-03 |
| ASF1B | ENSG00000105011.8 | 1.96E-03 |
| AC091878.1 | ENSG00000215196.4 | 1.97E-03 |
| RP11-211N11.5 | ENSG00000234393.1 | 1.98E-03 |
| NAT8L | ENSG00000185818.7 | 1.98E-03 |
| NECAB1 | ENSG00000123119.11 | 1.99E-03 |
| ZNF331 | ENSG00000130844.16 | 2.00E-03 |
| FOLR3 | ENSG00000110203.8 | 2.01E-03 |
| EHD3 | ENSG00000013016.14 | 2.01E-03 |
| PTCHD4 | ENSG00000244694.7 | 2.02E-03 |
| BOLA3-AS1 | ENSG00000225439.2 | 2.03E-03 |
| SEC23A | ENSG00000100934.14 | 2.04E-03 |
| IL1RAPL1 | ENSG00000169306.9 | 2.08E-03 |
| BEX4 | ENSG00000102409.9 | 2.09E-03 |
| AKAP12 | ENSG00000131016.16 | 2.09E-03 |
| NRG3 | ENSG00000185737.12 | 2.10E-03 |
| LINC00961 | ENSG00000235387.1 | 2.11E-03 |
| CLCN3P1 | ENSG00000232000.2 | 2.11E-03 |
| CEP83-AS1 | ENSG00000278916.1 | 2.11E-03 |
| CHAF1A | ENSG00000167670.15 | 2.11E-03 |
| SCHIP1 | ENSG00000151967.18 | 2.11E-03 |
| OSBPL1A | ENSG00000141447.16 | 2.14E-03 |
| CHST14 | ENSG00000169105.7 | 2.14E-03 |
| SLC16A7 | ENSG00000118596.11 | 2.17E-03 |
| LINC00989 | ENSG00000250334.5 | 2.28E-03 |
| ACOT2 | ENSG00000119673.14 | 2.30E-03 |
| RP11-620J15.3 | ENSG00000257698.1 | 2.31E-03 |
| EPDR1 | ENSG00000086289.11 | 2.32E-03 |
| RP11-624L4.1 | ENSG00000259345.5 | 2.32E-03 |
| PRL | ENSG00000172179.11 | 2.36E-03 |
| NTN1 | ENSG00000065320.8 | 2.37E-03 |
| TCF7L1 | ENSG00000152284.4 | 2.38E-03 |
| ITGAV | ENSG00000138448.11 | 2.41E-03 |
| RP11-59D5__B.2 | ENSG00000236345.1 | 2.41E-03 |
| ZBTB10 | ENSG00000205189.11 | 2.41E-03 |
| GALNT13 | ENSG00000144278.14 | 2.43E-03 |
| CDK15 | ENSG00000138395.14 | 2.44E-03 |
| MIR99AHG | ENSG00000215386.10 | 2.46E-03 |
| RP11-81K2.2 | ENSG00000279036.1 | 2.47E-03 |
| RP11-64C12.8 | ENSG00000267069.1 | 2.49E-03 |
| LINC00648 | ENSG00000259129.5 | 2.51E-03 |
| LINC00968 | ENSG00000246430.6 | 2.52E-03 |
| GLP2R | ENSG00000065325.12 | 2.52E-03 |
| WHAMMP3 | ENSG00000276141.4 | 2.53E-03 |
| LGR6 | ENSG00000133067.17 | 2.54E-03 |
| NTAN1 | ENSG00000157045.8 | 2.56E-03 |
| FAM153B | ENSG00000182230.11 | 2.58E-03 |
| PDE7B | ENSG00000171408.13 | 2.60E-03 |
| ITGA5 | ENSG00000161638.10 | 2.61E-03 |
| FAM133A | ENSG00000179083.6 | 2.61E-03 |
| STC1 | ENSG00000159167.11 | 2.63E-03 |
| PHF24 | ENSG00000122733.12 | 2.63E-03 |
| MSC-AS1 | ENSG00000235531.9 | 2.68E-03 |
| RP11-61I13.3 | ENSG00000235033.7 | 2.68E-03 |
| NPAS3 | ENSG00000151322.18 | 2.68E-03 |
| ADRA1D | ENSG00000171873.7 | 2.68E-03 |
| HAVCR1 | ENSG00000113249.12 | 2.68E-03 |
| AC011239.2 | ENSG00000279526.1 | 2.70E-03 |
| ADGRL4 | ENSG00000162618.12 | 2.74E-03 |
| GOLGA8T | ENSG00000261247.1 | 2.75E-03 |
| ZNF208 | ENSG00000160321.14 | 2.76E-03 |
| GPX3 | ENSG00000211445.11 | 2.78E-03 |
| LAMA4 | ENSG00000112769.18 | 2.79E-03 |
| CCDC23 | ENSG00000177868.11 | 2.82E-03 |
| AC020571.3 | ENSG00000229056.2 | 2.84E-03 |
| RP11-557L19.1 | ENSG00000272002.1 | 2.84E-03 |
| CACNB4 | ENSG00000182389.18 | 2.84E-03 |
| KIAA1755 | ENSG00000149633.11 | 2.87E-03 |
| RP11-513O13.1 | ENSG00000279881.1 | 2.89E-03 |
| ATP2B3 | ENSG00000067842.17 | 2.89E-03 |
| LOX | ENSG00000113083.12 | 2.89E-03 |
| ST8SIA6 | ENSG00000148488.15 | 2.90E-03 |
| RP11-180I4.4 | ENSG00000268926.2 | 2.93E-03 |
| RP11-876N24.5 | ENSG00000263013.1 | 2.94E-03 |
| TMEM45A | ENSG00000181458.10 | 2.95E-03 |
| HIF3A | ENSG00000124440.15 | 2.97E-03 |
| TUBA3D | ENSG00000075886.10 | 2.98E-03 |
| CCDC42B | ENSG00000186710.11 | 2.99E-03 |
| ELAVL3 | ENSG00000196361.9 | 3.01E-03 |
| SHC4 | ENSG00000185634.11 | 3.02E-03 |
| ZNF257 | ENSG00000197134.11 | 3.03E-03 |
| MAGED4 | ENSG00000154545.16 | 3.03E-03 |
| RASSF8-AS1 | ENSG00000246695.7 | 3.06E-03 |
| CELF4 | ENSG00000101489.18 | 3.07E-03 |
| AFF3 | ENSG00000144218.18 | 3.09E-03 |
| GRTP1 | ENSG00000139835.13 | 3.10E-03 |
| BCHE | ENSG00000114200.9 | 3.14E-03 |
| TPST1 | ENSG00000169902.13 | 3.15E-03 |
| TIGD6 | ENSG00000164296.6 | 3.16E-03 |
| WNK3 | ENSG00000196632.10 | 3.18E-03 |
| C6orf48 | ENSG00000204387.12 | 3.18E-03 |
| CPNE6 | ENSG00000100884.9 | 3.25E-03 |
| TET1 | ENSG00000138336.8 | 3.25E-03 |
| MUSK | ENSG00000030304.12 | 3.26E-03 |
| PER1 | ENSG00000179094.13 | 3.27E-03 |
| OGN | ENSG00000106809.10 | 3.29E-03 |
| FBXO17 | ENSG00000269190.5 | 3.30E-03 |
| RP11-401O9.4 | ENSG00000273388.1 | 3.30E-03 |
| AP000476.1 | ENSG00000237484.5 | 3.32E-03 |
| LINC00310 | ENSG00000227456.7 | 3.33E-03 |
| GFRA2 | ENSG00000168546.10 | 3.35E-03 |
| NDUFAF3 | ENSG00000178057.14 | 3.35E-03 |
| CTD-2536I1.2 | ENSG00000280304.1 | 3.39E-03 |
| MOSPD1 | ENSG00000101928.12 | 3.45E-03 |
| PLAT | ENSG00000104368.17 | 3.46E-03 |
| LINC00619 | ENSG00000204187.5 | 3.46E-03 |
| C19orf26 | ENSG00000099625.12 | 3.49E-03 |
| IGFBP7-AS1 | ENSG00000245067.6 | 3.52E-03 |
| ZNF300 | ENSG00000145908.12 | 3.54E-03 |
| RP11-553A21.3 | ENSG00000231652.2 | 3.56E-03 |
| MDP1 | ENSG00000213920.8 | 3.58E-03 |
| NKAIN2 | ENSG00000188580.13 | 3.59E-03 |
| ARAP1-AS2 | ENSG00000245148.2 | 3.59E-03 |
| MOGAT1 | ENSG00000124003.12 | 3.60E-03 |
| AL592528.1 | ENSG00000205424.1 | 3.64E-03 |
| EFNA3 | ENSG00000143590.13 | 3.65E-03 |
| GPR173 | ENSG00000184194.5 | 3.68E-03 |
| GABARAPL2 | ENSG00000034713.7 | 3.70E-03 |
| G0S2 | ENSG00000123689.5 | 3.70E-03 |
| SV2B | ENSG00000185518.11 | 3.73E-03 |
| RGS4 | ENSG00000117152.13 | 3.75E-03 |
| KCNIP1 | ENSG00000182132.12 | 3.77E-03 |
| RAMP1 | ENSG00000132329.10 | 3.78E-03 |
| THRB | ENSG00000151090.17 | 3.84E-03 |
| AP001626.2 | ENSG00000235023.1 | 3.86E-03 |
| AFAP1L1 | ENSG00000157510.13 | 3.86E-03 |
| MAGI2-AS3 | ENSG00000234456.7 | 3.86E-03 |
| AC016995.3 | ENSG00000231367.5 | 3.88E-03 |
| LRAT | ENSG00000121207.11 | 3.88E-03 |
| RP11-681H18.2 | ENSG00000277801.1 | 3.90E-03 |
| RP13-514E23.1 | ENSG00000261496.1 | 3.93E-03 |
| PCDHB16 | ENSG00000272674.3 | 3.95E-03 |
| DOCK4 | ENSG00000128512.19 | 3.97E-03 |
| CDO1 | ENSG00000129596.4 | 3.99E-03 |
| RP6-109B7.5 | ENSG00000273289.1 | 3.99E-03 |
| KIAA1324L | ENSG00000164659.14 | 3.99E-03 |
| USP51 | ENSG00000247746.4 | 3.99E-03 |
| CTD-2008L17.2 | ENSG00000206129.3 | 4.00E-03 |
| AC004947.2 | ENSG00000233760.1 | 4.03E-03 |
| DZIP1 | ENSG00000134874.17 | 4.05E-03 |
| TCN2 | ENSG00000185339.8 | 4.06E-03 |
| RP11-713P17.3 | ENSG00000204241.7 | 4.08E-03 |
| PTPN6 | ENSG00000111679.16 | 4.11E-03 |
| IGFBP7 | ENSG00000163453.11 | 4.11E-03 |
| AC114730.2 | ENSG00000235151.1 | 4.12E-03 |
| RP11-542B15.1 | ENSG00000203585.3 | 4.12E-03 |
| CCNA1 | ENSG00000133101.9 | 4.12E-03 |
| GPR156 | ENSG00000175697.10 | 4.12E-03 |
| RP11-632L2.2 | ENSG00000278177.1 | 4.15E-03 |
| MFGE8 | ENSG00000140545.14 | 4.18E-03 |
| IGHE | ENSG00000211891.5 | 4.20E-03 |
| RP11-108M12.3 | ENSG00000258592.1 | 4.22E-03 |
| VIPAS39 | ENSG00000151445.15 | 4.23E-03 |
| HGF | ENSG00000019991.15 | 4.25E-03 |
| SPIRE1 | ENSG00000134278.14 | 4.25E-03 |
| ZNF229 | ENSG00000278318.4 | 4.26E-03 |
| ZNF423 | ENSG00000102935.11 | 4.27E-03 |
| RP11-274H2.3 | ENSG00000240032.1 | 4.29E-03 |
| CERS5 | ENSG00000139624.12 | 4.30E-03 |
| NACAD | ENSG00000136274.8 | 4.31E-03 |
| BICC1 | ENSG00000122870.11 | 4.33E-03 |
| AP4S1 | ENSG00000100478.14 | 4.33E-03 |
| GTF2H2C | ENSG00000183474.15 | 4.34E-03 |
| PCDHB3 | ENSG00000113205.4 | 4.35E-03 |
| EYA1 | ENSG00000104313.17 | 4.38E-03 |
| RP11-600F24.1 | ENSG00000243904.1 | 4.38E-03 |
| THBS1 | ENSG00000137801.10 | 4.39E-03 |
| QKI | ENSG00000112531.16 | 4.43E-03 |
| PHKG1 | ENSG00000164776.9 | 4.45E-03 |
| CHRFAM7A | ENSG00000166664.13 | 4.48E-03 |
| MTTP | ENSG00000138823.12 | 4.50E-03 |
| CD109 | ENSG00000156535.13 | 4.52E-03 |
| PRSS3 | ENSG00000010438.16 | 4.53E-03 |
| DPT | ENSG00000143196.4 | 4.56E-03 |
| EDNRB | ENSG00000136160.14 | 4.58E-03 |
| KLHDC2 | ENSG00000165516.10 | 4.58E-03 |
| HAGLR | ENSG00000224189.6 | 4.60E-03 |
| MSRB3 | ENSG00000174099.10 | 4.61E-03 |
| RP11-398K22.12 | ENSG00000229852.2 | 4.62E-03 |
| ARMCX2 | ENSG00000184867.13 | 4.62E-03 |
| NAP1L6 | ENSG00000204118.1 | 4.63E-03 |
| CSMD2 | ENSG00000121904.17 | 4.64E-03 |
| DIRC1 | ENSG00000174325.4 | 4.66E-03 |
| TREML4 | ENSG00000188056.11 | 4.69E-03 |
| ACVR1 | ENSG00000115170.13 | 4.69E-03 |
| APBB1 | ENSG00000166313.18 | 4.71E-03 |
| ANO4 | ENSG00000151572.16 | 4.72E-03 |
| COL24A1 | ENSG00000171502.14 | 4.72E-03 |
| C1QTNF4 | ENSG00000172247.3 | 4.73E-03 |
| CAND2 | ENSG00000144712.11 | 4.73E-03 |
| EEF1A2 | ENSG00000101210.10 | 4.74E-03 |
| AK4 | ENSG00000162433.14 | 4.75E-03 |
| WDR17 | ENSG00000150627.15 | 4.76E-03 |
| LINC00475 | ENSG00000225511.6 | 4.78E-03 |
| AC009948.5 | ENSG00000223960.6 | 4.79E-03 |
| RP11-155D18.13 | ENSG00000280422.1 | 4.79E-03 |
| CLIP3 | ENSG00000105270.14 | 4.80E-03 |
| PCDHB13 | ENSG00000187372.11 | 4.80E-03 |
| AC000403.4 | ENSG00000278727.1 | 4.82E-03 |
| RP3-388E23.2 | ENSG00000234084.1 | 4.82E-03 |
| NTMT1 | ENSG00000148335.14 | 4.85E-03 |
| ADRA1B | ENSG00000170214.3 | 4.85E-03 |
| RAB19 | ENSG00000146955.10 | 4.85E-03 |
| CRB1 | ENSG00000134376.14 | 4.85E-03 |
| ZNF474 | ENSG00000164185.4 | 4.87E-03 |
| GPR176 | ENSG00000166073.8 | 4.87E-03 |
| PSD4 | ENSG00000125637.15 | 4.87E-03 |
| LGALS12 | ENSG00000133317.14 | 4.88E-03 |
| RPL21P40 | ENSG00000235670.1 | 4.89E-03 |
| MYL3 | ENSG00000160808.9 | 4.92E-03 |
| NPTX1 | ENSG00000171246.5 | 4.95E-03 |
| LINC01140 | ENSG00000267272.5 | 4.98E-03 |
| NUDT10 | ENSG00000122824.10 | 5.01E-03 |
| POT1-AS1 | ENSG00000224897.6 | 5.01E-03 |
| AC002456.2 | ENSG00000223969.5 | 5.05E-03 |
| PCCA | ENSG00000175198.14 | 5.07E-03 |
| UBE2QL1 | ENSG00000215218.3 | 5.08E-03 |
| C6orf120 | ENSG00000185127.6 | 5.08E-03 |
| GLT8D1 | ENSG00000016864.16 | 5.10E-03 |
| DIMT1 | ENSG00000086189.9 | 5.10E-03 |
| CHRD | ENSG00000090539.15 | 5.10E-03 |
| ZFPM2 | ENSG00000169946.13 | 5.13E-03 |
| TGFB2 | ENSG00000092969.11 | 5.14E-03 |
| C5 | ENSG00000106804.7 | 5.16E-03 |
| DMRTC1B | ENSG00000184911.14 | 5.16E-03 |
| CAV1 | ENSG00000105974.11 | 5.17E-03 |
| TNFAIP8L3 | ENSG00000183578.5 | 5.19E-03 |
| HSPD1P11 | ENSG00000251348.1 | 5.19E-03 |
| EBF2 | ENSG00000221818.8 | 5.21E-03 |
| GJA1 | ENSG00000152661.7 | 5.21E-03 |
| C1QTNF2 | ENSG00000145861.7 | 5.22E-03 |
| PCDHA1 | ENSG00000204970.9 | 5.22E-03 |
| CTB-134H23.3 | ENSG00000260908.1 | 5.23E-03 |
| FLJ16779 | ENSG00000275620.1 | 5.23E-03 |
| LINC00922 | ENSG00000261742.5 | 5.24E-03 |
| ARGLU1 | ENSG00000134884.13 | 5.24E-03 |
| MASP1 | ENSG00000127241.16 | 5.25E-03 |
| SOCS2 | ENSG00000120833.13 | 5.29E-03 |
| SOX7 | ENSG00000171056.7 | 5.31E-03 |
| CITED2 | ENSG00000164442.9 | 5.31E-03 |
| NREP | ENSG00000134986.13 | 5.32E-03 |
| SLITRK4 | ENSG00000179542.15 | 5.34E-03 |
| RNF217 | ENSG00000146373.16 | 5.34E-03 |
| WASF1 | ENSG00000112290.12 | 5.36E-03 |
| SYT6 | ENSG00000134207.14 | 5.40E-03 |
| MMP19 | ENSG00000123342.15 | 5.41E-03 |
| PTPRD | ENSG00000153707.15 | 5.44E-03 |
| RP11-311D14.1 | ENSG00000248778.1 | 5.46E-03 |
| ILDR1 | ENSG00000145103.12 | 5.46E-03 |
| REEP4 | ENSG00000168476.11 | 5.49E-03 |
| RP4-545K15.5 | ENSG00000261101.2 | 5.49E-03 |
| ZRANB2-AS2 | ENSG00000229956.9 | 5.51E-03 |
| IL34 | ENSG00000157368.10 | 5.52E-03 |
| WBSCR17 | ENSG00000185274.11 | 5.52E-03 |
| SV2A | ENSG00000159164.9 | 5.52E-03 |
| KCNK2 | ENSG00000082482.13 | 5.58E-03 |
| SIAH3 | ENSG00000215475.4 | 5.60E-03 |
| RP11-114H24.2 | ENSG00000260776.5 | 5.61E-03 |
| FAM181B | ENSG00000182103.4 | 5.62E-03 |
| PPP1R14A | ENSG00000167641.10 | 5.64E-03 |
| NAMA | ENSG00000271086.5 | 5.66E-03 |
| IL1R1 | ENSG00000115594.11 | 5.69E-03 |
| CCNDBP1 | ENSG00000166946.13 | 5.69E-03 |
| RAB9B | ENSG00000123570.3 | 5.70E-03 |
| PDGFD | ENSG00000170962.12 | 5.71E-03 |
| ZC3H12C | ENSG00000149289.10 | 5.71E-03 |
| APOC3 | ENSG00000110245.11 | 5.71E-03 |
| RP11-284F21.7 | ENSG00000229953.1 | 5.72E-03 |
| UBE2Q2 | ENSG00000140367.11 | 5.75E-03 |
| STARD9 | ENSG00000159433.11 | 5.75E-03 |
| NT5C1A | ENSG00000116981.3 | 5.77E-03 |
| CFHR1 | ENSG00000244414.6 | 5.77E-03 |
| PROSER2-AS1 | ENSG00000225778.5 | 5.78E-03 |
| IL1RL1 | ENSG00000115602.16 | 5.78E-03 |
| RP11-594N15.3 | ENSG00000260398.1 | 5.82E-03 |
| BRINP1 | ENSG00000078725.12 | 5.83E-03 |
| CCNO | ENSG00000152669.8 | 5.83E-03 |
| SYT4 | ENSG00000132872.11 | 5.85E-03 |
| LMOD3 | ENSG00000163380.15 | 5.85E-03 |
| AR | ENSG00000169083.15 | 5.85E-03 |
| SCG2 | ENSG00000171951.4 | 5.86E-03 |
| PIM1 | ENSG00000137193.13 | 5.87E-03 |
| MAPK4 | ENSG00000141639.11 | 5.89E-03 |
| RAB12 | ENSG00000206418.3 | 5.89E-03 |
| TMEM200B | ENSG00000253304.1 | 5.89E-03 |
| LCN6 | ENSG00000267206.5 | 5.94E-03 |
| VGLL3 | ENSG00000206538.7 | 5.99E-03 |
| AL022393.9 | ENSG00000280107.1 | 5.99E-03 |
| BRMS1L | ENSG00000100916.13 | 6.00E-03 |
| RP11-368L12.1 | ENSG00000260658.5 | 6.01E-03 |
| FNDC1 | ENSG00000164694.16 | 6.03E-03 |
| RP11-433J20.1 | ENSG00000236990.1 | 6.03E-03 |
| DDIT4L | ENSG00000145358.6 | 6.03E-03 |
| RP11-490O6.2 | ENSG00000262420.3 | 6.05E-03 |
| ZBTB16 | ENSG00000109906.13 | 6.06E-03 |
| THSD7A | ENSG00000005108.15 | 6.12E-03 |
| LINC00856 | ENSG00000230417.10 | 6.13E-03 |
| TECTA | ENSG00000109927.9 | 6.14E-03 |
| RP11-221N13.3 | ENSG00000256268.1 | 6.19E-03 |
| TAL1 | ENSG00000162367.11 | 6.19E-03 |
| C6 | ENSG00000039537.13 | 6.24E-03 |
| LINC00997 | ENSG00000281332.1 | 6.26E-03 |
| CTD-2033D15.2 | ENSG00000276107.1 | 6.28E-03 |
| LIPM | ENSG00000173239.13 | 6.29E-03 |
| MYOZ3 | ENSG00000164591.13 | 6.33E-03 |
| ISLR2 | ENSG00000167178.15 | 6.35E-03 |
| GFRA3 | ENSG00000146013.10 | 6.35E-03 |
| VIPR2 | ENSG00000106018.13 | 6.35E-03 |
| RECK | ENSG00000122707.11 | 6.36E-03 |
| RP11-286H14.8 | ENSG00000243230.1 | 6.37E-03 |
| SOCS3 | ENSG00000184557.4 | 6.38E-03 |
| ELANE | ENSG00000197561.6 | 6.38E-03 |
| PITX2 | ENSG00000164093.15 | 6.39E-03 |
| AC124944.3 | ENSG00000226155.1 | 6.39E-03 |
| COLEC11 | ENSG00000118004.17 | 6.44E-03 |
| RET | ENSG00000165731.17 | 6.45E-03 |
| RGS5 | ENSG00000143248.12 | 6.45E-03 |
| SNCA | ENSG00000145335.15 | 6.46E-03 |
| IGSF21 | ENSG00000117154.11 | 6.46E-03 |
| DIRC3 | ENSG00000231672.6 | 6.48E-03 |
| RP11-136C24.3 | ENSG00000273291.5 | 6.50E-03 |

Table S2: The genes most associated with disease-free survival (RFS) of patients with stomach adenocarcinoma determined by GEPIA database.

| Gene Symbol | Gene ID | P-Value (Survival dfs) |
| --- | --- | --- |
| LINC01529 | ENSG00000225872.2 | 8.03E-07 |
| ANXA8 | ENSG00000265190.6 | 1.12E-06 |
| GPR87 | ENSG00000138271.5 | 4.82E-06 |
| ALOX12P2 | ENSG00000262943.7 | 8.29E-06 |
| AC009542.2 | ENSG00000231794.5 | 1.05E-05 |
| WNT7A | ENSG00000154764.5 | 1.75E-05 |
| SORCS3 | ENSG00000156395.12 | 2.73E-05 |
| RP3-439F8.1 | ENSG00000234869.1 | 2.75E-05 |
| CTD-2591A1.1 | ENSG00000280159.1 | 2.76E-05 |
| NTRK3 | ENSG00000140538.16 | 2.78E-05 |
| ITIH3 | ENSG00000162267.12 | 2.81E-05 |
| TCEAL5 | ENSG00000204065.2 | 3.15E-05 |
| ATP5F1P5 | ENSG00000254944.1 | 3.17E-05 |
| AC002480.3 | ENSG00000232759.1 | 3.44E-05 |
| SLC35F3 | ENSG00000183780.12 | 3.49E-05 |
| MUSK | ENSG00000030304.12 | 4.55E-05 |
| TEKT2 | ENSG00000092850.11 | 4.56E-05 |
| POU1F1 | ENSG00000064835.10 | 4.99E-05 |
| MYOZ3 | ENSG00000164591.13 | 5.00E-05 |
| CTB-134H23.3 | ENSG00000260908.1 | 5.30E-05 |
| CTD-2114J12.1 | ENSG00000253525.1 | 5.99E-05 |
| C7orf57 | ENSG00000164746.13 | 6.02E-05 |
| RN7SKP296 | ENSG00000223117.1 | 6.03E-05 |
| ABCA6 | ENSG00000154262.12 | 6.19E-05 |
| RP11-430H10.1 | ENSG00000254427.1 | 6.20E-05 |
| RP11-759A24.1 | ENSG00000279645.1 | 6.31E-05 |
| CGB5 | ENSG00000189052.6 | 6.53E-05 |
| NKAIN4 | ENSG00000101198.14 | 6.66E-05 |
| RP11-169F17.1 | ENSG00000263711.5 | 7.73E-05 |
| RP11-401O9.4 | ENSG00000273388.1 | 8.45E-05 |
| ANKRD6 | ENSG00000135299.16 | 8.48E-05 |
| NTF4 | ENSG00000225950.7 | 9.44E-05 |
| BNC1 | ENSG00000169594.12 | 9.75E-05 |
| MAGI2 | ENSG00000187391.17 | 1.10E-04 |
| TCL6 | ENSG00000187621.14 | 1.15E-04 |
| ELANE | ENSG00000197561.6 | 1.18E-04 |
| LRR1 | ENSG00000165501.16 | 1.21E-04 |
| CCDC178 | ENSG00000166960.16 | 1.22E-04 |
| CCNA1 | ENSG00000133101.9 | 1.22E-04 |
| BLK | ENSG00000136573.12 | 1.28E-04 |
| LINC00648 | ENSG00000259129.5 | 1.30E-04 |
| LTBP3 | ENSG00000168056.14 | 1.31E-04 |
| GULP1 | ENSG00000144366.15 | 1.35E-04 |
| COL6A4P1 | ENSG00000230524.8 | 1.37E-04 |
| MAB21L1 | ENSG00000180660.7 | 1.40E-04 |
| TMEM132C | ENSG00000181234.9 | 1.44E-04 |
| POT1-AS1 | ENSG00000224897.6 | 1.45E-04 |
| RP11-613F22.8 | ENSG00000256588.1 | 1.48E-04 |
| PNPT1P1 | ENSG00000229241.1 | 1.49E-04 |
| MIR99AHG | ENSG00000215386.10 | 1.52E-04 |
| COL4A3 | ENSG00000169031.18 | 1.54E-04 |
| TECTA | ENSG00000109927.9 | 1.55E-04 |
| GEMIN8 | ENSG00000046647.13 | 1.57E-04 |
| AC005753.1 | ENSG00000278925.1 | 1.67E-04 |
| RP11-278J6.4 | ENSG00000279130.1 | 1.68E-04 |
| UPK1B | ENSG00000114638.7 | 1.69E-04 |
| EFNA5 | ENSG00000184349.12 | 1.71E-04 |
| DMRTA2 | ENSG00000142700.11 | 1.75E-04 |
| VTN | ENSG00000109072.13 | 1.77E-04 |
| CACNB4 | ENSG00000182389.18 | 1.77E-04 |
| C9orf152 | ENSG00000188959.9 | 1.78E-04 |
| C19orf84 | ENSG00000262874.1 | 1.84E-04 |
| WNT7B | ENSG00000188064.9 | 1.87E-04 |
| CDK15 | ENSG00000138395.14 | 1.92E-04 |
| RP11-449J21.5 | ENSG00000267128.1 | 1.93E-04 |
| TMEM132E | ENSG00000181291.6 | 1.96E-04 |
| RP11-327J17.9 | ENSG00000277135.1 | 1.98E-04 |
| TRIM54 | ENSG00000138100.13 | 1.98E-04 |
| RP11-713P17.3 | ENSG00000204241.7 | 1.99E-04 |
| ACSM6 | ENSG00000173124.14 | 1.99E-04 |
| TOB2P1 | ENSG00000176933.5 | 1.99E-04 |
| AKAP12 | ENSG00000131016.16 | 2.05E-04 |
| TTTY14 | ENSG00000176728.7 | 2.38E-04 |
| PCDHAC1 | ENSG00000248383.4 | 2.42E-04 |
| NPY1R | ENSG00000164128.6 | 2.50E-04 |
| RP11-620J15.3 | ENSG00000257698.1 | 2.55E-04 |
| LINC01152 | ENSG00000256124.5 | 2.56E-04 |
| GFRA3 | ENSG00000146013.10 | 2.64E-04 |
| MRPS2 | ENSG00000122140.10 | 2.72E-04 |
| ACADL | ENSG00000115361.7 | 2.81E-04 |
| NR2F2-AS1 | ENSG00000247809.7 | 2.85E-04 |
| TF | ENSG00000091513.14 | 2.88E-04 |
| RP11-834C11.6 | ENSG00000249388.1 | 2.90E-04 |
| RP11-349H17.2 | ENSG00000275805.1 | 2.93E-04 |
| EBF4 | ENSG00000088881.20 | 2.95E-04 |
| NAA10 | ENSG00000102030.15 | 2.96E-04 |
| TMEM37 | ENSG00000171227.6 | 2.99E-04 |
| CYP8B1 | ENSG00000180432.5 | 3.02E-04 |
| KCNN1 | ENSG00000105642.15 | 3.03E-04 |
| CYP46A1 | ENSG00000036530.8 | 3.03E-04 |
| CCDC152 | ENSG00000198865.9 | 3.04E-04 |
| RP11-59D5__B.2 | ENSG00000236345.1 | 3.04E-04 |
| GPR15 | ENSG00000154165.4 | 3.07E-04 |
| SV2A | ENSG00000159164.9 | 3.09E-04 |
| DAPK1 | ENSG00000196730.12 | 3.16E-04 |
| GCGR | ENSG00000215644.9 | 3.21E-04 |
| APBB1 | ENSG00000166313.18 | 3.26E-04 |
| GRPEL1 | ENSG00000109519.12 | 3.29E-04 |
| LINC00565 | ENSG00000260910.1 | 3.33E-04 |
| CYP24A1 | ENSG00000019186.9 | 3.35E-04 |
| CTD-3157E16.1 | ENSG00000265519.1 | 3.37E-04 |
| PCDHA1 | ENSG00000204970.9 | 3.39E-04 |
| RP4-803A2.1 | ENSG00000217644.5 | 3.41E-04 |
| TUB | ENSG00000166402.8 | 3.46E-04 |
| P2RX2 | ENSG00000187848.12 | 3.49E-04 |
| CTD-3010D24.3 | ENSG00000263893.2 | 3.50E-04 |
| GRID1 | ENSG00000182771.17 | 3.50E-04 |
| AC007319.1 | ENSG00000224063.5 | 3.55E-04 |
| MPRIP-AS1 | ENSG00000225442.2 | 3.55E-04 |
| LINC01426 | ENSG00000234380.1 | 3.59E-04 |
| DKK1 | ENSG00000107984.9 | 3.80E-04 |
| C10orf10 | ENSG00000165507.8 | 3.80E-04 |
| RP6-74O6.6 | ENSG00000272824.1 | 3.82E-04 |
| PACS1 | ENSG00000175115.11 | 3.92E-04 |
| RP4-806M20.4 | ENSG00000268649.3 | 3.95E-04 |
| RP11-482D24.3 | ENSG00000257918.1 | 3.95E-04 |
| GLIS2 | ENSG00000126603.8 | 4.05E-04 |
| PCDH15 | ENSG00000150275.17 | 4.07E-04 |
| MRPL35 | ENSG00000132313.14 | 4.07E-04 |
| FLOT2 | ENSG00000132589.15 | 4.16E-04 |
| STK32A | ENSG00000169302.14 | 4.19E-04 |
| AC006460.2 | ENSG00000228509.5 | 4.19E-04 |
| CYP11A1 | ENSG00000140459.17 | 4.23E-04 |
| PCOLCE2 | ENSG00000163710.7 | 4.23E-04 |
| CTD-2015G9.2 | ENSG00000261175.5 | 4.31E-04 |
| RP11-60A14.1 | ENSG00000279778.1 | 4.42E-04 |
| ARRDC3 | ENSG00000113369.8 | 4.46E-04 |
| ARHGAP31-AS1 | ENSG00000241155.1 | 4.53E-04 |
| LINC00652 | ENSG00000179935.9 | 4.54E-04 |
| RP11-433P17.3 | ENSG00000262118.1 | 4.58E-04 |
| CTD-2311M21.3 | ENSG00000261821.2 | 4.61E-04 |
| RP11-181K12.1 | ENSG00000279481.1 | 4.61E-04 |
| RP11-44D5.1 | ENSG00000270409.1 | 4.62E-04 |
| CLDN6 | ENSG00000184697.6 | 4.68E-04 |
| TRHDE-AS1 | ENSG00000236333.3 | 4.68E-04 |
| AP000662.4 | ENSG00000254602.1 | 4.79E-04 |
| RP11-63M22.1 | ENSG00000260558.1 | 4.89E-04 |
| KCNS1 | ENSG00000124134.8 | 4.91E-04 |
| NOVA1 | ENSG00000139910.19 | 4.91E-04 |
| CLDN11 | ENSG00000013297.10 | 4.94E-04 |
| TCEAL7 | ENSG00000182916.7 | 4.98E-04 |
| CSMD1 | ENSG00000183117.17 | 4.99E-04 |
| TGM5 | ENSG00000104055.14 | 5.03E-04 |
| CKS2 | ENSG00000123975.4 | 5.08E-04 |
| RP11-981P6.1 | ENSG00000258302.2 | 5.13E-04 |
| TRPC1 | ENSG00000144935.14 | 5.20E-04 |
| PCDHB4 | ENSG00000081818.3 | 5.31E-04 |
| CH17-360D5.2 | ENSG00000276850.4 | 5.36E-04 |
| AIFM1 | ENSG00000156709.13 | 5.41E-04 |
| LINC00506 | ENSG00000281392.1 | 5.41E-04 |
| ABCA8 | ENSG00000141338.13 | 5.48E-04 |
| RP11-435O5.2 | ENSG00000237857.2 | 5.52E-04 |
| CTB-134H23.2 | ENSG00000196796.5 | 5.58E-04 |
| CFHR1 | ENSG00000244414.6 | 5.59E-04 |
| C9orf40 | ENSG00000135045.6 | 5.66E-04 |
| WNT1 | ENSG00000125084.11 | 5.68E-04 |
| SLC30A8 | ENSG00000164756.12 | 5.73E-04 |
| RARB | ENSG00000077092.18 | 5.81E-04 |
| FAM174B | ENSG00000185442.12 | 5.84E-04 |
| CPEB1 | ENSG00000214575.9 | 5.98E-04 |
| RP11-102K13.5 | ENSG00000278309.1 | 6.03E-04 |
| ST6GAL2 | ENSG00000144057.15 | 6.20E-04 |
| TPST1 | ENSG00000169902.13 | 6.21E-04 |
| IRX6 | ENSG00000159387.7 | 6.28E-04 |
| LINC00346 | ENSG00000255874.2 | 6.46E-04 |
| FLJ16779 | ENSG00000275620.1 | 6.51E-04 |
| RPL23AP58 | ENSG00000228657.1 | 6.53E-04 |
| AP001626.2 | ENSG00000235023.1 | 6.57E-04 |
| KCNT2 | ENSG00000162687.16 | 6.57E-04 |
| KLHL38 | ENSG00000175946.8 | 6.75E-04 |
| AR | ENSG00000169083.15 | 6.83E-04 |
| THSD4 | ENSG00000187720.14 | 6.87E-04 |
| FGF14 | ENSG00000102466.15 | 6.95E-04 |
| SCN5A | ENSG00000183873.15 | 7.09E-04 |
| RIMS4 | ENSG00000101098.12 | 7.10E-04 |
| PCDHB5 | ENSG00000113209.8 | 7.17E-04 |
| ARMCX4 | ENSG00000196440.11 | 7.17E-04 |
| C17orf50 | ENSG00000270806.1 | 7.25E-04 |
| PTPRQ | ENSG00000139304.12 | 7.32E-04 |
| CAPG | ENSG00000042493.15 | 7.33E-04 |
| PRTG | ENSG00000166450.12 | 7.35E-04 |
| SLCO4A1 | ENSG00000101187.15 | 7.40E-04 |
| UBXN7-AS1 | ENSG00000225822.4 | 7.42E-04 |
| RP11-209D14.4 | ENSG00000266126.1 | 7.49E-04 |
| C12orf77 | ENSG00000226397.7 | 7.53E-04 |
| HERC2P5 | ENSG00000260644.6 | 7.55E-04 |
| RP11-445P17.8 | ENSG00000224034.1 | 7.61E-04 |
| IGHD | ENSG00000211898.7 | 7.66E-04 |
| RP11-1260E13.4 | ENSG00000262061.5 | 7.72E-04 |
| ZDHHC2 | ENSG00000104219.12 | 7.73E-04 |
| NDUFA8 | ENSG00000119421.6 | 7.80E-04 |
| RP11-1191J2.5 | ENSG00000272927.1 | 7.92E-04 |
| WDR91 | ENSG00000105875.13 | 7.94E-04 |
| LINC00540 | ENSG00000276476.2 | 8.03E-04 |
| LURAP1 | ENSG00000171357.5 | 8.13E-04 |
| RP11-94C24.13 | ENSG00000275897.1 | 8.36E-04 |
| PALM | ENSG00000099864.17 | 8.37E-04 |
| FBXO27 | ENSG00000161243.8 | 8.42E-04 |
| RP11-80I15.1 | ENSG00000223849.1 | 8.51E-04 |
| RP11-697N18.1 | ENSG00000251354.3 | 8.71E-04 |
| SLC4A5 | ENSG00000188687.15 | 8.73E-04 |
| RP11-80A15.1 | ENSG00000258744.1 | 8.75E-04 |
| RP11-474C8.8 | ENSG00000274124.1 | 8.82E-04 |
| RP11-497E19.1 | ENSG00000205562.2 | 8.85E-04 |
| AC004538.3 | ENSG00000230333.6 | 8.92E-04 |
| DYNLRB2 | ENSG00000168589.14 | 9.21E-04 |
| RP1-29C18.8 | ENSG00000235111.1 | 9.27E-04 |
| CTD-2555C10.3 | ENSG00000259230.1 | 9.27E-04 |
| CTC-487M23.7 | ENSG00000272389.1 | 9.28E-04 |
| SMURF1 | ENSG00000198742.9 | 9.35E-04 |
| SH3GL3 | ENSG00000140600.16 | 9.35E-04 |
| FAM216B | ENSG00000179813.6 | 9.45E-04 |
| GJA1P1 | ENSG00000176857.5 | 9.46E-04 |
| SIGLEC6 | ENSG00000105492.15 | 9.47E-04 |
| HSPB1P2 | ENSG00000230216.1 | 9.48E-04 |
| PLCXD3 | ENSG00000182836.9 | 9.51E-04 |
| TTC21B-AS1 | ENSG00000224490.5 | 9.53E-04 |
| LSAMP-AS1 | ENSG00000240922.1 | 9.56E-04 |
| ITIH4 | ENSG00000055955.15 | 9.67E-04 |
| YPEL4 | ENSG00000166793.10 | 9.70E-04 |
| CLEC4F | ENSG00000152672.7 | 9.77E-04 |
| CALCR | ENSG00000004948.13 | 9.78E-04 |
| RP5-1139B12.2 | ENSG00000269890.1 | 9.80E-04 |
| RP11-823P9.4 | ENSG00000279107.1 | 9.87E-04 |
| AL591893.1 | ENSG00000229021.2 | 9.90E-04 |
| PCDHA13 | ENSG00000239389.7 | 9.93E-04 |
| SCARA5 | ENSG00000168079.16 | 9.94E-04 |
| VWA5B1 | ENSG00000158816.15 | 9.94E-04 |
| RP11-890B15.2 | ENSG00000254842.6 | 9.95E-04 |
| PCDHAC2 | ENSG00000243232.4 | 1.01E-03 |
| TP53TG3D | ENSG00000205456.11 | 1.01E-03 |
| RP11-522B15.3 | ENSG00000259275.2 | 1.04E-03 |
| RP11-161I6.2 | ENSG00000263745.5 | 1.04E-03 |
| RFPL1S | ENSG00000225465.8 | 1.04E-03 |
| C1QTNF4 | ENSG00000172247.3 | 1.05E-03 |
| ANKRD53 | ENSG00000144031.11 | 1.05E-03 |
| CTSG | ENSG00000100448.3 | 1.06E-03 |
| RP11-399B17.1 | ENSG00000278962.1 | 1.06E-03 |
| RP11-353N14.2 | ENSG00000262772.1 | 1.08E-03 |
| AC019117.2 | ENSG00000236039.1 | 1.08E-03 |
| PCDHB12 | ENSG00000120328.6 | 1.08E-03 |
| C9orf41 | ENSG00000156017.12 | 1.08E-03 |
| RAB40A | ENSG00000172476.3 | 1.10E-03 |
| RP11-483C6.1 | ENSG00000262119.1 | 1.10E-03 |
| RTL1 | ENSG00000254656.1 | 1.11E-03 |
| PCNA | ENSG00000132646.10 | 1.11E-03 |
| SLC45A1 | ENSG00000162426.14 | 1.11E-03 |
| RP5-1185I7.1 | ENSG00000232756.1 | 1.11E-03 |
| HNRNPA1P12 | ENSG00000220157.4 | 1.12E-03 |
| NALCN | ENSG00000102452.15 | 1.13E-03 |
| LINC01260 | ENSG00000132832.9 | 1.13E-03 |
| FAM198A | ENSG00000144649.8 | 1.14E-03 |
| RP11-306O13.1 | ENSG00000213121.2 | 1.15E-03 |
| RPP25L | ENSG00000164967.9 | 1.15E-03 |
| RP11-307C18.1 | ENSG00000272950.1 | 1.17E-03 |
| SYPL2 | ENSG00000143028.8 | 1.18E-03 |
| LRAT | ENSG00000121207.11 | 1.18E-03 |
| RP11-81H3.2 | ENSG00000251138.6 | 1.18E-03 |
| HAVCR1 | ENSG00000113249.12 | 1.18E-03 |
| LRRTM1 | ENSG00000162951.10 | 1.19E-03 |
| LRFN5 | ENSG00000165379.13 | 1.19E-03 |
| RP11-542B15.1 | ENSG00000203585.3 | 1.20E-03 |
| LINC00473 | ENSG00000223414.2 | 1.22E-03 |
| RP1-302G2.5 | ENSG00000262179.2 | 1.24E-03 |
| FAM153B | ENSG00000182230.11 | 1.24E-03 |
| ONECUT1 | ENSG00000169856.8 | 1.25E-03 |
| RBP4 | ENSG00000138207.12 | 1.25E-03 |
| RP11-209M4.1 | ENSG00000267253.1 | 1.25E-03 |
| EGOT | ENSG00000235947.1 | 1.26E-03 |
| L3MBTL3 | ENSG00000198945.7 | 1.27E-03 |
| MMACHC | ENSG00000132763.14 | 1.27E-03 |
| DRC7 | ENSG00000159625.14 | 1.27E-03 |
| NPTX1 | ENSG00000171246.5 | 1.28E-03 |
| WTAPP1 | ENSG00000255282.6 | 1.28E-03 |
| GRIK4 | ENSG00000149403.11 | 1.30E-03 |
| GNAS-AS1 | ENSG00000235590.7 | 1.31E-03 |
| AC004813.1 | ENSG00000279777.1 | 1.32E-03 |
| TCP11 | ENSG00000124678.17 | 1.32E-03 |
| RP11-806O11.1 | ENSG00000253671.1 | 1.32E-03 |
| OR5K2 | ENSG00000231861.2 | 1.33E-03 |
| RP11-631M6.3 | ENSG00000251682.1 | 1.34E-03 |
| RNF183 | ENSG00000165188.13 | 1.35E-03 |
| RERG | ENSG00000134533.6 | 1.38E-03 |
| SH3GL2 | ENSG00000107295.9 | 1.40E-03 |
| CLRN3 | ENSG00000180745.4 | 1.40E-03 |
| DCLK1 | ENSG00000133083.14 | 1.41E-03 |
| ZNF667 | ENSG00000198046.11 | 1.41E-03 |
| CTD-2215E18.1 | ENSG00000251513.2 | 1.42E-03 |
| SCTR | ENSG00000080293.9 | 1.43E-03 |
| PRSS50 | ENSG00000206549.12 | 1.44E-03 |
| AP001626.1 | ENSG00000225431.1 | 1.44E-03 |
| KPNA7 | ENSG00000185467.7 | 1.45E-03 |
| RP11-712P20.2 | ENSG00000266965.1 | 1.46E-03 |
| ZNF474 | ENSG00000164185.4 | 1.46E-03 |
| ENPP1 | ENSG00000197594.11 | 1.46E-03 |
| RP11-48B3.3 | ENSG00000254162.1 | 1.47E-03 |
| RP11-295M18.6 | ENSG00000272823.1 | 1.47E-03 |
| CTD-2536I1.2 | ENSG00000280304.1 | 1.47E-03 |
| KIRREL-IT1 | ENSG00000226520.1 | 1.48E-03 |
| FLJ45079 | ENSG00000204283.3 | 1.48E-03 |
| FAM153C | ENSG00000204677.10 | 1.48E-03 |
| CNTN2 | ENSG00000184144.9 | 1.50E-03 |
| AC013275.2 | ENSG00000231013.1 | 1.52E-03 |
| SLC15A2 | ENSG00000163406.10 | 1.53E-03 |
| UNC13C | ENSG00000137766.16 | 1.54E-03 |
| PCDHA3 | ENSG00000255408.3 | 1.54E-03 |
| FGF19 | ENSG00000162344.3 | 1.57E-03 |
| SLC5A10 | ENSG00000154025.15 | 1.59E-03 |
| CD300LG | ENSG00000161649.12 | 1.59E-03 |
| RP11-454L9.2 | ENSG00000259318.1 | 1.59E-03 |
| RPL21P40 | ENSG00000235670.1 | 1.60E-03 |
| TBC1D27 | ENSG00000128438.10 | 1.61E-03 |
| AC092162.1 | ENSG00000230552.5 | 1.61E-03 |
| RP11-284F21.10 | ENSG00000272405.1 | 1.62E-03 |
| ASCL1 | ENSG00000139352.3 | 1.62E-03 |
| SEC14L3 | ENSG00000100012.11 | 1.64E-03 |
| KLB | ENSG00000134962.6 | 1.64E-03 |
| BACH1-AS1 | ENSG00000232118.2 | 1.64E-03 |
| LA16c-3G11.7 | ENSG00000241838.3 | 1.65E-03 |
| RP11-474O21.5 | ENSG00000272482.1 | 1.65E-03 |
| RP11-438L7.3 | ENSG00000255967.1 | 1.67E-03 |
| KIF5A | ENSG00000155980.11 | 1.67E-03 |
| ZSCAN10 | ENSG00000130182.7 | 1.70E-03 |
| FKBP4P6 | ENSG00000268234.1 | 1.71E-03 |
| RP11-616M22.11 | ENSG00000273551.1 | 1.71E-03 |
| AC011997.1 | ENSG00000222017.1 | 1.72E-03 |
| RP11-789C17.1 | ENSG00000265413.1 | 1.72E-03 |
| BEST3 | ENSG00000127325.18 | 1.74E-03 |
| SUMO4 | ENSG00000177688.6 | 1.76E-03 |
| RP11-37N22.1 | ENSG00000214803.3 | 1.78E-03 |
| ARAP1-AS2 | ENSG00000245148.2 | 1.78E-03 |
| RP11-462L8.1 | ENSG00000229656.6 | 1.79E-03 |
| RP11-586K12.1 | ENSG00000279795.1 | 1.79E-03 |
| CYP27C1 | ENSG00000186684.12 | 1.80E-03 |
| RAB34 | ENSG00000109113.17 | 1.80E-03 |
| MNS1 | ENSG00000138587.5 | 1.81E-03 |
| KRT79 | ENSG00000185640.5 | 1.81E-03 |
| PI3 | ENSG00000124102.4 | 1.82E-03 |
| FAM193B | ENSG00000146067.15 | 1.82E-03 |
| LRTM2 | ENSG00000166159.10 | 1.83E-03 |
| CDC42P5 | ENSG00000253439.1 | 1.83E-03 |
| CFAP221 | ENSG00000163075.12 | 1.84E-03 |
| RP11-266K4.14 | ENSG00000275367.1 | 1.85E-03 |
| RP4-565E6.1 | ENSG00000227733.8 | 1.85E-03 |
| SVEP1 | ENSG00000165124.17 | 1.85E-03 |
| SYN1 | ENSG00000008056.12 | 1.86E-03 |
| XXyac-YX155B6.5 | ENSG00000232265.7 | 1.88E-03 |
| OGN | ENSG00000106809.10 | 1.90E-03 |
| ITFG2 | ENSG00000111203.11 | 1.90E-03 |
| GALNT16 | ENSG00000100626.16 | 1.92E-03 |
| SUSD5 | ENSG00000173705.8 | 1.92E-03 |
| CTD-2195B23.3 | ENSG00000269652.1 | 1.92E-03 |
| ZNF367 | ENSG00000165244.6 | 1.93E-03 |
| PP2D1 | ENSG00000183977.13 | 1.93E-03 |
| RPL7AP65 | ENSG00000228000.1 | 1.94E-03 |
| RNF219-AS1 | ENSG00000234377.7 | 1.94E-03 |
| NIPAL4 | ENSG00000172548.14 | 1.94E-03 |
| TVP23A | ENSG00000166676.14 | 1.96E-03 |
| HNRNPA1P8 | ENSG00000229251.3 | 1.97E-03 |
| CTD-2530N21.4 | ENSG00000254064.1 | 1.98E-03 |
| HCAR1 | ENSG00000196917.5 | 1.98E-03 |
| INHBA-AS1 | ENSG00000224116.6 | 1.98E-03 |
| RP11-266K4.9 | ENSG00000215241.3 | 1.99E-03 |
| A2M-AS1 | ENSG00000245105.2 | 2.02E-03 |
| NUDT2 | ENSG00000164978.17 | 2.02E-03 |
| NIM1K | ENSG00000177453.7 | 2.02E-03 |
| RNF165 | ENSG00000141622.13 | 2.02E-03 |
| HCG4B | ENSG00000227262.3 | 2.02E-03 |
| EFCAB12 | ENSG00000172771.11 | 2.03E-03 |
| AC108676.1 | ENSG00000244675.2 | 2.03E-03 |
| RN7SL683P | ENSG00000242330.3 | 2.03E-03 |
| SPIRE1 | ENSG00000134278.14 | 2.04E-03 |
| NTNG1 | ENSG00000162631.18 | 2.04E-03 |
| JAZF1-AS1 | ENSG00000234336.6 | 2.06E-03 |
| FGF14-AS2 | ENSG00000272143.1 | 2.06E-03 |
| RP11-23D24.2 | ENSG00000238755.3 | 2.07E-03 |
| NBAT1 | ENSG00000260455.1 | 2.08E-03 |
| SP7 | ENSG00000170374.5 | 2.08E-03 |
| CPNE8 | ENSG00000139117.13 | 2.08E-03 |
| AC093642.1 | ENSG00000280119.1 | 2.08E-03 |
| SERPINA4 | ENSG00000100665.11 | 2.08E-03 |
| LINC00202-2 | ENSG00000231976.7 | 2.11E-03 |
| NPAS3 | ENSG00000151322.18 | 2.12E-03 |
| MAB21L2 | ENSG00000181541.5 | 2.13E-03 |
| WNT10B | ENSG00000169884.13 | 2.13E-03 |
| RP11-303E16.6 | ENSG00000261838.5 | 2.13E-03 |
| RPL10P1 | ENSG00000217026.3 | 2.15E-03 |
| TBX6 | ENSG00000149922.10 | 2.17E-03 |
| CPT1C | ENSG00000169169.14 | 2.18E-03 |
| KIRREL2 | ENSG00000126259.19 | 2.18E-03 |
| PZP | ENSG00000126838.9 | 2.19E-03 |
| FOXE1 | ENSG00000178919.8 | 2.19E-03 |
| OCA2 | ENSG00000104044.15 | 2.21E-03 |
| RPL23AP23 | ENSG00000236863.2 | 2.21E-03 |
| KIF21B | ENSG00000116852.14 | 2.22E-03 |
| DMRTC1B | ENSG00000184911.14 | 2.22E-03 |
| SCRN1 | ENSG00000136193.16 | 2.23E-03 |
| OPN3 | ENSG00000054277.12 | 2.23E-03 |
| USE1 | ENSG00000053501.12 | 2.23E-03 |
| NBPF14 | ENSG00000270629.5 | 2.23E-03 |
| PPP4R1L | ENSG00000124224.16 | 2.26E-03 |
| NUDT7 | ENSG00000140876.11 | 2.28E-03 |
| IGSF10 | ENSG00000152580.8 | 2.29E-03 |
| RP11-110I1.14 | ENSG00000271751.1 | 2.30E-03 |
| RP11-382B18.1 | ENSG00000279417.1 | 2.30E-03 |
| TSNAXIP1 | ENSG00000102904.14 | 2.31E-03 |
| RP11-243M5.2 | ENSG00000280200.1 | 2.31E-03 |
| ONECUT2 | ENSG00000119547.5 | 2.32E-03 |
| UG0898H09 | ENSG00000274956.2 | 2.33E-03 |
| RP11-463O9.9 | ENSG00000270020.1 | 2.33E-03 |
| TSPY26P | ENSG00000235217.6 | 2.34E-03 |
| ZBTB20-AS1 | ENSG00000241560.5 | 2.34E-03 |
| ZFPM2 | ENSG00000169946.13 | 2.35E-03 |
| LINC01176 | ENSG00000281404.1 | 2.35E-03 |
| TBC1D3B | ENSG00000274808.4 | 2.36E-03 |
| RP11-867G23.10 | ENSG00000254510.1 | 2.36E-03 |
| KLHL14 | ENSG00000197705.9 | 2.36E-03 |
| RP11-470L19.2 | ENSG00000235407.1 | 2.36E-03 |
| TET1 | ENSG00000138336.8 | 2.38E-03 |
| MAN2B1 | ENSG00000104774.12 | 2.38E-03 |
| MAP2 | ENSG00000078018.19 | 2.39E-03 |
| RP4-724E13.2 | ENSG00000228204.2 | 2.40E-03 |
| CLTA | ENSG00000122705.16 | 2.40E-03 |
| CTC-480C2.1 | ENSG00000250874.1 | 2.40E-03 |
| CFH | ENSG00000000971.15 | 2.41E-03 |
| RP11-426C22.4 | ENSG00000259807.1 | 2.41E-03 |
| GBA2 | ENSG00000070610.14 | 2.41E-03 |
| PTPN5 | ENSG00000110786.17 | 2.41E-03 |
| POLR3DP1 | ENSG00000214626.2 | 2.42E-03 |
| CTF1 | ENSG00000150281.6 | 2.42E-03 |
| SIGLEC17P | ENSG00000171101.13 | 2.43E-03 |
| STXBP6 | ENSG00000168952.15 | 2.43E-03 |
| GDF6 | ENSG00000156466.9 | 2.43E-03 |
| CCNDBP1 | ENSG00000166946.13 | 2.43E-03 |
| TNNT3 | ENSG00000130595.16 | 2.43E-03 |
| RP11-123C21.2 | ENSG00000270986.1 | 2.44E-03 |
| CTD-3035K23.3 | ENSG00000279713.1 | 2.45E-03 |
| Mar-10 | ENSG00000173838.11 | 2.46E-03 |
| COMMD10 | ENSG00000145781.8 | 2.47E-03 |
| ELAVL3 | ENSG00000196361.9 | 2.47E-03 |
| CDC20B | ENSG00000164287.12 | 2.47E-03 |
| NDRG4 | ENSG00000103034.14 | 2.48E-03 |
| TMEM240 | ENSG00000205090.8 | 2.49E-03 |
| SDCCAG3P2 | ENSG00000181101.7 | 2.49E-03 |
| CCDC181 | ENSG00000117477.12 | 2.49E-03 |
| SERPINA5 | ENSG00000188488.13 | 2.49E-03 |
| TIGD6 | ENSG00000164296.6 | 2.49E-03 |
| XXbac-BPG154L12.4 | ENSG00000225914.1 | 2.49E-03 |
| FTO-IT1 | ENSG00000260936.1 | 2.50E-03 |
| RP11-73M18.6 | ENSG00000270108.1 | 2.52E-03 |
| ZMAT2 | ENSG00000146007.10 | 2.54E-03 |
| ABCA9-AS1 | ENSG00000231749.3 | 2.55E-03 |
| ABHD8 | ENSG00000127220.5 | 2.56E-03 |
| RP4-665N4.4 | ENSG00000232862.5 | 2.56E-03 |
| ISCA1 | ENSG00000135070.13 | 2.56E-03 |
| RPL5P17 | ENSG00000243859.3 | 2.57E-03 |
| RP11-297J22.1 | ENSG00000271709.1 | 2.58E-03 |
| LOXL4 | ENSG00000138131.3 | 2.59E-03 |
| RP11-544L8__B.4 | ENSG00000175967.3 | 2.60E-03 |
| GS1-120K12.4 | ENSG00000260976.1 | 2.62E-03 |
| RP11-1000B6.7 | ENSG00000276724.1 | 2.63E-03 |
| KCNS2 | ENSG00000156486.7 | 2.63E-03 |
| BIRC7 | ENSG00000101197.12 | 2.65E-03 |
| LRRC3B | ENSG00000179796.11 | 2.65E-03 |
| NOX5 | ENSG00000255346.9 | 2.66E-03 |
| TSPYL2 | ENSG00000184205.14 | 2.67E-03 |
| CTB-33G10.1 | ENSG00000243829.1 | 2.67E-03 |
| FAM231D | ENSG00000272058.2 | 2.68E-03 |
| TTTY15 | ENSG00000233864.7 | 2.69E-03 |
| CASC10 | ENSG00000204682.5 | 2.70E-03 |
| ASF1B | ENSG00000105011.8 | 2.70E-03 |
| HMGB1P23 | ENSG00000253770.1 | 2.71E-03 |
| RP11-894J14.2 | ENSG00000279144.1 | 2.71E-03 |
| SLC18A2 | ENSG00000165646.11 | 2.72E-03 |
| ZNF677 | ENSG00000197928.10 | 2.72E-03 |
| RP11-64B16.2 | ENSG00000213144.2 | 2.73E-03 |
| PNMA1 | ENSG00000176903.4 | 2.74E-03 |
| CTD-2509G16.2 | ENSG00000255002.1 | 2.74E-03 |
| ZNF660 | ENSG00000144792.9 | 2.74E-03 |
| PCDHB18P | ENSG00000146001.5 | 2.74E-03 |
| APOA1-AS | ENSG00000235910.1 | 2.74E-03 |
| ZNF790-AS1 | ENSG00000267254.5 | 2.75E-03 |
| MIR181A1HG | ENSG00000229989.3 | 2.75E-03 |
| BCAS3 | ENSG00000141376.20 | 2.75E-03 |
| FAT2 | ENSG00000086570.12 | 2.77E-03 |
| TUBA4B | ENSG00000243910.7 | 2.78E-03 |
| PRKG1-AS1 | ENSG00000236671.7 | 2.78E-03 |
| RP11-148O21.2 | ENSG00000255354.1 | 2.78E-03 |
| CASC18 | ENSG00000257859.1 | 2.79E-03 |
| CDHR4 | ENSG00000187492.8 | 2.79E-03 |
| RPS15AP38 | ENSG00000237668.1 | 2.81E-03 |
| CASP5 | ENSG00000137757.10 | 2.81E-03 |
| ERBB4 | ENSG00000178568.13 | 2.83E-03 |
| UNC13D | ENSG00000092929.11 | 2.83E-03 |
| RP11-936I5.1 | ENSG00000266998.1 | 2.84E-03 |
| BCL2L10 | ENSG00000137875.4 | 2.84E-03 |
| MGAT5B | ENSG00000167889.12 | 2.85E-03 |
| DTX1 | ENSG00000135144.7 | 2.85E-03 |
| NECAP1P2 | ENSG00000234632.1 | 2.86E-03 |
| RP11-615I2.2 | ENSG00000260577.2 | 2.87E-03 |
